# The Influence of Early Life Stress on the Development of Neural Representations during Mentalizing

**DOI:** 10.64898/2026.09.18.752602

**Authors:** Miro Ilomäki, Jallu Lindblom, Marjo Flykt, Mervi Vänskä, Patrik Wikman

## Abstract

Early life stress (ELS) is associated with altered social cognition, but the neurodevelopmental mechanisms of this association remain understudied. In the present longitudinal study spanning 20-years, eighty-nine young adults performed the Reading the Mind in the Eyes Test (RMET) with a matched control condition during fMRI. Two ELS indices were inspected separately: a prospective cumulative risk score based on caregiver wellbeing during pregnancy and first year of life, and a retrospective self-report of adverse experiences by the participants in late adolescence. Neural outcomes were characterized by univariate activation, representational similarity analysis (RSA), and inter-subject RSA. Inter-subject RSA tested two pairwise approaches to intersubject similarity in ELS: absolute pairwise difference, and pairwise average. Prospective ELS was associated with better RMET performance, and one region survived correction across 360 cortical regions tested in RSA and IS-RSA: left anterior inferior frontal sulcus. The IS-RSA effect was pairwise average: low-exposure participants shared a more typical neural representational geometry, while high-exposure participants demonstrated increasing neural representational idiosyncrasy. Additionally, higher prospective ELS was associated with weaker RMET-versus-control separation within the same region. Retrospective ELS yielded no behavioral nor neural associations. The present study offers novel findings relating to neural representations of mentalizing using the RMET, alongside their neurodevelopmental susceptibility to ELS. The present study additionally highlights implications relating to the differentiation of mentalizing from lexical-semantic and semantic-control demands for future research.

## Introduction

Humans possess, to varying degrees, the capacity to “hold mind in mind” (Allen et al., 2008), that is, to internally represent the mental states of others and oneself. This so-called mentalizing capacity is foundational for cooperation, communication, and other means of navigating social relationships and interactions (Arabadzhiev & Paunova, 2024; Fonagy & Allison, 2012). Crucially, mentalizing develops through the interplay between biological maturation and experiences shaped by the social environment (Kim, 2015; Hughes et al., 2005; Gergely & Watson, 1999). However, children’s developmental sensitivity to early social experiences also predisposes them to vulnerability: early life stress (ELS) can shape the developmental trajectory of social cognition and the neural architecture underlying it. It is therefore necessary to ask not only which neural systems support the representation of others’ mental states and how such information is encoded and organized in the brain, but also whether and how these representations vary as a function of ELS.

At the conceptual level, mental state attribution has been denoted with several partly overlapping concepts besides mentalizing, such as theory of mind, mindreading, social cognition, and perspective-taking. While a perfect expert consensus on the precise use of this heterogenous conceptual landscape is yet to be reached, agreement on the proper definitions of these concepts has been investigated (Quesque et al., 2024). For example, Theory of Mind (ToM), coined by Premack and Woodruff (1978), is defined as the use of folk psychological knowledge and heuristics to think about one’s own and other people’s mental states (Quesque et al., 2024). Mentalizing (Fonagy, 1991), on the other hand, is defined as the ability to attribute mental states (e.g., knowledge, intentions, emotions, perception) to self and others (Quesque et al., 2024; Fonagy & Allison, 2012). While conceptual heterogeneity still exists within and between different fields of research that study and theorize about social cognitive processes relating to mental state attribution, in the present study, in line with expert consensus, we will use the concept ‘mentalizing’.

In the present study, mentalizing was evaluated using the Reading the Mind in the Eyes Task (RMET; Baron-Cohen et al., 2001), which requires participants to infer mental states from images of the eye region of different people. Empirically, RMET is the most often employed task for evaluating mentalizing in neurotypical adults (Yeung et al., 2024). Additionally, RMET has also been consistently utilized in neuroimaging research regarding neural correlates of mentalizing (Schurz et al., 2021; Schurtz et al, 2014). At the neural level, converging evidence from studies utilizing the RMET, alongside other mentalizing tasks, implicates a distributed network in mentalizing and related social-cognitive processes. Meta-analytical work has established the following bilateral regions as consistently belonging to this network: superior temporal sulcus (STS), middle temporal gyrus (MTG), temporoparietal junction (TPJ), medial prefrontal cortex (mPFC), inferior frontal gyrus (IFG), and precuneus (Maliske et al., 2023; Schurz et al., 2021; Molenberghs et al., 2016; Schurz et al., 2014; Mar, 2011; Frith & Frith, 2006). Despite the converging evidence, research specifically focused on the neural correlates of mentalizing in neurotypical and non-clinical (i.e. without psychiatric diagnoses) adult samples remains sparse.

Whether indexed by adverse or traumatic childhood experiences, or by moderate but chronic stress exposures, ELS has been associated with diminished or disrupted mentalizing and social cognitive ability (Gorgellino et al., 2025; Mackey et al., 2025; Yang & Huang, 2024; Martin-Gagnon et al., 2023; Wagner-Skacel et al., 2022; Rokita et al., 2018; Pechtel & Pizzagalli, 2011). However, prospective and longitudinal research investigating the influence of various forms of ELS on mentalizing capacity in the non-clinical population is lacking, and most of this previous research focuses on relatively extreme ELS events, such as abuse and maltreatment by caregivers. A recent review of the literature (Mackey et al., 2025) suggests that retrospective accounts of ELS are associated with diminished mentalizing capacities even among adults with no psychiatric diagnoses. Yet the same authors highlight that subgroup, sensitivity, and publication bias analyses were precluded due to the limited amount of previous research. Current prospective research on the influence of more moderate forms of ELS on mentalizing capacities in the non-clinical population is thus lacking.

Few neurodevelopmental studies (e.g., Trujillo-Llano et al., 2024; Cracco et al., 2020; Vai et al., 2017; Nolte et al., 2013) have been devoted towards unravelling the influence of ELS on the neural processes underlying mentalizing. Previous neurodevelopmental research is overwhelmingly univariate, focusing on task-based activation or resting-state functional connectivity instead of task-based multivariate neural activation patterns. Multivariate approaches, such as representational similarity analysis (RSA; Kriegeskorte, 2008), characterize the information carried by distributed activity patterns. Although such methods are well established and have been applied in general research on mentalizing and social cognition (Buergi et al., 2026; Golec-Staśkiewicz et al., 2022; Freeman et al., 2018; Thornton et al., 2019; Thornton & Mitchell, 2018; Koster-Hale et al., 2017), almost no attention has been devoted to investigating multivariate neural representations of mentalizing in ELS related developmental contexts.

In the present whole-brain fMRI study, we use a 20-year spanning longitudinal, non-clinical sample of Finnish families followed from pregnancy to the children’s early adulthood, while utilizing two qualitatively different measures of ELS and a multivariate neural representational paradigm. Using the RMET, we investigate whether ELS is associated with individual differences in the neural representation of mentalizing. To do so, we employ univariate activation analyses, representational similarity analyses, and inter-subject representational similarity analyses. This approach allows us to distinguish between three related questions: whether ELS is associated with differences in overall neural activation during mentalizing, whether it is associated with differences in the organization of local activity patterns during mentalizing, and whether those with similar ELS exposure show similar neural representational structure. By integrating these levels of analysis, we aim to characterize both the anatomical distribution and representational organization of mentalizing-related neural processing, and what developmental influence ELS might have on them.

## Methods

### Participants

The participants in the present study comprise a subsample of a larger Miracles of Development (MIDE) research project sample of 953 Finnish families followed from pregnancy onward. Half of the children were conceived with assisted reproductive technology (ART; n = 484, 51%) and the remainder naturally (NC; n = 469, 49%). At recruitment during pregnancy, parents had to be Finnish-speaking; for the NC group, additional criteria were no history of infertility and maternal age > 25 years to align with the higher age of ART mothers. For fuller cohort descriptions, see Vänskä et al. (2011) and Flykt et al. (2021).

To obtain an fMRI subsample that reflected the full range of early life stress (ELS), we used stratified sampling with disproportionate allocation (Parsons, 2017) based on a Prospective ELS index. This index comprised 20 indicators of maternal/paternal mental health and family-relationship difficulties assessed in mid-pregnancy (2nd trimester) and when the child was 2 and 12 months old (see “Prospective ELS operationalization” for details). Using z-scores on this index, the original MIDE cohort was divided into four equal strata: low (z < −0.42), moderate-low (−0.42 ≤ z < 0.24), moderate-high (0.24 ≤ z < 0.90), and high (z ≥ 0.90). The target was to recruit 24 participants per stratum, balancing offspring sex and parental infertility history, and including only cases with ≤ 8 missing ELS items. In total, 92 participants were enrolled; when specific cells were depleted, nearby cells were used as replacements. Representation across strata (χ²(3) = 0.35, p = .951) and balance for child sex (χ²(3) = 1.55, p = .671) and parental fertility history (χ²(3) = 0.21, p = .976) were confirmed.

For this overall neuroimaging subsample, both resting-state and task fMRI from 92 young adults aged 18–21 years (M = 19.06, SD = 0.77; 55% female) were collected. Eligibility required right-handedness, native Finnish, normal hearing, normal or corrected-to-normal vision, and no current psychiatric or neurological diagnoses. Three participants exceeded the head-motion threshold (> 0.2 mm mean framewise displacement) and were excluded, yielding a final sample of 89 for the present study. Participants were compensated €15 per hour (total session length 2– 3 h). The fMRI protocol and all prior study phases were approved by the Ethics Committee of the Hospital District of Helsinki and Uusimaa, Finland.

### Early life stress assessment

We employ and analyze two different operationalizations of ELS: Prospective ELS and Retrospective ELS. Prospective ELS was assessed at three time-points: during pregnancy (T1), when the child was 2 months old (T2), and at 12 months (T3), using parent-report questionnaires covering two domains: parental mental health and family relationship functioning. At all three waves, both mothers and fathers completed the General Health Questionnaire (GHQ-36; mothers: α = 0.91–0.94, fathers: α = 0.92–0.94; Goldberg & Hillier, 1979) and the Beck Depression Inventory (BDI-13; mothers: α = 0.75–0.84, fathers: α = 0.80–0.83; Beck et al., 1961). The GHQ indexes depression, anxiety, insomnia, and social dysfunction, whereas the BDI focuses specifically on depressive symptoms. Family relationship difficulties were assessed at T2 and T3 (but not during pregnancy) with the Dyadic Adjustment Scale (DAS; mothers: α = 0.92–0.93, fathers: α = 0.91–0.91; Spanier, 1976) and the Parenting Stress Index (PSI-36; mothers: α = 0.90– 0.90, fathers: α = 0.91–0.91; Abidin, 1997). The DAS captures interparental conflict and low affection; the PSI reflects parenting distress and challenges in the parent–child relationship.

Complete data across the 20 Prospective ELS variables were available for 84% of participants (n = 77); 9% (n = 8) were missing eight variables and 7% (n = 7) were missing one to four variables. Missingness was addressed via Expectation–Maximization (EM) imputation using information from the broader cohort. We then derived a cumulative-risk composite (Ettekal et al., 2019) by averaging each questionnaire across time and across parents, standardizing those averages, and computing their means to obtain the total Prospective ELS score (M = 0.00, SD = 0.86, range = −1.50 to 2.34). The 20-variable set showed good internal consistency (α = 0.88).

Retrospective ELS was assessed using the self-report questionnaire items adapted from the Revised Adverse Childhood Experiences questionnaire (ACES; Finkelhor et al. 2015) approximately a year prior to fMRI data collection when the participants were 17–19 years old (M = 18.23, SD = 0.34). Two items with a three-point Likert scale (0 = never, 1 = sometimes, 2 = often) assessed the following: Emotional abuse (e.g., “Did a parent or other adult in the household … swear at, insult, or put you down?”); Physical abuse (e.g., “Did a parent or other adult in the household … push, grab, shove, or slap you?”); Emotional neglect (e.g., “Did you … feel that no one in your family loved you or thought you were important or special?”); and Parent treated violently (e.g., “Was your parent … pushed, grabbed, slapped, or had something thrown at her/him?”). Two additional items were added to capture common ELS events for interparental psychological violence (e.g., “Have you seen your parent being threatened by violence at home?”; Ellonen et al. 2008). Binary response items (0 = no, 2 = yes) assessed the following: Family alcohol and drug problems (“Did you live with anyone who was a problem drinker or alcoholic, or who used street drugs?”); Peer victimization (“Have you been bullied in school?”); Parents’ divorce (“Did your parents separate/divorce?”); Family mental illness (“Was a household member mentally ill?”); Death of a close person (“Have you ever lost anyone close to you by death?”); Family somatic illness (“Has any family member had a serious illness during your life?”); and Other serious adversities (“Have you experienced other adversities, such as accidents, victimization, or natural catastrophes?”). The total Retrospective ELS score was calculated by averaging the two-item domains and then summing the scores of the 12 domains (M = 4.45, SD = 2.60; range = 0–11.50). In the current study sample, Retrospective ELS and Prospective ELS were uncorrelated (r = 0.001, p = 0.96).

### Procedure

Participants completed a revised Reading the Mind in the Eyes task (RMET; Baron-Cohen et al., 2001; fMRI implementation adapted from Moor et al., 2012) translated into the native language of the participants (Finnish). The RMET is a forced-choice paradigm comprising four consecutive blocks within a single fMRI run (Figure 1). Each block consisted of 18 trials, yielding a total of 72 trials. Two blocks implemented a mentalizing condition (36 mentalizing trials in total) where participants had to infer a mental state from a photograph stimulus of a person’s eyes by selecting the best-matching word from four alternatives (e.g. “irritated, thoughtful, encouraging, sympathetic”). Two other blocks implemented a perceptual control condition (36 control trials in total) using the same face/eye-region photographs but required a non-mentalizing judgment: participants selected which of four options best described the depicted person’s age-by-sex category (young/old × male/female). Participants responded by pressing one of four buttons on a controller with their right hand. The control condition’s response option positions were altered between trials so participants would have to read which button corresponded to which option in each control trial. Block order was counterbalanced across participants by reversing the block order for half of the participants. Stimulus orders within each block were fixed and therefore identical across participants for each block.

**Figure 1.**
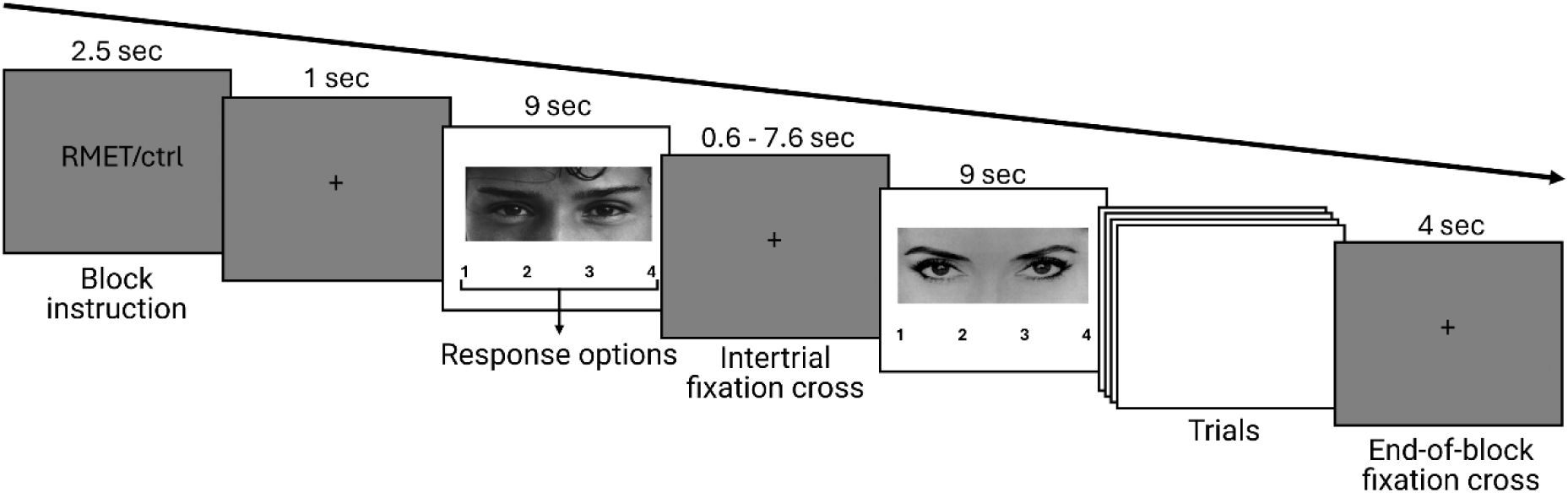
The experimental procedure of a Reading the Mind in the Eyes block. Each of the four blocks, divided into two mentalizing blocks and two control (sex + age assessment using same photograph stimuli) blocks, comprised a brief instruction screen (2.5 seconds), an initial fixation cross (1 second), followed by stimulus presentations (9 seconds) and jittered intertrial fixation crosses (0.6 – 7.6 seconds). In the mentalizing condition, the participant had to choose one of four options that best describes what the person in the photograph is thinking or feeling (e.g., “ärsyyntynyt, miettivä, kannustava, myötätuntoinen”; english version equivalents: “irritated, thoughtful, encouraging, sympathetic”). In the control condition, the participant had to choose one of four options that best describes the age (young/old) and sex (female/male) of the person in the image. Each of the four blocks comprised 18 trials and ended with an end-of-block fixation cross (4 seconds). Total run duration of the fully completed task was approximately 16.6 minutes.

Each block began with an instruction screen presented for 2.5 s. Trials then proceeded in an event-related manner. On each trial, a stimulus photograph was presented for 9 s. A response to each trial was valid if it occurred within the stimulus presentation, before the inter-trial interval. After each stimulus presentation, an inter-trial fixation cross was shown for a jittered duration (0.6 s to 7.6 s; mean ∼4 s). In addition to the jittered ITIs between stimulus presentations, a fixation cross was shown for four seconds after each block’s final stimulus presentation. The experiment was implemented using Presentation (ver. 20.0 build 09.04.17; Neurobehavioural Systems, Inc., Albany, CA, USA). Scanner synchronization was established using the first recorded scanner pulse as time zero for the run. For completed runs, run duration after the first pulse was approximately 16.6 minutes. Response rates were near ceiling, indicating that participants almost always produced a response within the 9 second time window (0.983 and 0.987 in the RMET blocks and 0.999 in both control blocks).

Importantly, some previous studies have raised issues regarding the psychometric properties of the RMET, and thus caution is warranted when interpreting results relating to task performance. For comprehensive evaluations on the psychometric properties of the RMET, see Hafner et al. (2026), Higgins et al. (2023), and Olderbak et al. (2015).

### Behavioral analyses

We tested whether ELS was associated with overall performance on the RMET. Accuracy was defined for each participant as the proportion of correctly answered trials across all RMET trials. Both Prospective and Retrospective ELS indices were examined. Both indices were entered into a separate ordinary least squares regression predicting RMET accuracy, adjusting for background covariates (participant sex, mother’s age, mother’s socioeconomic status (measured as highest completed education level), and ART-status). All predictors were mean-centered prior to model fitting. Inference on the ELS coefficient was based on a two-tailed t-test of the corresponding regression weight; we report the unstandardized coefficient with its 95% confidence interval, the partial correlation as a standardized effect size, and the effective sample size for each model. Models were fit by listwise deletion across the outcome, predictor, and background covariates, yielding N = 89 for the Prospective ELS index and N = 85 for the Retrospective ELS index.

Ordinary least squares was pre-specified as the primary estimator and applied identically to both indices. Because the two adversity measures index distinct exposures assessed by different methods and at different points in development, and were empirically uncorrelated in this sample, they were treated as separate hypotheses rather than as a single test family, and p-values are reported uncorrected.

### MRI acquisition

Imaging took place at the Advanced Magnetic Imaging (AMI) Centre, Aalto NeuroImaging (Aalto University School of Science, Espoo, Finland) on a 3T Siemens MAGNETOM Skyra with a 20-channel head coil. Across the session we acquired task fMRI, structural MRI, and resting-state data. For all participants the session began with a go/no-go task (see Ilomäki et al., 2025), followed by the Reading the Mind in the Eyes Task. Next, a high-resolution T1-weighted anatomical scan was obtained (MPRAGE; 3D, matrix 256 × 256; 1 mm isotropic voxels). Participants then completed a social-media paradigm in which they posted opinions to a sham Facebook group and received peer feedback (see Wikman et al., 2022). The session concluded with resting-state fMRI during which participants lay still with eyes open (see Ilomäki et al., 2022).

All functional runs used whole-brain echo-planar imaging (EPI) with 43 contiguous oblique slices (TR = 2,500 ms; TE = 32 ms; flip angle = 75°; matrix 64 × 64; field of view = 20 cm; slice thickness = 3.0 mm; in-plane resolution = 3.125 × 3.125 × 3.0 mm).

### Preprocessing

Structural and functional MRI data were preprocessed using fMRIPrep v20.2.5 (Esteban et al., 2019). T1-weighted (T1w) images were corrected for intensity non-uniformity using N4BiasFieldCorrection (ANTs 2.3.3; Tustison et al., 2010; Avants et al., 2008) and skull-stripped with the ANTs antsBrainExtraction.sh workflow (Nipype implementation) using the OASIS30ANTs template. Brain tissue segmentation into gray matter (GM), white matter (WM), and cerebrospinal fluid (CSF) was performed on the skull-stripped T1w image with FSL FAST (5.0.9; Zhang et al., 2001). For surface-based analyses, cortical surface reconstruction was performed with FreeSurfer (recon-all; Dale el al., 1999). Nonlinear normalization of anatomical images to standard space(s) (MNI152NLin6Asym and MNI152NLin2009cAsym) was carried out with antsRegistration (ANTs 2.3.3) using brain-extracted T1w images.

For each BOLD run, fMRIPrep generated a reference volume and its skull-stripped counterpart. Susceptibility distortion correction used a fieldmap-less approach in which the BOLD reference was co-registered to the participant’s T1w reference with inverted contrast (Huntenburg, 2014; Wang et al., 2017) via antsRegistration, with the deformation constrained to the phase-encoding axis and informed by an average fieldmap template (Treiber et al., 2016). The distortion-corrected BOLD reference was then registered to the T1w image using FreeSurfer’s bbregister (boundary-based registration; Greve & Fischl, 2009) with six degrees of freedom. Head motion parameters were estimated relative to the BOLD reference prior to any spatiotemporal filtering using FSL MCFLIRT (5.0.9; Jenkinson et al., 2002), and slice-timing correction was applied using AFNI 3dTshift (20160207; Cox & Hyde, 1997). Volumetric resampling applied a single composite transform combining motion correction, susceptibility distortion correction, and BOLD-to-T1w registration; data were resampled to each subject’s T1w space and to standard space as required. All volumetric resampling used antsApplyTransforms (ANTs) with Lanczos interpolation to minimize smoothing (Lanczos, 1964). In addition, BOLD time series were sampled to FreeSurfer surface space and resampled to a common fsaverage surface for surface-based analyses.

Confound time series provided by fMRIPrep included framewise displacement (FD; both Power’s absolute-sum formulation and Jenkinson’s RMS displacement; Power et al., 2014; Jenkinson et al., 2002), DVARS, and mean signals from CSF, WM, and whole-brain masks (global signal). In addition, component-based noise regressors (aCompCor; Behzadi et al., 2007) were extracted.

### First-level analysis

We estimated two complementary first-level general linear models (GLMs) from the preprocessed data using FEAT (FSL v6.00). A surface-based GLM was used for the univariate analyses, whereas a volumetric event-related GLM was used for all multivariate analyses (searchlight RSA, ROI-based RSA, IS-RSA, connectivity, and manifold geometry). Both models used fMRIPrep derivatives in their respective target spaces (fsaverage for the surface model and MNI152NLin2009cAsym for the volumetric model). Both models applied a 100 s high-pass temporal filter and FILM prewhitening (Woolrich et al., 2001). Event regressors used the standard three-column format (onset, duration = 5 s, amplitude = 1) and were convolved with the canonical gamma hemodynamic response function. Identical fMRIprep derivated nuisance regressors entered both GLMs: six rigid-body motion parameters, three anatomical aCompCor components, and framewise displacement (with the undefined first-volume FD value set to 0). Subjects with mean framewise displacement exceeding 0.2 mm were excluded prior to first-level estimation, leaving N = 89 subjects.

For the surface-based first level analysis, fsaverage-resampled BOLD time series (left and right hemispheres modeled separately; full-density mesh, ∼160k vertices per hemisphere) were spatially smoothed at 5 mm FWHM along the surface prior to model estimation. The design included five event regressors with temporal derivatives: RMET-correct, RMET-incorrect, control-correct, control-incorrect, and a button-press response regressor. Seven contrasts were estimated per hemisphere: each of the four task conditions individually, Task (RMET > control), Accuracy (correct > incorrect), and a Task × Accuracy interaction.

For the volumetric trial-wise model, we fit a least-squares-all GLM in which each of the 72 trials comprising the run were modeled as their own event regressors within a single model, yielding one cope per trial via single-regressor t-contrasts against the implicit baseline. Temporal derivatives were not included in this model to conserve degrees of freedom in the dense 72-regressor design. Instruction screens and inter-trial intervals were left unmodeled and contributed to baseline. The choice of nuisance regressors followed recommendations for beta-series modeling: motion parameters, FD, and anatomical aCompCor components were retained, while global signal regression was deliberately omitted, since GSR can remove shared neural variance that contributes to representational geometry across trials (Mumford et al., 2012; Aquino et al., 2020).

### Second-level analyses

#### Univariate group-level within-subjects analysis

Group-level surface-based analyses were performed using FreeSurfer [v6.0.0] (mri_glmfit). Subject-level contrast maps were entered into a one-sample group model (ordinary least squares) testing whether the mean contrast estimate differed from zero across participants (N = 89). Vertex-wise t-statistics were computed separately for each hemisphere.

Correction for multiple comparisons was performed with mri_glmfit-sim using a vertex-wise cluster-forming threshold of p < .001 (−log₁₀(p) > 3.0) and FreeSurfer’s precomputed cluster-size distributions (Z Monte Carlo simulation, 10,000 iterations; Hagler et al., 2006; Greve & Fischl, 2018). The simulation appropriate to the smoothness estimated from the group-level model residuals was selected automatically. Positive and negative tails were tested separately for each contrast, and each hemisphere was corrected independently.

A cluster-wise threshold of p < .025 was applied within each hemisphere and each direction. Because each combination of hemisphere and direction constitutes a separate family, this threshold controls the family-wise error rate at .05 across the two hemispheres but not additionally across the two directions; clusters not also meeting the more conservative threshold of p < .0125, which would control the family-wise error rate at .05 across both hemispheres and both directions, are flagged in the supplementary tables and treated as exploratory. Surviving clusters were retained as vertex-wise masks. Automatic anatomical labels denote the Desikan– Killiany (aparc) parcel containing each cluster’s peak vertex.

### Univariate group-level between-subjects analysis

For both Prospective and Retrospective ELS index we fit two between-subjects models. The unadjusted model contained only the ELS predictor and estimates the total association between ELS and the contrast estimate. The adjusted model additionally included background covariates of no interest, and estimates the component of that association independent of these variables.

Adjusted models were pre-specified as primary; unadjusted models were fit to establish whether covariate inclusion masked or created effects.

Participants were filtered per model to those with complete data on all model terms (Prospective ELS models: N = 89; Retrospective ELS models: N = 85). All continuous variables were mean-centered prior to entry into the design matrix. Group-level analyses used FreeSurfer’s FSGD framework with the dods parameterization, and the contrast tested the ELS slope (zero on the intercept, unity on the ELS term, zero on all covariate slopes). Cluster correction used parameters identical to the within-subjects analyses (vertex-wise threshold p < .001; precomputed Z Monte Carlo simulation, 10,000 iterations; cluster-wise p < .025 within each hemisphere and direction; both tails tested).

### Representational Similarity Analysis

#### Neural RDM construction

To characterize local multivariate representational structure, we conducted searchlight RSA in volumetric MNI152NLin2009cAsym space. For each participant, the 72 trial-wise COPE images from the beta-series GLM served as input. All COPE images were verified to share an identical voxel grid (shape and affine). Searchlight centers were restricted to gray matter using a group-level mask. For each participant, the fMRIPrep-derived gray matter probability map (label-GM_probseg.nii.gz) was resampled to the common functional grid using nearest-neighbor interpolation, averaged voxel-wise across participants, and thresholded at 0.25 to yield a binary group GM mask. This threshold ensured that searchlight centers fell within cortical and subcortical gray matter while accommodating inter-individual variability in tissue boundaries. Spherical searchlight neighborhoods were defined using rsatoolbox (rsatoolbox.util.searchlight.get_volume_searchlight) with a radius of 3 voxels (approximately 9 mm at 3.125 × 3.125 × 3.0 mm anisotropic resolution, yielding ∼123 voxels per sphere) and a coverage threshold of 0.50, requiring at least 50% of voxels within each sphere to fall inside the same group GM mask.

Prior to computing representational dissimilarities, the trial-wise COPE data were flattened from four-dimensional arrays (72 trials × X × Y × Z) into two-dimensional matrices (72 trials × voxels). Voxels containing NaN values in any trial were identified and set to zero across all trials, effectively excluding them from subsequent computations by contributing a constant (zero-variance) pattern. Each voxel was then z-scored across trials. Although correlation distance is intrinsically insensitive to voxel-level mean differences in BOLD response (Popal et al., 2019), per-voxel z-scoring additionally equalizes across-trial variance across voxels within each searchlight, preventing strongly task-modulated voxels from disproportionately driving the resulting representational distances. We include this normalization deliberately, on the grounds that subsequent between-subjects inference targets pattern geometry rather than activation magnitude in any small subset of voxels (cf. Diedrichsen & Kriegeskorte, 2017). Voxels with zero variance across trials were assigned values of zero rather than undefined quantities to prevent numerical instabilities.

For each searchlight sphere, a 72 × 72 trial-by-trial representational dissimilarity matrix (RDM) was computed using correlation distance (1 − Pearson r). The upper triangle was extracted as a representational dissimilarity vector (RDV) of length 2,556 unique pairwise distances. The RDV, along with three-dimensional voxel coordinates and image affine, was saved per participant for subsequent model comparison.

To test whether local neural representations distinguished mentalizing from the control task, we constructed a binary model RDM encoding task-condition membership. The 72 trials comprised 36 task (mentalizing) trials and 36 control (age-by-sex categorization) trials. The model RDM assigned a dissimilarity of 0 to within-condition trial pairs (RMET–RMET and control–control) and a dissimilarity of 1 to between-condition pairs, yielding a categorical partition of representational space according to task demands. The upper triangle was vectorized to produce a model RDV of 2,556 elements matching the dimensionality of the neural RDVs.

As our primary searchlight statistic, we computed the mean between-condition dissimilarity minus the mean within-condition dissimilarity (Nili et al., 2014), a direct index of how strongly local pattern geometry separates the two task conditions. Positive values indicate that between-condition trial pairs are more dissimilar in their local activity patterns than within-condition pairs, which is the RSA signature of condition-discriminative representation. The resulting value at each searchlight was written into a three-dimensional NIfTI volume in MNI152NLin2009cAsym space, yielding one RSA effect map per participant.

#### Within-subjects group-level inference

Group-level statistical inference on the searchlight RSA maps was performed using FSL’s randomise (Winkler et al., 2014) with threshold-free cluster enhancement (TFCE; Smith & Nichols, 2009). Because task-evoked pattern differences in this paradigm were spatially diffuse across the cortex, an unadjusted one-sample test against zero produced extensive significant effects that did not usefully discriminate regions of interest. To focus inference on regions with spatially specific model–brain correspondence, each participant’s RSA effect map was centered by subtracting the mean across all finite in-mask voxels before group analysis. This reframes the null hypothesis from “RSA effect > 0” to “RSA effect exceeds this participant’s whole-brain average,”. Non-finite voxels were set to zero to comply with FSL’s input requirements, and a group mask was constructed by retaining voxels with finite values across all participants.

A one-sample design was specified with a single column of ones. Two directional tests were conducted: a positive contrast testing whether the group-mean centered RSA value was greater than zero, and a negative contrast testing whether it was less than zero. Each tail was evaluated with 20,000 sign-flip permutations and TFCE correction, with the two tails Bonferroni-corrected (1 − p > 0.975 per tail) for an effective two-tailed FWE p < 0.05.

#### Between-subjects group-level inference: ELS moderation

Voxel-wise whole-brain searchlight inference (Figure 2A) for between-subjects effects is statistically demanding, and ELS effects on representational patterns are expected to be subtle relative to inter-individual variability in the precise anatomical locations of higher-level cognition (Frost & Goebel, 2012). Thus, to test whether ELS exposure moderated local representational patterns with greater statistical power and tolerance for inter-subject (functional and anatomical) variability, we conducted between-subjects analyses at the parcel level defined by the HCP-MMP 1.0 cortical parcellation (Glasser et al., 2016).

**Figure 2.**
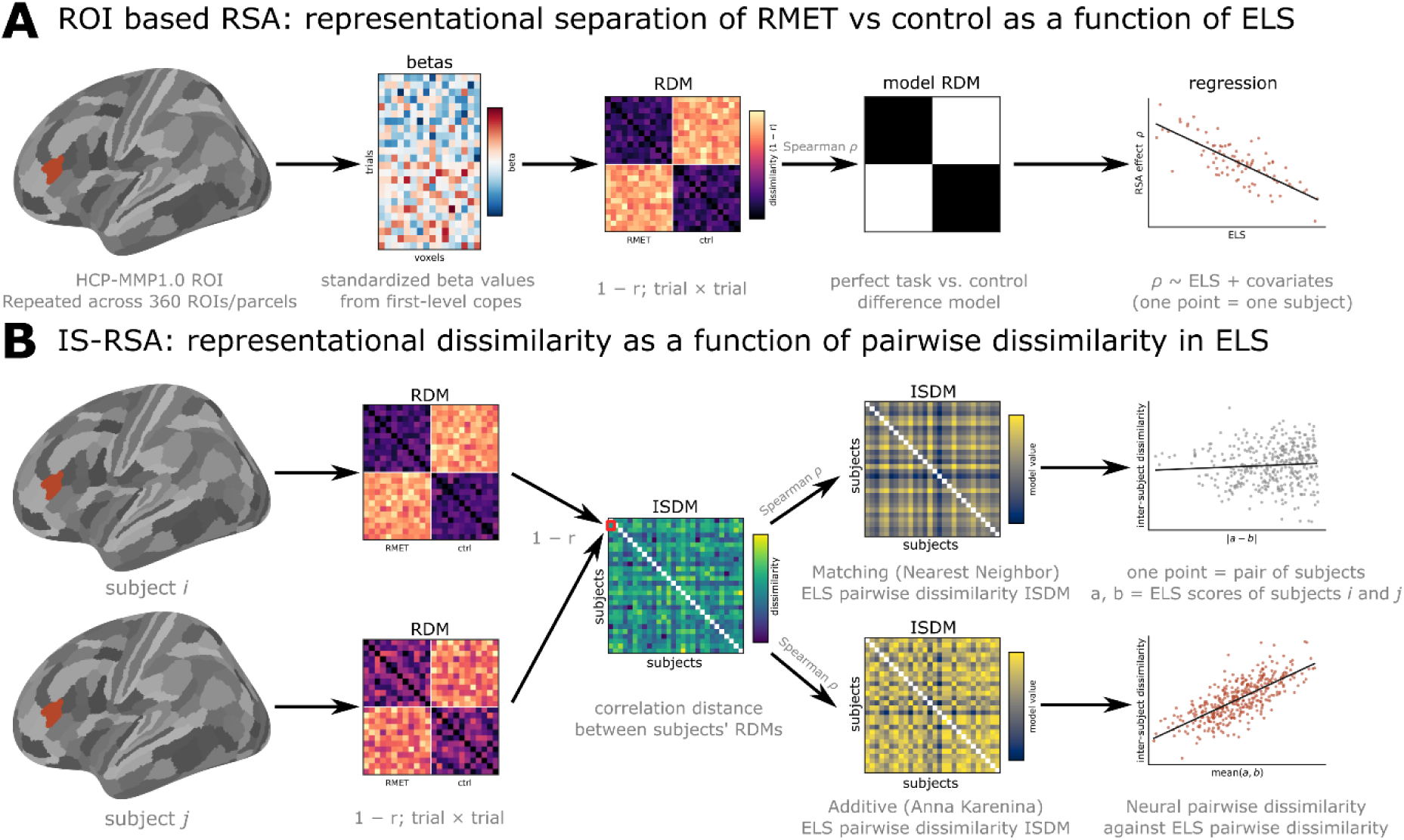
A schematic of the between-subjects analyses of the association between early life stress (ELS) and neural representations. Both analyses were performed for each brain region defined by the HCP-MMP1.0 cortical parcellation that divides the cortex into 360 regions (180 per hemisphere). In the subject-level analysis (A), for each participant and region, a trial × trial representational dissimilarity matrix (RDM; 72 × 72) was computed as the correlation distance (1 – r) between the standardized multivoxel beta patterns of each pair of trials. Each RDM was compared with a model RDM encoding a perfect mentalizing versus non-mentalizing distinction (white = same condition, black = different condition) using a rank-order correlation (Spearman ρ) over the upper-triangle elements. The resulting subject-level RSA effect was then regressed on ELS, adjusting for background covariates (mother’s age and socioeconomic status, participant’s sex, and assisted reproductive technology status). Each point in the scatterplot represents one participant. In the inter-subject analysis (B), inter-subject dissimilarity matrices (ISDMs) encoding pairwise dissimilarity between all pairs of participants were constructed for both ELS and neural representations. Neural ISDM values were computed for each region as the correlation distance (1 − r) between the vectorized upper triangles of the two participants’ RDMs. Two pairwise approaches were used for ELS: a nearest-neighbor (”matching”) model, in which each pair was assigned the absolute difference between the participants’ ELS values, and an Anna Karenina (”additive”) model, in which each pair was assigned the participants’ mean ELS value; a positive additive effect indicates that individuals with higher ELS have increasingly idiosyncratic representations. Neural ISDM association with ELS ISDMs were analyzed with partial Spearman correlation, adjusting for the background covariates listed in (A), each entered as a covariate ISDM constructed under the same pairwise coding. Each point in the scatterplots represents one unique pair of participants.

For each participant, voxel-level patterns were aggregated within each of the 360 HCP-MMP cortical parcels. Voxel-to-parcel assignment was performed by nearest-neighbor lookup against the volumetric HCP-MMP atlas, restricted to voxels falling within the same group GM mask used for the searchlight pipeline. As in the searchlight pipeline, each voxel was z-scored across trials prior to correlation-distance computation, and voxels with zero variance across trials (including NaN-zeroed voxels) were dropped within each ROI. For each ROI, a 72 × 72 RDM was computed using correlation distance (1 − Pearson r), and the per-subject RSA effect for the task-vs-control model was computed as the mean between-condition dissimilarity minus the mean within-condition dissimilarity, matching the searchlight statistic.

Per-subject RSA effects were mean-centered across ROIs prior to between-subjects inference, focusing the test on regional specificity relative to each participant’s brain-wide average representational structure (see above). For each of the two ELS measures (Prospective and Retrospective), two model flavors were estimated: a bare model with ELS as the sole predictor, and an adjusted model that included background covariates. All continuous predictors were mean-centered prior to model fitting. Per-model subject filtering retained only participants with complete data on all model terms, yielding N = 89 for all Prospective ELS analyses and N = 85 for all Retrospective ELS analyses (due to missing Retrospective ELS data for four participants).

Inference at each ROI was conducted via 20,000 permutations of subject labels on the ELS variable, with the regression slope on the residualized ELS predictor (after partialling out covariates from both predictor and outcome) recomputed for each permuted dataset. Family-wise error correction across the 360 HCP-MMP ROIs was performed via the maximum-statistic null distribution. Both positive and negative ELS-slope contrasts were tested, with two-tailed inference established via Bonferroni correction over the two tails (per-tail p < 0.025) to maintain family-wise error at 0.05 per model. Benjamini-Hochberg false discovery rate correction across ROIs was additionally computed as a secondary criterion.

#### Inter-subject Representational Similarity Analysis

Beyond testing whether ELS modulates the strength of the task-versus-control representational distinction in any specific ROI, we additionally tested whether inter-subject similarity in ELS exposure was associated with inter-subject similarity in local representational structure independent of an a priori model RDM. This question is complementary to the between-subjects ROI analysis: rather than asking how ELS modulates one representational contrast, inter-subject representational similarity analysis (IS-RSA; Finn et al., 2020) asks how ELS structures the inter-subject space of full representational dissimilarity matrices.

ROI-level IS-RSA was conducted on the same HCP-MMP1.0 ROI RDMs constructed for the between-subjects ROI analysis described above. For each ROI, an inter-subject neural dissimilarity matrix (ISDM) was constructed by computing 1 − Spearman *ρ* between each pair of participants’ vectorized upper triangles of their RDMs, yielding one dissimilarity value for every participant pair (Figure 2B).

Predictor ISDMs were constructed from each ELS index under two complementary models that make different claims about how exposure structures inter-subject neural similarity. A “matching” model, corresponding to the nearest-neighbor (NN) formulation of Finn et al. (2020), defined pairwise predictor distance as the absolute difference between participants’ ELS values, |a − b|. It tests the hypothesis that participants with similar exposure share representational structure, at any point on the exposure continuum. An “additive”, or Anna Karenina (AK), model defined pairwise distance as the mean of the two participants’ ELS values, (a + b)/2. It tests the hypothesis that representational structure becomes systematically more (or less) idiosyncratic as exposure increases.

Directional hypotheses were declared a priori and differ between the two statistics. The matching (NN) statistic was tested one-tailed positive: participants with similar exposure have similar representational geometry. The additive (AK) statistic was tested two-tailed. Because the outcome is neural dissimilarity, a positive additive effect (AK+) indicates that pairs high in mean exposure are more dissimilar, that is, representational idiosyncrasy at high ELS and convergence at low ELS, while a negative effect (AK−) indicates convergence at high exposure.

For both Prospective and Retrospective ELS index we estimated an unadjusted model (predictor ISDM only) and an adjusted model including background covariate ISDMs. Continuous covariates were entered as absolute pairwise differences; binary covariates as 0/1 mismatch indicators. The IS-RSA statistic was a partial Spearman correlation, implemented as the linear-on-ranks approximation standard in the literature: the neural ISDM, the predictor ISDM, and each covariate ISDM were rank-transformed across participant pairs; the ranked neural and ranked predictor vectors were then linearly residualised against the ranked covariate design matrix (intercept-only in unadjusted models); and the partial Spearman *ρ* was computed as the Pearson correlation between the standardized rank residuals.

Inference was conducted by 20,000 permutations of participant labels on the ELS index, with the entire rank-first residualisation procedure (including construction of the predictor ISDM) recomputed for each permuted predictor. The same permutations were shared across statistics and across parcels, so that the dependence between them is preserved in the null distribution.

Family-wise error was controlled at two levels using the maximum-statistic null distribution. The first (p FWE-P) takes the maximum across the 360 HCP-MMP parcels within a single statistic. The second (p FWE-F) takes the maximum across parcels and across both statistics (matching + additive) together, using the signed *ρ* for the one-tailed matching statistic and |*ρ*| for the two-tailed additive statistic. The two ELS indices were treated as conceptually distinct exposures rather than as one family, and no correction was thus applied across the. The unadjusted and adjusted models of the same index are nested variants of one analysis rather than independent tests and are likewise not corrected against each other. Finally, Benjamini–Hochberg false discovery rate across parcels is additionally reported in the supplementary table for the results.

The matching and additive statistics were estimated separately, each with a single ELS predictor in the model, and inference was conducted on these single-predictor statistics as described above. However, we additionally fitted a combined model containing both predictors,

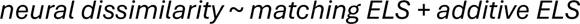

estimated by ordinary least squares on the same rank-transformed, covariate-residualised and standardized vectors. This combined model’s coefficients each estimate the association of one predictor with neural dissimilarity while holding the other constant, in addition to the background covariate adjustment applied throughout. In simple terms, it asks whether an effect reflects pairwise similarity in exposure between two participants or the average exposure level of the pair once both compete for the same variance. This model was specified after the confirmatory analysis, was not included in the family-wise correction, and is reported descriptively to characterize established effects.

#### Representational geometry characteristics

To characterize the inter-subject RSA result, we projected each subject’s representational geometry into a shared two-dimensional embedding and examined how that geometry varied with ELS. For each subject, the regional trial-by-trial RDM (72 × 72; 1 − Pearson *r* between beta-series patterns) was embedded in two dimensions using classical multidimensional scaling (MDS). Classical MDS recovers a point configuration whose pairwise Euclidean distances approximate the RDM as closely as a given number of dimensions allows. Because 72 trials define a geometry of far higher dimensionality than two, some structure is lost by the embedding. We quantify the loss as the proportion of positive eigenvalue mass carried by the first two dimensions and report it in the results.

Importantly, an MDS solution is determined only up to rotation and reflection: two subjects with highly similar representational geometry can produce embedding configurations that appear unrelated, because the axes returned by the algorithm carry no shared meaning between subjects. We resolved this with generalized Procrustes analysis (GPA). In GPA, first, each embedding configuration was mean-centered and then rotated, with reflection permitted, onto a common reference configuration. Then, the reference was updated to the mean of the aligned configurations and the procedure repeated until convergence. The reference was initialized with the classical MDS solution of the group-mean RDM. Alignment was restricted to rotation and reflection, while scaling was deliberately excluded. Rescaling each configuration to match the group mean would minimize subject-wise differences in configuration size into the fit, losing necessary information for establishing condition (task vs. control) separation. To confirm that the restriction did not distort the result, we verified that condition separation measured in the aligned two-dimensional frame agreed with the equivalent quantity computed on the full RDMs.

For transparency, we report three properties of the GPA alignment: the median rotation applied, the number of subjects requiring a reflection, and the reduction in mean distance between individual configurations and the group mean (which indexes how much of the between-subject disagreement was attributable to only orientation).

GPA aligned subject-wise embeddings were then stacked along a third axis denoting Prospective ELS. For display, each trial’s aligned coordinate was regressed on ELS across subjects and the fitted line plotted; condition centroid trajectories were obtained the same way, from the subject-wise RMET and control centroids. These lines are thus cross-sectional fits across subjects, not within-subject trajectories: no subject travels along any particular line displayed, and they should be read as showing how geometry differs between people at different ELS levels, not how any individual’s representational geometry changes.

Additionally, two quantities were computed on the full RDMs, without reference to the MDS embeddings: condition separation and typicality. Condition separation was calculated as the mean representational dissimilarity between trials of different conditions minus the mean dissimilarity between trials of the same condition, divided by the overall mean dissimilarity of that subject’s RDM. Scaling by the overall mean makes the index comparable across subjects with different absolute dissimilarity levels. Higher values indicate more sharply differentiated task versus control representations. Typicality, on the other hand, was derived from the inter-subject distance matrix (ISDM), the matrix of Spearman correlation distances between subjects’ vectorized RDMs. Here, each subject’s mean distance to all other subjects indexes how much that subject’s geometry differs from the group as a whole. We report this as typicality, defined as one minus that mean distance, so that higher values indicate a geometry closer to the group norm. Both quantities were regressed on ELS with adjustment for background covariates and are reported as partial correlations.

## Results

### Behavioral results

The sample average RMET accuracy was .748 (SD = .093, range .417–.917). Between-subjects, we tested whether Prospective or Retrospective ELS indices predicted RMET accuracy using linear regression, controlling for background covariates (participant sex, mother’s age, mother’s SES, and ART-status). The Prospective and Retrospective ELS measures were entered into separate models and treated as distinct exposures rather than as a single test family; p-values are therefore reported uncorrected (see Methods).

Higher Prospective ELS was associated with greater RMET accuracy (*b* = 0.024, 95% CI [0.001, 0.048], partial *r* = .22, *t*(83) = 2.05, *p* = .044, *N* = 89; Figure 3A). Retrospective ELS showed an association in the same direction, but weaker and not statistically significant (*b* = 0.005, 95% CI [−0.002, 0.013], partial *r* = .15, *t*(79) = 1.35, *p* = .18, *N* = 85; Figure 3B). The two ELS indices were themselves uncorrelated (*ρ* = .05, *p* = .66, N = 85).

**Figure 3.**
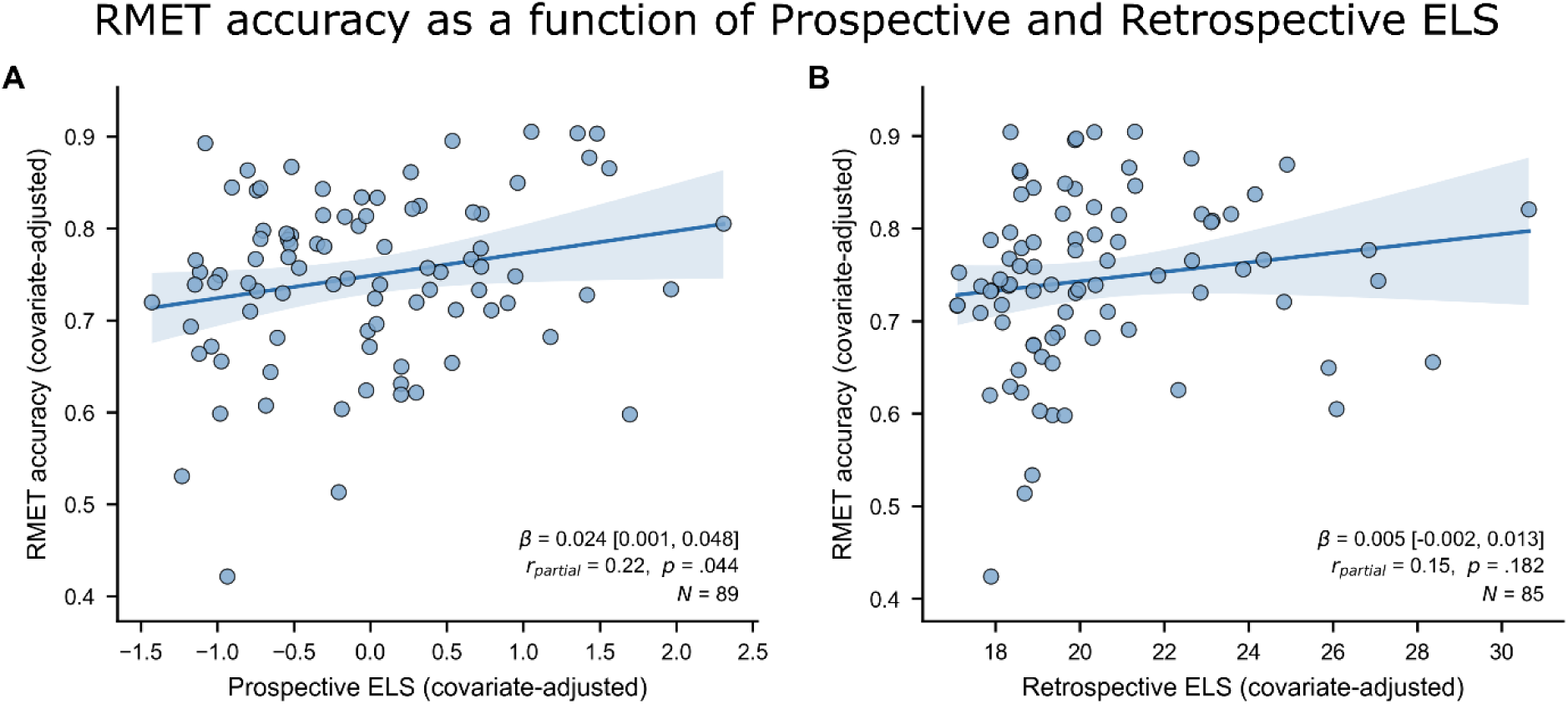
RMET accuracy as a function of Prospective and Retrospective ELS. Added-variable (partial regression) plots showing per-subject RMET accuracy against Prospective ELS (A) and retrospective ELS (B). Higher Prospective ELS was associated with greater RMET accuracy (*b* = 0.024, 95% CI [0.001, 0.048], *t*(83) = 2.05, *p* = .044); retrospective ELS showed a positive but non-significant association (*b* = 0.005, 95% CI [−0.002, 0.013], *t*(79) = 1.35, *p* = .18). Both accuracy and each ELS index are residualised on the background covariates (participant sex, mother’s age, mother’s socioeconomic status, and assisted reproductive technology status) and mean centered. Each point represents one participant; blue lines show the fitted regression, shaded bands the 95% confidence interval.

### Univariate neural results

#### Within-subjects task activation

The mentalizing > control contrast (Figure 4) produced robust bilateral activation across a distributed network consistent with prior work on mentalizing (Schurz et al., 2021; Molenberghs et al., 2016; Schurz et al., 2014).

**Figure 4.**
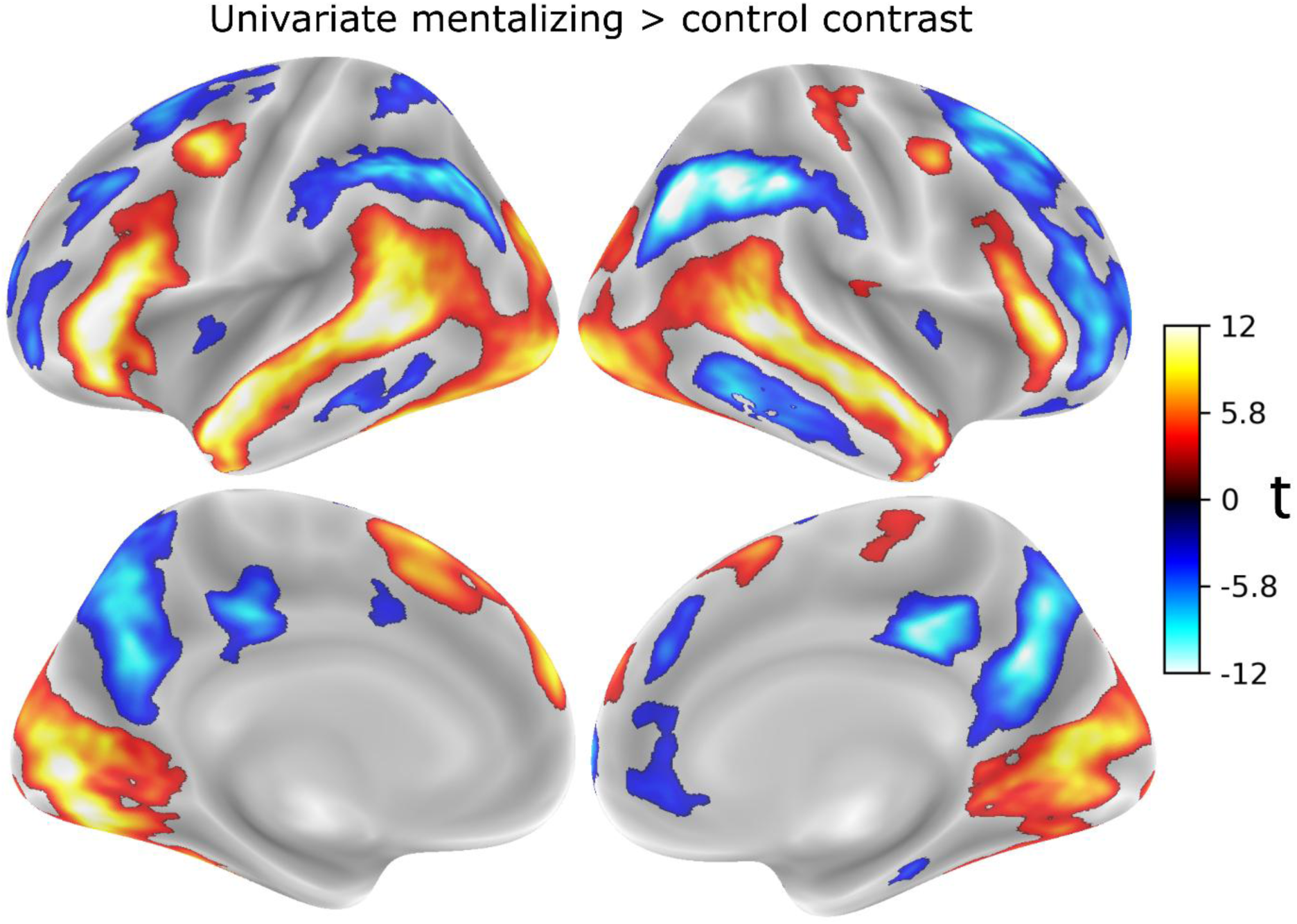
Cortical surface map of mentalizing versus control activation in the RMET paradigm. Group-level t-statistics for the mentalizing > control contrast (*N* = 89, df = 88), displayed on the inflated fsaverage surface (top row, lateral views; bottom row, medial views; left column, left hemisphere; right column, right hemisphere). Maps are thresholded using a vertex-wise cluster-forming threshold of *p* < .001 with cluster-wise family-wise error correction at *p* < .025 within each hemisphere and each direction; vertices not surviving correction are shown in grey. The color scale is truncated at |*t*| = 10 for display; peak values reach *t* = 13.82. Warm colors indicate greater activation during mentalizing than control trials; cool colors indicate the reverse. Mentalizing trials engaged bilateral superior temporal cortex extending along the superior temporal sulcus, inferior frontal gyrus (pars triangularis), dorsomedial superior frontal cortex, and left caudal middle frontal cortex, consistent with prior work on the cognitive neuroscience of mental-state attribution. Relative deactivation was observed across default mode regions, including precuneus, posterior cingulate cortex, and inferior parietal and supramarginal cortex bilaterally, together with bilateral insula.

Positive-tail activation was dominated in each hemisphere by a single very large temporal cluster: in the left hemisphere centered on the banks of the superior temporal sulcus (15991.9 mm², 26,420 vertices; peak *t* = 13.77 at (−50, −41, 4)), and in the right hemisphere a comparably extensive cluster whose peak fell in anterior superior temporal cortex (13401.1 mm², 22489 vertices; peak *t* = 13.00 at (50, −4, −17)). Inferior frontal gyrus (pars triangularis) was engaged bilaterally (left: 3312.7 mm², peak *t* = 13.82 at (−52, 30, 7); right: 1369.8 mm², peak *t* = 11.72 at (54, 27, 8)); notably, the left inferior frontal peak was the strongest in the map, exceeding that of the more extensive superior temporal cluster. Dorsomedial superior frontal cortex was engaged bilaterally (left: 1626.6 mm², peak *t* = 9.32 at (−8, 58, 27); right: 206.7 mm², peak *t* = 6.13 at (8, 56, 31)), together with left caudal middle frontal cortex (830.3 mm², peak *t* = 10.37 at (−43, 3, 48)). Smaller right-hemisphere clusters were observed in postcentral (peak *t* = 5.48), precentral (peak *t* = 8.02), paracentral (peak *t* = 4.30), and supramarginal cortex (peak *t* = 4.19).

The negative tail encompassed the canonical default mode network bilaterally. The largest clusters were left inferior parietal cortex (3290.1 mm², peak *t* = −9.81 at (−43, −63, 44)) and left precuneus (3270.2 mm², peak *t* = −9.92 at (−4, −66, 28)), with right-hemisphere homologues in supramarginal cortex (4223.2 mm², peak *t* = −13.48 at (52, −40, 44)) and precuneus (2047.1 mm², peak *t* = −12.36 at (5, −59, 28)). Posterior cingulate cortex was deactivated bilaterally (left: 509.6 mm², peak *t* = −8.89; right: 678.2 mm², peak *t* = −11.32), as were dorsal and rostral prefrontal regions, including a large right superior frontal cluster (5421.9 mm², peak *t* = −10.96 at (21, 24, 53)) and three separate left rostral middle frontal clusters (714.8, 704.1, and 374.6 mm²; peak *t* = −6.60, −7.15, and −6.75). Bilateral insula (left: peak *t* = −4.02; right: peak *t* = −5.09), right lateral orbitofrontal cortex (179.7 mm², peak *t* = −4.38), right parahippocampal cortex (113.5 mm², peak *t* = −5.24), and lateral temporal cortex (left middle temporal: 485.0 mm², peak *t* = −6.01 at (−62, −35, −16); right inferior temporal: 1304.3 mm², peak *t* = −8.68 at (55, −50, −12)) also showed relative deactivation during mentalizing.

Full cluster tables for the correct > incorrect and condition × correctness interaction contrasts are provided in Supplementary Table S1, with surface visualizations in Supplementary Figure S1.

#### Between-subjects ELS moderation of task activation

Prospective ELS did not moderate any of the three contrasts. No cluster survived correction in either hemisphere, in either tail, in either the adjusted or the unadjusted model, for the mentalizing > control, correct > incorrect, or interaction contrasts.

Retrospective ELS yielded a single cluster across the entire between-subjects analysis, but only in the unadjusted model (Supplementary Figure S1). Higher Retrospective ELS scores were associated with a smaller mentalizing > control difference in left middle temporal cortex (104.6 mm², 216 vertices; peak *t* = −4.63, df = 83, at (−51, −36, −9); cluster-wise *p* = .0166, 90% CI [.0150, .0182]). No cluster reached the corrected threshold anywhere in the adjusted model, and no cluster survived for either model in the remaining two contrasts, in the right hemisphere, or in the positive tail.

### Representational Similarity Analysis Results

#### Searchlight RSA: within-subjects task-vs-control representation

After within-subject centering and whole-brain FWE correction, the task-versus-control model RDM was significantly represented across a distributed set of cortical regions (Figure 5, *p* < .05 TFCE-FWE). Regions showing reliably stronger-than-average model–brain correspondence comprised the canonical mentalizing network, thus substantially overlapping with the regions identified in the positive tail of the univariate task-vs-control contrast: [bilateral/left/right] precuneus, dorsomedial prefrontal cortex, inferior frontal gyrus, and regions along the superior temporal sulcus.

**Figure 5.**
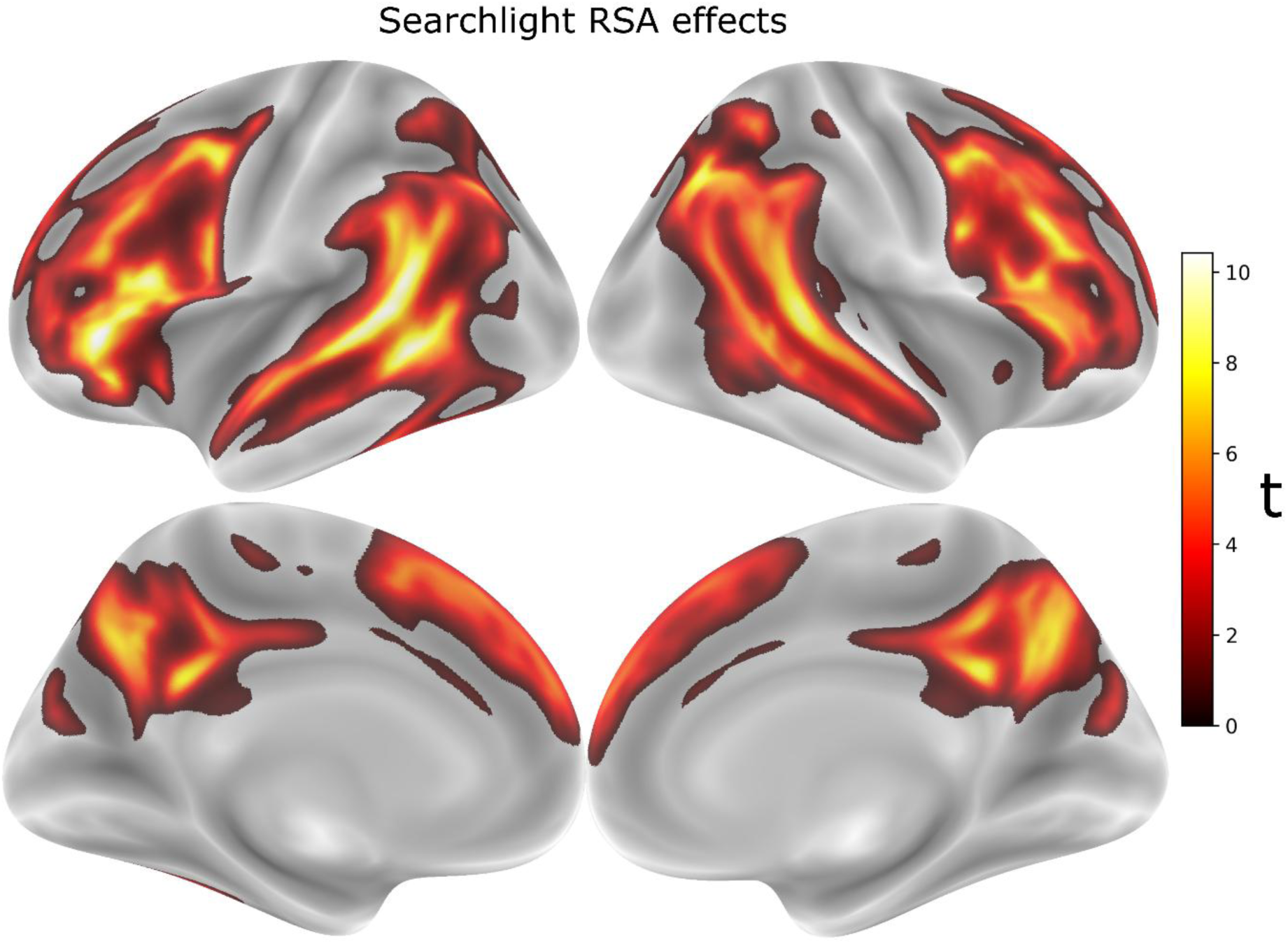
Within-subjects representational similarity for RMET versus control trials. Group-level t-statistic map from a searchlight RSA contrasting representational dissimilarity for RMET versus control trials, thresholded at cluster-corrected *p* < 0.05 (TFCE) and projected to the inflated fsaverage surface (top: lateral views; bottom: medial views). Significant clusters were found bilaterally in posterior superior temporal sulcus (pSTS), temporoparietal junction (TPJ), dorsomedial prefrontal cortex (dmPFC), inferior frontal gyrus (IFG), dorsolateral prefrontal cortex (dlPFC), and precuneus, corresponding to the canonical mentalizing network. The t-statistic map was smoothed with a 5mm FWHM Gaussian kernel prior to surface projection for visualization purposes only.

#### Between-subjects ROI-level RSA: ELS moderation

To assess whether ELS exposure moderated local representational structure, we conducted between-subjects analyses at the level of HCP-MMP cortical parcels. For Prospective ELS in the adjusted model, one ROI surviving FWE correction was identified: left inferior frontal sulcus, anterior division (lh_IFSa; Pearson *r* = -0.40, FWE-corrected *p* = .019, FDR-corrected *q* = .018; Figure 6). The negative slope indicates that participants with higher Prospective ELS showed smaller model-to-neural RDM correlations. No other ROIs survived FWE correction in any other ELS variable × model adjustment × direction combination, including all Retrospective ELS analyses (both bare and adjusted) and the Prospective ELS bare model.

**Figure 6.**
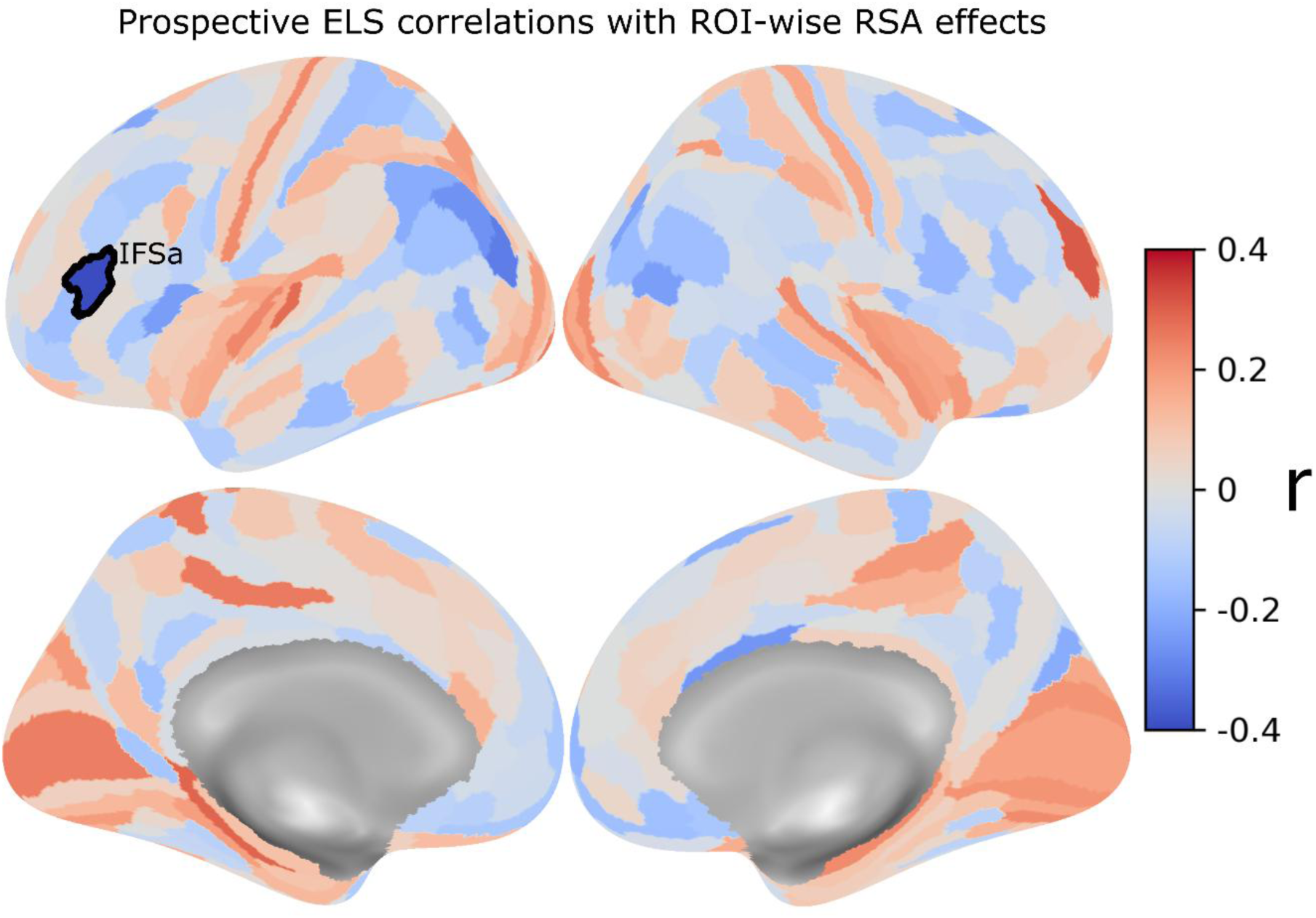
ROI-wise representational similarity analysis (RSA) of the task-versus-control model, regressed on Prospective ELS. For each HCP-MMP atlas region (360 ROIs), per-subject RDM correlation with a categorical task-versus-control model RDM was regressed on ELS controlling for sex, mother’s age, mother’s socioeconomic status, and assisted reproductive technology status. Pearson r values across subjects are displayed on the inflated fsaverage surface. Left IFSa (outlined) showed the strongest association (*r* = -0.40, FWE-corrected *p* = .019, FDR *q* = .018), indicating that higher Prospective ELS was associated with weaker categorical task-versus-control representational distinction in this region. No other ROI survived FWE or FDR correction.

#### Inter-subject Representational Similarity Analysis

IS-RSA tested whether inter-subject dissimilarity in Prospective or Retrospective ELS exposure was associated with inter-subject dissimilarity in neural representational structure in the 360 HCP-MMP parcels (Figure 7). Two models were tested for both ELS indices: matching (nearest neighbor) model that characterized pairwise dissimilarity in ELS as a pair’s absolute difference in exposure, and an additive (Anna Karenina) model that characterized the average ELS exposure of a pair. Full results for every parcel under every correction are reported in Supplementary Table S2.

**Figure 7.**
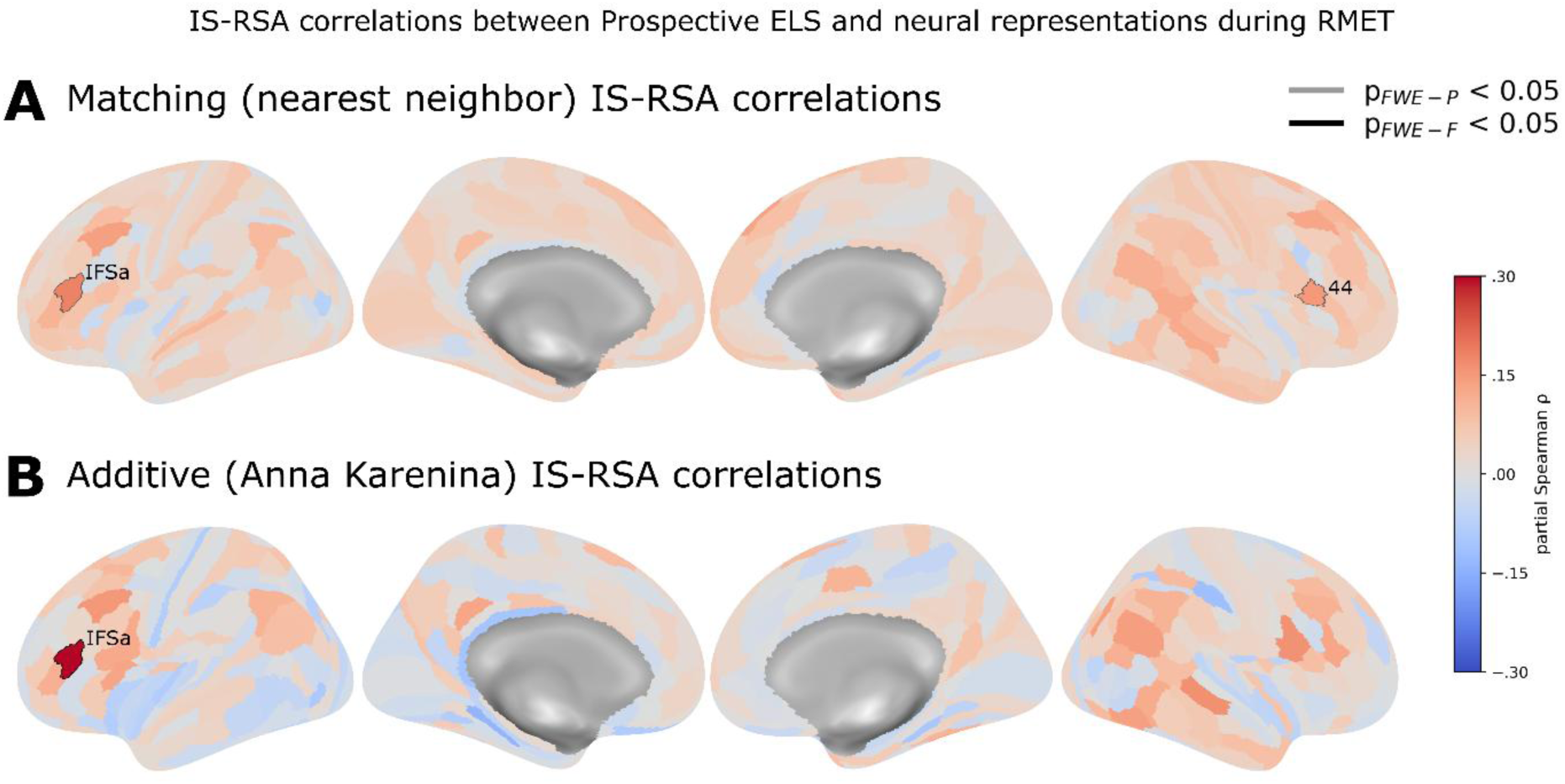
Inter-subject representational similarity analysis correlations between prospective ELS and neural representational geometry during the RMET. Correlations are partial Spearman correlations between the inter-subject neural dissimilarity matrix (ISDM) and the prospective ELS predictor ISDM, shown for all 360 HCP-MMP1.0 parcels on the inflated fsaverage surface). Neural ISDMs were computed as 1 − Spearman ρ between each pair of participants’ ROI representational dissimilarity vectors across all 3,916 participant pairs (N = 89). (A) Matching (nearest-neighbor) model, in which pairwise predictor distance is the absolute difference in ELS scores, |a − b|; positive values indicate that participants with more similar ELS exposure have more similar representational geometry. (B) Additive (Anna Karenina) model, in which pairwise predictor distance is the mean of the two participants’ ELS scores, (a + b)/2; positive values indicate that pairs higher in mean exposure are representationally more dissimilar, i.e. idiosyncrasy at high ELS and convergence at low ELS. Both models depict results for the inter-subject dissimilarity adjusted for background covariates: mother’s age and socioeconomic status, participant sex, and assisted reproductive technology status, entered as absolute pairwise differences for continuous covariates and 0/1 mismatch indicators for binary covariates. Color indicates the partial Spearman rho between rank-transformed, covariate-residualised ISDMs, scaled from −.30 to .30. Parcel outlines indicate significance under permutation testing (20,000 permutations of participant labels, with the full predictor-ISDM construction and rank-residualisation recomputed for each permutation): grey outlines denote p_FWE-P < .05, family-wise corrected across the 360 parcels within a statistic using the maximum-statistic null distribution. Black outlines denote p_FWE-F < .05, corrected across parcels and across both the matching and additive statistics. The matching statistic was tested one-tailed positive and the additive statistic two-tailed. Left anterior inferior frontal sulcus (HCPMMP 1.0 label: IFSa; index: 82) survived family-wise correction across both parcels and statistics for the additive model (ρ = .300, p_FWE-F < .05; panel B). For the matching model, the same left IFSa (*ρ* = .181, *p* FWE-P = .006, *p* FWE-F = .555) and a right IFG parcel (HCPMMP1.0 label: 44; index: 274; *ρ* = .148, *p* FWE-P = .047, *p* FWE-F = .930) survived the family-wise correction across all parcels, but not across both statistics.

For Prospective ELS, an additive effect was observed in the background covariate adjusted model for one parcel that survived correction across all parcels and both confirmatory statistics (*ρ* = .300 (*p* < .001, *p* FWE-P = .009, *p* FWE-F = .009)): left inferior frontal sulcus, anterior division (HCP-MMP label: IFSa; id: 82). This additive effect in the left IFSa was positive, meaning that participant pairs higher in mean exposure had more dissimilar representational geometries. A matching effect was also observed in IFSa, though it was weaker than the additive effect and didn’t survive the strictest family-wise correction (*ρ* = .181; *p* < .001, *p* FWE-P = .006, *p* FWE-F = .555). The combined model separated the two components and confirmed that most of the effect was additive: *β(additive)* = .269 (*p* = .002) against *β(matching)* = .097 (*p* = .045). This separation was robust to background covariate adjustments: *β(additive)* = .265, *p* = .003; *β(matching)* = .095, *p* = .052).

Right inferior frontal gyrus (HCP-MMP label: 44; id: 274) showed a matching effect only: *ρ* = .148 (*p* = .001, *p* FWE-P = .047, *p* FWE-F = .930) in the adjusted model, with *β(matching)* = .134 (*p* = .004) and no additive component (*ρ* = .088, *p* = .292; *β(additive)* = .046, *p* = .591). However, this effect survived correction only across parcels within the matching statistic, but not correction across the full confirmatory family (all parcels and both statistics).

Retrospective ELS produced no parcel surviving correction under either statistic in either model. The strongest parcel-wise effects were far from corrected significance (minimum *p* FWE-P = .616 in the adjusted model). No parcel in any model survived Benjamini–Hochberg false discovery rate (FDR) correction at *q* < .05. The smallest FDR obtained was *q* = .072 for the matching statistic in left IFSa in the unadjusted Prospective ELS model (*q* = .108 in the adjusted model).

#### Post hoc characterization of representational geometry

The following analyses are descriptive characterizations of the IS-RSA result in left IFSa. They were performed after inference was complete, involve no additional hypothesis test, and are reported to describe what the effect looks like at the level of the RDM.

To characterize and visualize how neural representations during the RMET changed as a function of Prospective ELS, a two-dimensional multidimensional scaling (MDS) embedding was used alongside Procrustes alignment (Figure 8A). The MDS embeddings captured a median of 27.0% of the positive eigenvalue mass of individual subjects’ RDMs (range 16.7–45.1%), and 17.8% of the group-mean RDM. Individual RDMs were thus somewhat more compressible in two dimensions than the group average, consistent with each subject’s leading dimensions being partly subject-specific. Alignment with Generalized Procrustes Analysis (GPA) converged in 10 iterations. It applied a median rotation of 118.2° and required a reflection in 48 of 89 subjects, confirming that the raw embeddings carried no common orientation and could not have been compared without alignment. Alignment reduced the mean distance between individual configurations, and the group mean by 11.5% (3.894 to 3.446).

**Figure 8.**
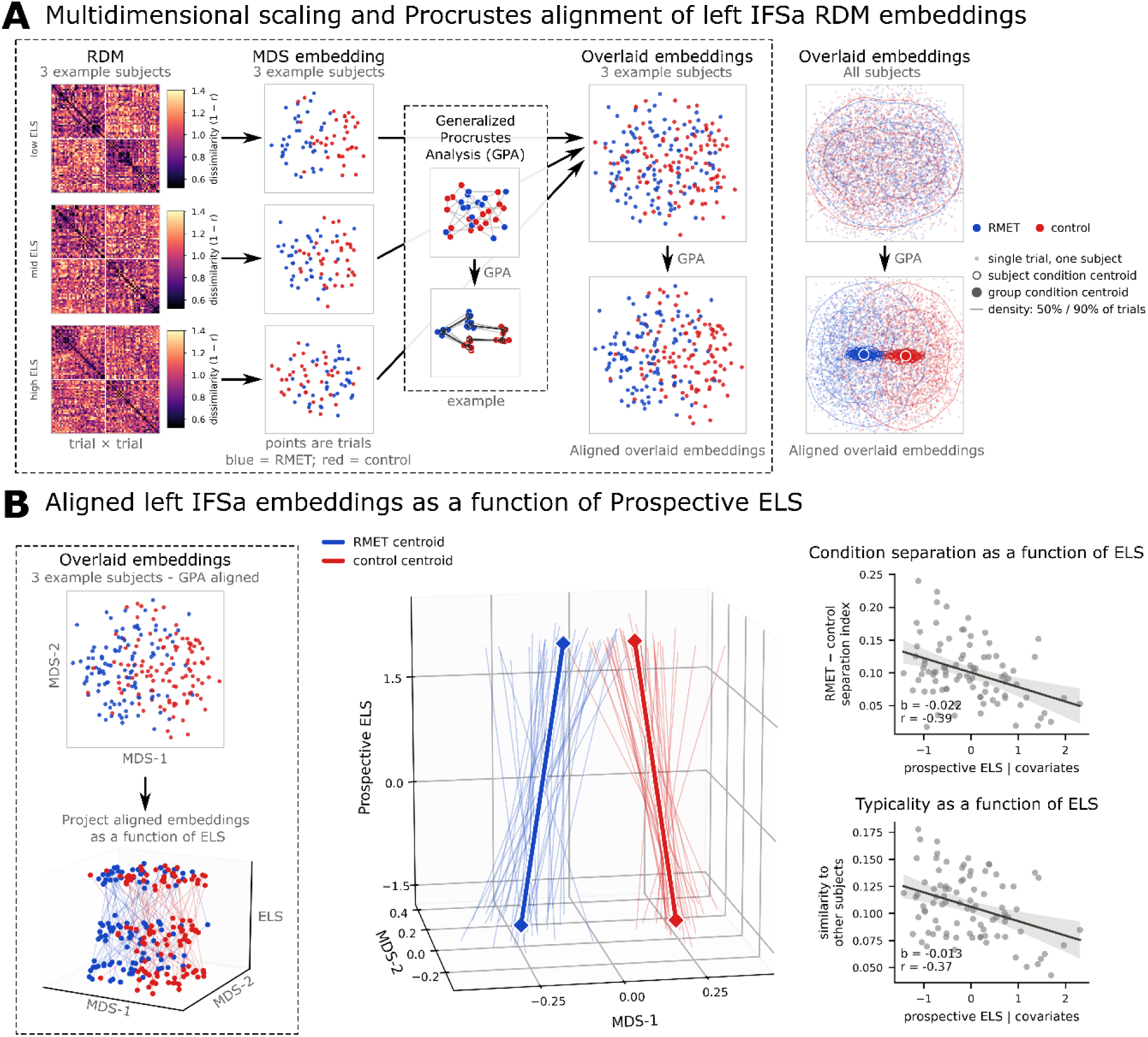
Multidimensional scaling and Procrustes alignment of left IFSa representational geometry as a function of Prospective ELS. (A) Pipeline for aligning representational dissimilarity matrix (RDM) embeddings. Left: trial-by-trial RDMs (72 × 72; 1 − Pearson r between beta-series patterns) for three example subjects at low, intermediate and high Prospective ELS. Trials are ordered RMET then control; color indicates dissimilarity. Each RDM was embedded in two dimensions by classical multidimensional scaling (MDS). Each point is one trial (blue, RMET; red, control). Because MDS recovers a configuration only up to rotation and reflection, the axes of these embeddings carry no shared meaning across subjects, and the solutions are not directly comparable. Generalized Procrustes analysis (GPA) resolves this by iteratively rotating and reflecting each configuration onto the group mean configuration; scaling is withheld, so configuration size is preserved. The three example subject’s MDS embeddings are then shown overlaid before (top) and after (bottom) alignment. Right: all 89 subjects overlaid, before (top) and after (bottom) GPA. Small dots are single trials in single subjects; open rings are individual subjects’ condition centroids; filled markers are the group condition centroids; contours enclose 50% and 90% of each condition’s trials. (B) Aligned embeddings stacked along Prospective ELS. Left: the three example subjects’ aligned configurations, then the same configurations positioned along a third axis given by prospective ELS. Centre: all subjects. Thin lines show, for each trial, the linear fit of its aligned coordinate on prospective ELS across subjects; thick lines show the corresponding fits for the RMET and control condition centroids, with diamonds marking the fitted positions at the lowest and highest observed ELS. The fits are across subjects, not within-subject trajectories. The centroid trajectories converge as Prospective ELS increases, indicating less sharply differentiated task and control representations at higher levels of exposure. Right: the two quantities describing change in representational geometry between subjects, each computed on the full 72 × 72 RDMs and adjusted for background covariates (participant sex, mother’s age, mother’s socioeconomic status, and assisted reproductive technology status). Top: condition separation, the mean between-condition dissimilarity minus the mean within-condition dissimilarity, scaled by the overall mean dissimilarity of that subject’s RDM (partial r = −.39). Bottom: typicality, one minus each subject’s mean correlation distance to all other subjects’ RDMs, so that higher values indicate a geometry closer to the group norm (partial r = −.37). One point per subject; shading is the 95% confidence band. Both quantities decrease with Prospective ELS: representations of the two conditions become less well separated, and individual geometries less typical of the group. Two-dimensional embeddings are shown for visualization only and capture a median of 27% of each subject’s representational geometry; all inferential analyses used the full RDMs. Separation measured in the aligned two-dimensional frame agreed closely with the same quantity computed on the full RDMs (r = .93), which supports the stacked projection as an illustration of the full-RDM result.

Separation between mentalizing and control representations decreased with Prospective ELS. Computed on the full RDMs and adjusted for background covariates, the association was partial r = −.393 (b = −.0221, p < .001, N = 89). The equivalent quantity measured in the aligned two-dimensional MDS embedding frame was partial r = −.368 (b = −.0678, p < .001), and the two measures agreed closely across subjects (r = .934). The convergence of the condition centroid trajectories in Figure 8B therefore reflects a property of the full representational geometry rather than an artefact of the projection.

Typicality, the proximity of a subject’s RDM to the group average, also decreased with Prospective ELS (partial r = −.378, p < .001). Equivalently, mean inter-subject dissimilarity increased across the same range. Mean inter-subject correlation distance in left IFSa was 0.893 (SD = 0.050), indicating that subjects shared relatively little representational structure in this parcel overall. Higher-ELS subjects departed further from the group norm than lower-ELS subjects, in addition to differentiating the two conditions less sharply.

## Discussion

Mentalizing, the ability to attribute mental states to others by observing their behavior, is a socially central and developmentally susceptible human capacity. In the present longitudinal study, we explored whether and how early life stress (ELS) was associated with mentalizing during Reading the Mind in the Eyes task (RMET), both in terms of behavioral performance, neural activation, and neural representations in a sample of young adults with no psychiatric diagnoses. Two qualitatively different operationalizations of ELS were employed separately across the analyses: Prospective ELS, measured across three time points as parental self-reported mental health and family relationship problems in pregnancy and infancy (when the child was 2 and 12 months old), and Retrospective ELS, measured as participant adverse childhood experiences self-reported during late adolescence.

Behaviorally, we observed a small positive association between mentalizing performance and Prospective ELS exposure. That is, participants with higher Prospective ELS exposure performed slightly better in the task than participants with lower exposure. The association between Retrospective ELS and mentalizing performance was also positive but did not reach statistical significance. While a decremental influence of ELS on social cognitive task performance is expected due to being observed in many previous studies (Mackey et al, 2025), our behavioral results did not replicate these findings. Our findings instead align with the adaptive calibration framework, among other similar narratives, that suggest that ELS could also influence context-sensitive specialization in social cue processing (Bérubé et al., 2023; Young et al., 2022; Ellis and Del Giudice, 2019; Frankenhuis and de Weerth, 2013; Del Giudice et al., 2011). Early social environments that place more social-emotional demands on the child might, despite being potentially detrimental for optimal psychological development, also predispose children to adaptive demands that result in increased sensitivity to perceiving and inferring social-emotional nuances.

Neurally, across participants, both univariate activation and searchlight representational similarity analysis (RSA) converged on a similar set of regions consistent with the canonical mentalizing network (Maliske et al., 2023; Schurz et al., 2021; Molenberghs et al., 2016; Schurz et al., 2014; Mar, 2011). Importantly for the present aims, the agreement between mean-activation and searchlight RSA supports the inference that the regions within the mentalizing network are not merely more active during mentalizing but also carry representational information that distinguishes mentalizing from non-mentalizing cognitive processing.

Between subjects, Prospective ELS was associated with how the left anterior inferior frontal sulcus (IFSa), a region bordering, and partly overlapping, Broca’s area, represented the mentalizing task. Two findings converge in the left IFSa. First, participants with higher Prospective ELS showed less separation between the mentalizing and control conditions, meaning that their neural response patterns during mentalizing and the control task resembled each other more closely. Second, inter-subject analysis showed that participants with higher Prospective ELS also resembled each other less: representations became increasingly individual as exposure rose, while participants at the low end of ELS shared a more common neural response pattern. This asymmetry is sometimes called an Anna Karenina (AK) effect, where the less ELS exposed share more similar neural response patterns, while those more ELS exposed become increasingly idiosyncratic (Finn et al., 2020).

A second but statistically weaker pattern appeared in IFSa and in the right inferior frontal gyrus (Brodmann area 44): participants with similar levels of Prospective ELS exposure had more similar neural representations, irrespective of exposure level (also known as a nearest neighbor (NN) effect). These two patterns make different claims: one (AK) considers the average level of exposure of a pair, the other (NN) the “match” between a pair. Because these two effects were partly correlated in our inter-subject data, we estimated each while holding the other constant. In left IFSa the average ELS exposure level-based AK effect held while the matching NN effect fell. In the right inferior frontal gyrus, the reverse was true: the matching NN effect was essentially unchanged while holding the AK effect constant. However, only the left IFSa AK result survived our strictest correction, and we treat the right inferior frontal gyrus finding as provisional.

These results bear on the long-standing question of what the RMET measures (Oakley et al., 2016; Peterson & Miller, 2012). Each trial requires the selection of a mental state word among four options that best matches an image of a person’s eyes, imposing lexical-semantic and semantic-control demands alongside mentalizing itself. Both regions carrying our ELS-related effects lie in the inferior frontal cortex. Left IFSa lies at least partly within the classical Broca’s territory, and area 44 can be considered its right-hemisphere homolog. Both regions are thus implicated in frontal contributions to mentalizing but also in semantic control (Diveica et al., 2021; Balgová et al., 2024; Turker et al., 2025). Additionally, even lesions of the left IFG have been shown to hinder RMET performance specifically (Dal Monte et al., 2014). That the task engaged the broad mentalizing network while ELS-related individual differences localized specifically to these inferior frontal cortex nodes suggests that ELS might modulate the semantic and verbal-evaluative component of mentalizing. Prospective ELS was, however, also associated with higher accuracy, which suggests the observed ELS related shifts in neural representations in the inferior frontal cortex might be a form of adaptive calibration (Frankenhuis & de Weerth, 2013). Thus, how far the development of mentalizing can be sensibly dissociated from development of semantic processing is a relevant question to untangle in future research.

Some noteworthy implications and limitations of our study are warranted. First, our neural findings relate to Prospective ELS and not to Retrospective ELS, and the two measures were uncorrelated in the sample. Given this low agreement, the two instruments identify substantially non-overlapping groups of participants, consistent with the broader literature on prospective vs. retrospective accounts of childhood adversity (Baldwin et al., 2019; Danese & Widom, 2020; Reuben et al., 2016). Two differences between the measures are relevant to interpreting this dissociation. First, the prospective measure records the caregiving environment as reported by parents at the time, whereas the retrospective measure records what the participant later recalls and appraises as adverse. Second, our prospective measure covers pregnancy and the first year of life, a period no participant can report on, while the retrospective measure spans childhood and adolescence. The dissociation may therefore reflect developmental timing or measurement method, and the present design cannot separate these two. Second, cumulative risk scores such as our Prospective ELS index sum qualitatively different circumstances (e.g., parental depressive symptoms, relationship conflict) into a single quantity. Two participants with the same score might still have nontrivially dissimilar early environments, and this potential mismatch grows with the score, as there are few ways to score low and many ways to score high. Increasing representational idiosyncrasy with increasing exposure could therefore reflect increasingly heterogeneous inputs rather than a single disorganizing process acting on the individual. Third, inclusion criteria for the participants included the lack of psychiatric diagnoses. Because ELS is a known predictor of future psychiatric problems (McLaughlin, 2020), it is conceivable that the influence of ELS on behavioral or neural responses to social cognitive tasks relies at least partly on the same factors that dictate whether ELS leads to psychiatric problems. Including only individuals with no psychiatric diagnoses might thus bias the sample towards those more resilient to the psychopathological influences of ELS. Future research should aim to elucidate how the presence of psychiatric symptoms or diagnoses, or lack thereof, relates to the influence of varying levels of ELS on behavioral and neural responses to different social cognitive tasks.

Finally, neural data were derived from a single imaging session, limiting the establishment of causal developmental trajectories of ELS as well as multi-run fMRI approaches (e.g. the use of Crossnobis distances) in RSA. The sample size, while adequate for the within-subjects effects, places limits on the precision of individual-differences and inter-subject estimates, and the regional and hemispheric specificity reported above should be treated as exploratory and hypothesis-generating. This and the above considerations point to a clearer direction for future work. Tasks that dissociate verbal-semantic from mentalizing demands, such as nonverbal or implicit mentalizing paradigms, false-belief tasks with matched semantic load, or animation-based stimuli, would allow a more direct test of whether the inferior frontal ELS effect reflects modulation of semantic control, of mentalizing, or of their interaction. Pairing such paradigms with the representational and inter-subject approaches employed here would help determine whether ELS exerts a general influence on controlled, verbally mediated social evaluation or a more specific influence on mental-state representations.

In sum, the present study sought to characterize the development of mentalizing related behavioral performance, brain activity, and neural representations, using two approaches to operationalizing early life stress. The results offer two key contributions. First, they demonstrate that the neural representation of mentalizing can be systematically related to problems in prospectively measured early parental mental health and relationship problems. Second, they highlight the inferior frontal cortex as a relevant target of mentalizing related functional vulnerability to ELS. The study also contributes to further theoretical and empirical work on the nature of mentalizing itself and highlights the need for an experimental disentanglement of the lexical-semantic and semantic-control components from other mentalizing related processes.

## Data availability

The raw data from the ongoing longitudinal study project are not readily available because participant privacy and ethical permissions do not allow public sharing of the data. Requests to access the datasets should be directed to.

## Author contributions

MI: Conceptualization, methodology, software, validation, formal analysis, data curation, writing – original draft, writing – review & editing, visualization, funding acquisition. JL: Conceptualization, investigation, data curation, writing – review & editing, supervision, project administration, funding acquisition. MF: investigation, data curation, writing – review & editing. MV: investigation, data curation, writing – review & editing. PW: conceptualization, methodology, software, validation, investigation, data curation, writing – review & editing, supervision, project administration.

## Acknowledgements

This study was a part of the Miracles of Development research project supported by the Academy of Finland (#3266413), and an individual grant from the Academy of Finland for JL (#323845). MI has received individual grants from the Finnish Brain Foundation, Signe and Ane Gyllenberg Foundation, Finnish Cultural Foundation, and salaried doctoral position in the Doctoral programme in Human Behavior of the University of Helsinki.

## References

Abidin, R. R. (1997). Parenting Stress Index: A measure of the parent–child system. In C. P. Zalaquett & R. J. Wood (Eds.), Evaluating stress: A book of resources (pp. 277–291). Scarecrow Education.

Allen, J. G., Fonagy, P., & Bateman, A. W. (2008). Mentalizing in clinical practice. American Psychiatric Publishing.

Aquino, K. M., Fulcher, B. D., Parkes, L., Sabaroedin, K., & Fornito, A. (2020). Identifying and removing widespread signal deflections from fMRI data: Rethinking the global signal regression problem. NeuroImage, 212, 116614. 10.1016/j.neuroimage.2020.116614

Arabadzhiev, Z. & Paunova, R. (2024). Complexity of mentalization. Frontiers in Psychology, 15:1353804. 10.3389/fpsyg.2024.1353804

Avants, B. B., Epstein, C. L., Grossman, M., & Gee, J. C. (2008). Symmetric diffeomorphic image registration with cross-correlation: evaluating automated labeling of elderly and neurodegenerative brain. Medical image analysis, 12(1), 26–41. 10.1016/j.media.2007.06.004

Baldwin, J. R., Reuben, A., Newbury, J. B., & Danese, A. (2019). Agreement Between Prospective and Retrospective Measures of Childhood Maltreatment: A Systematic Review and Meta-analysis. JAMA psychiatry, 76(6), 584–593. 10.1001/jamapsychiatry.2019.0097

Balgová, E., Diveica, V., Jackson, R. L., & Binney, R. J. (2024). Overlapping neural correlates underpin theory of mind and semantic cognition: Evidence from a meta-analysis of 344 functional neuroimaging studies. Neuropsychologia, 200, 108904. 10.1016/j.neuropsychologia.2024.108904

Baron-Cohen, S., Wheelwright, S., Hill, J., Raste, Y., & Plumb, I. (2001). The “Reading the mind in the eyes” Test revised version: A study with normal adults, and adults with Asperger syndrome or high-functioning autism. Journal of Child Psychology and Psychiatry, 42(2), 241–251. 10.1111/1469-7610.00715

Beck, A. T., Ward, C. H., Mendelson, M., Mock, J., & Erbaugh, J. (1961). An Inventory for Measuring Depression. Archives of General Psychiatry, 4: 561–571. 10.1001/archpsyc.1961.01710120031004

Behzadi, Y., Restom, K., Liau, J., & Liu, T. T. (2007). A component based noise correction method (CompCor) for BOLD and perfusion based fMRI. NeuroImage, 37(1), 90–101. 10.1016/j.neuroimage.2007.04.042

Bérubé, A., Turgeon, J., Blais, C., & Fiset, D. (2023). Emotion Recognition in Adults With a History of Childhood Maltreatment: A Systematic Review. Trauma, violence & abuse, 24(1), 278–294. 10.1177/15248380211029403

Buergi, N., Aydogan, G., Konovalov, A., & Ruff, C. C. (2026). A neural signature of adaptive mentalization. Nature Neuroscience, 1–11. 10.1038/s41593-026-02219-x

Cox, R. W., & Hyde, J. S. (1997). Software tools for analysis and visualization of fMRI data. NMR in biomedicine, 10(4-5), 171–178. 10.1002/(sici)1099-1492(199706/08)10:4/5<171::aid-nbm453>3.0.co;2-l

Cracco, E., Hudson, A. R., Van Hamme, C., Maeyens, L., Brass, M., & Mueller, S. C. (2020). Early interpersonal trauma reduces temporoparietal junction activity during spontaneous mentalising. Social Cognitive and Affective Neuroscience, 15(1), 12–22. 10.1093/scan/nsaa015

Dal Monte, O., Schintu, S., Pardini, M., Berti, A., Wassermann, E. M., Grafman, J., & Krueger, F. (2014). The left inferior frontal gyrus is crucial for reading the mind in the eyes: brain lesion evidence. Cortex; a journal devoted to the study of the nervous system and behavior, 58, 9–17. 10.1016/j.cortex.2014.05.002

Dale, A. M., Fischl, B., & Sereno, M. I. (1999). Cortical surface-based analysis. I. Segmentation and surface reconstruction. NeuroImage, 9(2), 179–194. 10.1006/nimg.1998.0395

Danese, A., & Widom, C. S. (2020). Objective and subjective experiences of child maltreatment and their relationships with psychopathology. Nature Human Behaviour, 4(8), 811–818. 10.1038/s41562-020-0880-3

Del Giudice, M., Ellis, B. J., & Shirtcliff, E. A. (2011). The Adaptive Calibration Model of stress responsivity. Neuroscience and Biobehavioral Reviews, 35(7), 1562–1592. 10.1016/j.neubiorev.2010.11.007

Diedrichsen, J., & Kriegeskorte, N. (2017). Representational models: A common framework for understanding encoding, pattern-component, and representational-similarity analysis. PLoS computational biology, 13(4), e1005508. 10.1371/journal.pcbi.1005508

Diveica, V., Koldewyn, K., & Binney, R. J. (2021). Establishing a role of the semantic control network in social cognitive processing: A meta-analysis of functional neuroimaging studies. NeuroImage, 245, 118702. 10.1016/j.neuroimage.2021.118702

Ellis, B. J., & Del Giudice, M. (2019). Developmental Adaptation to Stress: An Evolutionary Perspective. Annual Review of Psychology, 70, 111–139. 10.1146/annurev-psych-122216-011732

Esteban, O., Markiewicz, C. J., Blair, R. W., Moodie, C. A., Isik, A. I., Erramuzpe, A., Kent, J. D., Goncalves, M., DuPre, E., Snyder, M., Oya, H., Ghosh, S. S., Wright, J., Durnez, J., Poldrack, R. A., & Gorgolewski, K. J. (2019). fMRIPrep: a robust preprocessing pipeline for functional MRI. Nature methods, 16(1), 111–116. 10.1038/s41592-018-0235-4

Ettekal, I., Eiden, R. D., Nickerson, A. B., & Schuetze, P. (2019). Comparing alternative methods of measuring cumulative risk based on multiple risk indicators: Are there differential effects on children’s externalizing problems?. PloS one, 14(7), e0219134. 10.1371/journal.pone.0219134

Finkelhor, D., Shattuck, A., Turner, H., & Hamby, S. (2015). A revised inventory of Adverse Childhood Experiences. Child abuse & neglect, 48, 13–21. 10.1016/j.chiabu.2015.07.011

Finn, E. S., Glerean, E., Khojandi, A. Y., Nielson, D., Molfese, P. J., Handwerker, D. A., & Bandettini, P. A. (2020). Idiosynchrony: From shared responses to individual differences during naturalistic neuroimaging. NeuroImage, 215, 116828. 10.1016/j.neuroimage.2020.116828

Flykt, M., Vänskä, M., Punamäki, R. L., Heikkilä, L., Tiitinen, A., Poikkeus, P., & Lindblom, J. (2021). Adolescent Attachment Profiles Are Associated With Mental Health and Risk-Taking Behavior. Frontiers in Psychology, 12, 761864. 10.3389/fpsyg.2021.761864

Frankenhuis, W. E., & de Weerth, C. (2013). Does early-life exposure to stress shape or impair cognition? Current Directions in Psychological Science, 22(5), 407–412. 10.1177/0963721413484324

Freeman, J. B., Stolier, R. M., Brooks, J. A., & Stillerman, B. S. (2018). The neural representational geometry of social perception. Current Opinion in Psychology, 24, 83–91. 10.1016/j.copsyc.2018.10.003

Fonagy, P. (1991). Thinking about thinking: Some clinical and theoretical considerations in the treatment of a borderline patient. The International Journal of Psychoanalysis, 72(4), 639–656.

Fonagy, P., & Allison, E. (2012). What is mentalization? The concept and its foundations in developmental research. In N. Midgley & I. Vrouva (Eds.), Minding the child: Mentalization-based interventions with children, young people and their families (pp. 11–34). Routledge/Taylor & Francis Group.

Frith, C. D., & Frith, U. (2006). The neural basis of mentalizing. Neuron, 50(4), 531–534. 10.1016/j.neuron.2006.05.001

Frost, M. A., & Goebel, R. (2012). Measuring structural-functional correspondence: spatial variability of specialised brain regions after macro-anatomical alignment. NeuroImage, 59(2), 1369–1381. 10.1016/j.neuroimage.2011.08.035

Gergely, G., & Watson, J. (1999). Early social-emotional development: Contingency perception and the social biofeedback model. In P. Rochat (Ed.), Early social cognition: Understanding others in the first months of life (pp. 101–137). Hillsdale, NJ: Erlbaum.

Glasser, M. F., Coalson, T. S., Robinson, E. C., Hacker, C. D., Harwell, J., Yacoub, E., Ugurbil, K., Andersson, J., Beckmann, C. F., Jenkinson, M., Smith, S. M., & Van Essen, D. C. (2016). A multi-modal parcellation of human cerebral cortex. Nature, 536(7615), 171–178. 10.1038/nature18933

Goldberg, D. P., & Hillier, V. F. (1979). A scaled version of the General Health Questionnaire. Psychological medicine, 9(1), 139–145. 10.1017/s0033291700021644

Golec-Staśkiewicz, K., Pluta, A., Wojciechowski, J., Okruszek, Ł., Haman, M., Wysocka, J., & Wolak, T. (2022). Does the TPJ fit it all? Representational similarity analysis of different forms of mentalizing. Social Neuroscience, 17(5), 428–440. 10.1080/17470919.2022.2138536

Gorgellino, M., Kumar, G., Parkar, Y., Catalan, A., Fares-Otero, N., Debbané, M., Armando, M., & Alameda, L. (2025). The shadow of trauma: impaired mentalization in clinical populations -a systematic review. Psychological Medicine, 55, e186. 10.1017/S0033291725100822

Greve, D. N., & Fischl, B. (2009). Accurate and robust brain image alignment using boundary-based registration. NeuroImage, 48(1), 63–72. 10.1016/j.neuroimage.2009.06.060

Greve, D. N., & Fischl, B. (2018). False positive rates in surface-based anatomical analysis. NeuroImage, 171, 6–14. 10.1016/j.neuroimage.2017.12.072

Hafner, R. M., Johnson, B. N., Tone, E. B., Kivity, Y., Levy, K. N., & Bedwell, J. S. (2026). A multi-site psychometric evaluation of the Reading the Mind in the Eyes Test–Revised. Personality and Individual Differences, 250, 113526. 10.1016/j.paid.2025.113526

Hagler, D. J., Jr, Saygin, A. P., & Sereno, M. I. (2006). Smoothing and cluster thresholding for cortical surface-based group analysis of fMRI data. NeuroImage, 33(4), 1093–1103. 10.1016/j.neuroimage.2006.07.036

Higgins, W. C., Ross, R. M., Langdon, R., & Polito, V. (2023). The “Reading the Mind in the Eyes” Test Shows Poor Psychometric Properties in a Large, Demographically Representative U.S. Sample. Assessment, 30(6), 1777–1789. 10.1177/10731911221124342

Hughes, C., Jaffee, S. R., Happé, F., Taylor, A., Caspi, A., & Moffitt, T. E. (2005). Origins of Individual Differences in Theory of Mind: From Nature to Nurture? Child Development, 76(2), 356–370. 10.1111/j.1467-8624.2005.00850_a.x

Huntenburg, J. M. (2014). Evaluating nonlinear coregistration of BOLD EPI and T1w images [Master’s thesis, Freie Universität Berlin]. Max Planck Society.

Ilomäki, M., Lindblom, J., Salmela, V., Flykt, M., Vänskä, M., Salmi, J., Tolonen, T., Alho, K., Punamäki, R. L., & Wikman, P. (2022). Early life stress is associated with the default mode and fronto-limbic network connectivity among young adults. Frontiers in Behavioral Neuroscience, 16, 958580. 10.3389/fnbeh.2022.958580

Ilomäki, M., Lindblom, J., Flykt, M., Vänskä, M., Punamäki, R. L., & Wikman, P. (2025). Similarity in Early Life Stress Exposure Is Associated With Similarity in Neural Representations in Early Adulthood. Human Brain Mapping, 46(14), e70373. 10.1002/hbm.70373

Jenkinson, M., Bannister, P., Brady, M., & Smith, S. (2002). Improved optimization for the robust and accurate linear registration and motion correction of brain images. NeuroImage, 17(2), 825– 841. 10.1016/s1053-8119(02)91132-8

Kim, S. (2015). The mind in the making: Developmental and neurobiological origins of mentalizing. Personality Disorders: Theory, Research, and Treatment, 6(4), 356–365. 10.1037/per0000102

Koster-Hale, J., Richardson, H., Velez, N., Asaba, M., Young, L., & Saxe, R. (2017). Mentalizing regions represent distributed, continuous, and abstract dimensions of others’ beliefs. NeuroImage, 161, 9–18. 10.1016/j.neuroimage.2017.08.026

Kriegeskorte, N., Mur, M., & Bandettini, P. (2008). Representational similarity analysis - connecting the branches of systems neuroscience. Frontiers in Systems Neuroscience, 2, 4. 10.3389/neuro.06.004.2008

Lanczos, C. (1964). Evaluation of Noisy Data. Journal of the Society for Industrial and Applied Mathematics, Series B: Numerical Analysis, 1(1), 76–85. 10.1137/0701007

Mackey, M., Dunne, E., & Ahern, E. (2025). Early Life Adversity and Social Cognition in the General Adult Population: A Systematic Review and Meta-Analysis. Journal of Child & Adolescent Trauma, 1–21. 10.1007/s40653-025-00724-y

Maliske, L. Z., Schurz, M., & Kanske, P. (2023). Interactions within the social brain: Co-activation and connectivity among networks enabling empathy and Theory of Mind. Neuroscience and biobehavioral reviews, 147, 105080. 10.1016/j.neubiorev.2023.105080

Mar, R. A. (2011). The neural bases of social cognition and story comprehension. Annual Review of Psychology, 62, 103–134. 10.1146/annurev-psych-120709-145406

Martin-Gagnon, G., Normandin, L., Fonagy, P., & Ensink, K. (2023). Adolescent mentalizing and childhood emotional abuse: implications for depression, anxiety, and borderline personality disorder features. Frontiers in Psychology, 14, 1237735. 10.3389/fpsyg.2023.1237735

McLaughlin, K. A. (2020). Early life stress and psychopathology. In K. L. Harkness & E. P. Hayden (Eds.), The Oxford handbook of stress and mental health (pp. 45–74). Oxford University Press. 10.1093/oxfordhb/9780190681777.013.3

Molenberghs, P., Johnson, H., Henry, J. D., & Mattingley, J. B. (2016). Understanding the minds of others: A neuroimaging meta-analysis. Neuroscience & Biobehavioral Reviews, 65, 276–291. 10.1016/j.neubiorev.2016.03.020

Moor, B. G., Macks, Z. A., Güroglu, B., Rombouts, S. A., Molen, M. W., & Crone, E. A. (2012). Neurodevelopmental changes of reading the mind in the eyes. Social Cognitive and Affective Neuroscience, 7(1), 44–52. 10.1093/scan/nsr020

Mumford J. A. (2012). A power calculation guide for fMRI studies. Social cognitive and affective neuroscience, 7(6), 738–742. 10.1093/scan/nss059

Nili, H., Wingfield, C., Walther, A., Su, L., Marslen-Wilson, W., & Kriegeskorte, N. (2014). A toolbox for representational similarity analysis. PLoS Computational Biology, 10(4), e1003553. 10.1371/journal.pcbi.1003553

Nolte, T., Bolling, D. Z., Hudac, C. M., Fonagy, P., Mayes, L., & Pelphrey, K. A. (2013). Brain mechanisms underlying the impact of attachment-related stress on social cognition. Frontiers in Human Neuroscience, 7, 816. 10.3389/fnhum.2013.00816

Oakley, B. F. M., Brewer, R., Bird, G., & Catmur, C. (2016). Theory of mind is not theory of emotion: A cautionary note on the Reading the Mind in the Eyes Test. Journal of abnormal psychology, 125(6), 818–823. 10.1037/abn0000182

Olderbak, S., Wilhelm, O., Olaru, G., Geiger, M., Brenneman, M. W., & Roberts, R. D. (2015). A psychometric analysis of the reading the mind in the eyes test: toward a brief form for research and applied settings. Frontiers in Psychology, 6, 1503. 10.3389/fpsyg.2015.01503

Parsons, V. L. 2017. “Stratified Sampling.” In Wiley Statsref: Statistics Reference Online, edited by Balakrishnan N., Colton T., Everitt B., Piegorsch W., Ruggeri F., and Teugels J. L.. John Wiley & Sons. 10.1002/9781118445112.stat05999.pub2

Pechtel, P., & Pizzagalli, D. A. (2011). Effects of early life stress on cognitive and affective function: an integrated review of human literature. Psychopharmacology, 214(1), 55–70. 10.1007/s00213-010-2009-2

Peterson, E., & Miller, S. F. (2012). The eyes test as a measure of individual differences: how much of the variance reflects verbal IQ?. Frontiers in Psychology, 3, 220. 10.3389/fpsyg.2012.00220

Power, J. D., Mitra, A., Laumann, T. O., Snyder, A. Z., Schlaggar, B. L., & Petersen, S. E. (2014). Methods to detect, characterize, and remove motion artifact in resting state fMRI. NeuroImage, 84, 320–341. 10.1016/j.neuroimage.2013.08.048

Premack, D., & Woodruff, G. (1978). Does the chimpanzee have a theory of mind? Behavioral and Brain Sciences, 1(4), 515–526. 10.1017/S0140525X00076512

Quesque, F., Apperly, I., Baillargeon, R., Baron-Cohen, S., Becchio, C., Bekkering, H., … & Brass, M. (2024). Defining key concepts for mental state attribution. Communications Psychology, 2(1), 29. 10.1038/s44271-024-00077-6

Reuben, A., Moffitt, T. E., Caspi, A., Belsky, D. W., Harrington, H., Schroeder, F., Hogan, S., Ramrakha, S., Poulton, R., & Danese, A. (2016). Lest we forget: comparing retrospective and prospective assessments of adverse childhood experiences in the prediction of adult health. Journal of Child Psychology and Psychiatry, and Allied Disciplines, 57(10), 1103–1112. 10.1111/jcpp.12621

Rokita, K. I., Dauvermann, M. R., & Donohoe, G. (2018). Early life experiences and social cognition in major psychiatric disorders: a systematic review. European psychiatry, 53, 123–133. 10.1016/j.eurpsy.2018.06.006

Schurz, M., Radua, J., Aichhorn, M., Richlan, F., & Perner, J. (2014). Fractionating theory of mind: a meta-analysis of functional brain imaging studies. Neuroscience and biobehavioral reviews, 42, 9–34. 10.1016/j.neubiorev.2014.01.009

Schurz, M., Radua, J., Tholen, M. G., Maliske, L., Margulies, D. S., Mars, R. B., Sallet, J., & Kanske, P. (2021). Toward a hierarchical model of social cognition: A neuroimaging meta-analysis and integrative review of empathy and theory of mind. Psychological bulletin, 147(3), 293–327. 10.1037/bul0000303

Smith, S. M., & Nichols, T. E. (2009). Threshold-free cluster enhancement: addressing problems of smoothing, threshold dependence and localisation in cluster inference. NeuroImage, 44(1), 83–98. 10.1016/j.neuroimage.2008.03.061

Spanier, G. B. (1976). Measuring dyadic adjustment: New scales for assessing the quality of marriage and similar dyads. Journal of Marriage and the Family, 38(1), 15–28. 10.2307/350547

Thornton, M. A., & Mitchell, J. P. (2018). Theories of Person Perception Predict Patterns of Neural Activity During Mentalizing. Cerebral Cortex (New York, N.Y. : 1991), 28(10), 3505–3520. 10.1093/cercor/bhx216

Thornton, M. A., Weaverdyck, M. E., & Tamir, D. I. (2019). The brain represents people as the mental states they habitually experience. Nature Communications, 10(1), 2291. 10.1038/s41467-019-10309-7

Treiber, J. M., White, N. S., Steed, T. C., Bartsch, H., Holland, D., Farid, N., McDonald, C. R., Carter, B. S., Dale, A. M., & Chen, C. C. (2016). Characterization and Correction of Geometric Distortions in 814 Diffusion Weighted Images. PloS one, 11(3), e0152472. 10.1371/journal.pone.0152472

Trujillo-Llano, C., Sainz-Ballesteros, A., Suarez-Ardila, F., Gonzalez-Gadea, M. L., Ibáñez, A., Herrera, E., & Baez, S. (2024). Neuroanatomical markers of social cognition in neglected adolescents. Neurobiology of Stress, 31, 100642. 10.1016/j.ynstr.2024.100642

Turker, S., Fumagalli, B., Kuhnke, P., & Hartwigsen, G. (2025). The ‘reading’brain: Meta-analytic insight into functional activation during reading in adults. Neuroscience & Biobehavioral Reviews, 173, 106166. 10.1016/j.neubiorev.2025.106166

Tustison, N. J., Avants, B. B., Cook, P. A., Zheng, Y., Egan, A., Yushkevich, P. A., & Gee, J. C. (2010). N4ITK: improved N3 bias correction. IEEE transactions on medical imaging, 29(6), 1310–1320. 10.1109/TMI.2010.2046908

Yang, L., & Huang, M. (2024). Childhood maltreatment and mentalizing capacity: A meta-analysis. Child Abuse & Neglect, 149, 1–12. 10.1016/j.chiabu.2023.106623

Yeung, E. K. L., Apperly, I. A., & Devine, R. T. (2024). Measures of individual differences in adult theory of mind: A systematic review. Neuroscience and Biobehavioral Reviews, 157, 105481. 10.1016/j.neubiorev.2023.105481

Vai, B., Riberto, M., Ghiglino, D., Bollettini, I., Falini, A., Benedetti, F., & Poletti, S. (2018). Mild adverse childhood experiences increase neural efficacy during affective theory of mind. Stress (Amsterdam, Netherlands), 21(1), 84–89. 10.1080/10253890.2017.1398231

Vänskä, M., Punamäki, R. L., Tolvanen, A., Lindblom, J., Flykt, M., Unkila-Kallio, L., … & Tulppala, M. (2011). Maternal pre-and postnatal mental health trajectories and child mental health and development: Prospective study in a normative and formerly infertile sample. International Journal of Behavioral Development, 35(6), 517–531. 10.1177/0165025411417505

Wagner-Skacel, J., Riedl, D., Kampling, H., & Lampe, A. (2022). Mentalization and dissociation after adverse childhood experiences. Scientific Reports, 12(1), 6809. 10.1038/s41598-022-10787-8

Wang, S., Peterson, D. J., Gatenby, J. C., Li, W., Grabowski, T. J., & Madhyastha, T. M. (2017). Evaluation of Field Map and Nonlinear Registration Methods for Correction of Susceptibility Artifacts in Diffusion MRI. Frontiers in Neuroinformatics, 11, 17. 10.3389/fninf.2017.00017

Wikman, P., Moisala, M., Ylinen, A., Lindblom, J., Leikas, S., Salmela-Aro, K., Lonka, K., Güroğlu, B., & Alho, K. (2022). Brain Responses to Peer Feedback in Social Media Are Modulated by Valence in Late Adolescence. Frontiers in Behavioral Neuroscience, 16, 790478. 10.3389/fnbeh.2022.790478

Wimmer, H., & Perner, J. (1983). Beliefs about beliefs: representation and constraining function of wrong beliefs in young children’s understanding of deception. Cognition, 13(1), 103–128. 10.1016/0010-0277(83)90004-5

Winkler, A. M., Ridgway, G. R., Webster, M. A., Smith, S. M., & Nichols, T. E. (2014). Permutation inference for the general linear model. NeuroImage, 92(100), 381–397. 10.1016/j.neuroimage.2014.01.060

Woolrich, M. W., Ripley, B. D., Brady, M., & Smith, S. M. (2001). Temporal autocorrelation in univariate linear modeling of FMRI data. NeuroImage, 14(6), 1370–1386. 10.1006/nimg.2001.0931

Young, E. S., Frankenhuis, W. E., DelPriore, D. J., & Ellis, B. J. (2022). Hidden talents in context: Cognitive performance with abstract versus ecological stimuli among adversity-exposed youth. Child Development, 93(5), 1493–1510. 10.1111/cdev.13766

Zhang, Y., Brady, M., & Smith, S. (2001). Segmentation of brain MR images through a hidden Markov random field model and the expectation-maximization algorithm. IEEE transactions on medical imaging, 20(1), 45–57. 10.1109/42.906424

