## Supplementary Table S2 for "The Influence of Early Life Stress on the Development of Neural Representations during Mentalizing"

**Supplementary Table S2. Inter-subject representational similarity analysis across all HCP-MMP parcels: matching and Anna Karenina components of early life stress effects, under every correction applied.**

**Prospective ELS — Unadjusted (N = 89, 20,000 permutations, 360 parcels, r(|a − b|, mean) = .31)**

|  | | **Matching ρ(D, \|a − b\|)** | | | | | **Additive ρ(D, mean(a, b))** | | | | | **Joint model D ~ \|a − b\| + mean(a, b)** | | | |  |
| --- | --- | --- | --- | --- | --- | --- | --- | --- | --- | --- | --- | --- | --- | --- | --- | --- |
| **Hemi** | **Region** | **ρ** | **p** | **q** | **p FWE-P** | **p FWE-F** | **ρ** | **p** | **q** | **p FWE-P** | **p FWE-F** | **β \|a−b\|** | **p** | **β mean** | **p** | **Mechanism** |
| lh | V1 | .054 | .122 | .361 | .963 | >.999 | −.007 | .932 | .993 | >.999 | >.999 | .062 | .162 | −.026 | .749 | — |
| lh | MST | .004 | .437 | .560 | >.999 | >.999 | .031 | .417 | .993 | >.999 | >.999 | −.006 | .795 | .033 | .402 | — |
| lh | V6 | .020 | .272 | .441 | >.999 | >.999 | −.020 | .722 | .993 | >.999 | >.999 | .029 | .379 | −.029 | .614 | — |
| lh | V2 | .044 | .178 | .404 | .991 | >.999 | −.031 | .706 | .993 | >.999 | >.999 | .059 | .201 | −.050 | .558 | — |
| lh | V3 | .025 | .282 | .442 | >.999 | >.999 | −.027 | .743 | .993 | >.999 | >.999 | .037 | .404 | −.038 | .646 | — |
| lh | V4 | −.010 | .573 | .657 | >.999 | >.999 | −.055 | .499 | .993 | >.999 | >.999 | .009 | .853 | −.058 | .494 | — |
| lh | V8 | .015 | .345 | .488 | >.999 | >.999 | .000 | .999 | .999 | >.999 | >.999 | .017 | .666 | −.005 | .945 | — |
| lh | 4 | .043 | .181 | .404 | .992 | >.999 | −.020 | .801 | .993 | >.999 | >.999 | .055 | .229 | −.038 | .653 | — |
| lh | 3b | .033 | .209 | .418 | .999 | >.999 | .005 | .947 | .993 | >.999 | >.999 | .035 | .382 | −.006 | .927 | — |
| lh | FEF | .053 | .088 | .334 | .969 | >.999 | −.002 | .974 | .993 | >.999 | >.999 | .059 | .123 | −.021 | .761 | — |
| lh | PEF | −.025 | .766 | .811 | >.999 | >.999 | .011 | .845 | .993 | >.999 | >.999 | −.032 | .356 | .021 | .719 | — |
| lh | 55b | .036 | .205 | .418 | .998 | >.999 | −.029 | .709 | .993 | >.999 | >.999 | .050 | .255 | −.044 | .572 | — |
| lh | V3A | −.039 | .864 | .883 | >.999 | >.999 | −.048 | .424 | .993 | >.999 | >.999 | −.027 | .447 | −.039 | .523 | — |
| lh | RSC | −.027 | .786 | .830 | >.999 | >.999 | −.107 | **.049** | .993 | >.999 | >.999 | .007 | .822 | −.109 | .052 | — |
| lh | POS2 | .061 | .070 | .334 | .919 | >.999 | .051 | .476 | .993 | >.999 | >.999 | .050 | .211 | .035 | .636 | — |
| lh | V7 | .014 | .313 | .457 | >.999 | >.999 | .056 | .230 | .993 | >.999 | >.999 | −.004 | .884 | .058 | .234 | — |
| lh | IPS1 | −.013 | .615 | .694 | >.999 | >.999 | −.072 | .283 | .993 | >.999 | >.999 | .011 | .772 | −.076 | .271 | — |
| lh | FFC | .057 | .111 | .361 | .946 | >.999 | −.057 | .486 | .993 | >.999 | >.999 | .083 | .067 | −.083 | .323 | — |
| lh | V3B | .045 | .054 | .334 | .989 | >.999 | −.029 | .523 | .993 | >.999 | >.999 | .060 | **.032** | −.047 | .303 | Matching |
| lh | LO1 | −.001 | .510 | .610 | >.999 | >.999 | −.022 | .683 | .993 | >.999 | >.999 | .007 | .837 | −.024 | .662 | — |
| lh | LO2 | .042 | .130 | .365 | .993 | >.999 | .014 | .822 | .993 | >.999 | >.999 | .042 | .253 | .001 | .992 | — |
| lh | PIT | .047 | .114 | .361 | .986 | >.999 | .002 | .978 | .993 | >.999 | >.999 | .051 | .177 | −.014 | .836 | — |
| lh | MT | −.029 | .851 | .880 | >.999 | >.999 | −.035 | .443 | .993 | >.999 | >.999 | −.020 | .478 | −.028 | .543 | — |
| lh | A1 | .019 | .276 | .441 | >.999 | >.999 | −.058 | .273 | .993 | >.999 | >.999 | .041 | .195 | −.071 | .196 | — |
| lh | PSL | −.003 | .515 | .612 | >.999 | >.999 | −.033 | .691 | .993 | >.999 | >.999 | .008 | .858 | −.036 | .679 | — |
| lh | SFL | .075 | .058 | .334 | .770 | >.999 | .088 | .283 | .993 | >.999 | >.999 | .053 | .246 | .071 | .400 | — |
| lh | PCV | .020 | .284 | .442 | >.999 | >.999 | −.017 | .776 | .993 | >.999 | >.999 | .029 | .427 | −.026 | .681 | — |
| lh | STV | .052 | .138 | .384 | .970 | >.999 | .045 | .590 | .993 | >.999 | >.999 | .042 | .369 | .032 | .712 | — |
| lh | 7Pm | .026 | .229 | .427 | >.999 | >.999 | −.033 | .584 | .993 | >.999 | >.999 | .040 | .251 | −.045 | .464 | — |
| lh | 7m | .046 | .146 | .394 | .987 | >.999 | .054 | .478 | .993 | >.999 | >.999 | .033 | .448 | .044 | .574 | — |
| lh | POS1 | −.002 | .513 | .612 | >.999 | >.999 | −.056 | .412 | .993 | >.999 | >.999 | .017 | .662 | −.061 | .380 | — |
| lh | 23d | .029 | .211 | .419 | >.999 | >.999 | .078 | .196 | .993 | >.999 | >.999 | .005 | .890 | .076 | .220 | — |
| lh | v23ab | .044 | .080 | .334 | .989 | >.999 | .074 | .146 | .993 | >.999 | >.999 | .023 | .453 | .067 | .207 | — |
| lh | d23ab | .013 | .364 | .498 | >.999 | >.999 | .005 | .940 | .993 | >.999 | >.999 | .013 | .741 | .001 | .989 | — |
| lh | 31pv | .088 | **.005** | .279 | .571 | >.999 | .129 | **.018** | .993 | .996 | .996 | .053 | .104 | .112 | **.046** | AK+ |
| lh | 5m | .013 | .336 | .483 | >.999 | >.999 | .081 | .097 | .993 | >.999 | >.999 | −.014 | .635 | .086 | .089 | — |
| lh | 5mv | .053 | **.046** | .334 | .969 | >.999 | .017 | .733 | .993 | >.999 | >.999 | .052 | .088 | .001 | .982 | — |
| lh | 23c | .046 | .084 | .334 | .988 | >.999 | −.031 | .564 | .993 | >.999 | >.999 | .061 | .062 | −.050 | .369 | — |
| lh | 5L | .025 | .224 | .425 | >.999 | >.999 | −.001 | .978 | .993 | >.999 | >.999 | .029 | .394 | −.010 | .855 | — |
| lh | 24dd | .035 | .179 | .404 | .999 | >.999 | .016 | .802 | .993 | >.999 | >.999 | .034 | .373 | .006 | .934 | — |
| lh | 24dv | .058 | **.036** | .334 | .945 | >.999 | .005 | .926 | .993 | >.999 | >.999 | .062 | **.046** | −.015 | .785 | Matching |
| lh | 7AL | −.015 | .653 | .722 | >.999 | >.999 | .001 | .983 | .993 | >.999 | >.999 | −.017 | .637 | .006 | .918 | — |
| lh | SCEF | .027 | .271 | .441 | >.999 | >.999 | .001 | .987 | .993 | >.999 | >.999 | .030 | .517 | −.008 | .923 | — |
| lh | 6ma | .026 | .270 | .441 | >.999 | >.999 | −.021 | .780 | .993 | >.999 | >.999 | .036 | .390 | −.033 | .678 | — |
| lh | 7Am | .012 | .374 | .506 | >.999 | >.999 | .035 | .599 | .993 | >.999 | >.999 | .001 | .987 | .035 | .608 | — |
| lh | 7Pl | .022 | .260 | .441 | >.999 | >.999 | .043 | .465 | .993 | >.999 | >.999 | .010 | .777 | .040 | .511 | — |
| lh | 7PC | .031 | .222 | .425 | >.999 | >.999 | .063 | .349 | .993 | >.999 | >.999 | .012 | .763 | .060 | .391 | — |
| lh | LIPv | −.006 | .547 | .635 | >.999 | >.999 | .040 | .532 | .993 | >.999 | >.999 | −.020 | .594 | .046 | .483 | — |
| lh | VIP | .026 | .257 | .441 | >.999 | >.999 | .075 | .278 | .993 | >.999 | >.999 | .003 | .947 | .074 | .298 | — |
| lh | MIP | .010 | .387 | .518 | >.999 | >.999 | −.028 | .664 | .993 | >.999 | >.999 | .021 | .574 | −.034 | .602 | — |
| lh | 1 | .044 | .160 | .396 | .991 | >.999 | −.006 | .939 | .993 | >.999 | >.999 | .050 | .244 | −.021 | .788 | — |
| lh | 2 | .069 | **.044** | .334 | .849 | >.999 | .059 | .383 | .993 | >.999 | >.999 | .055 | .152 | .042 | .548 | — |
| lh | 3a | −.013 | .647 | .718 | >.999 | >.999 | −.083 | .156 | .993 | >.999 | >.999 | .014 | .681 | −.088 | .147 | — |
| lh | 6d | .005 | .446 | .564 | >.999 | >.999 | .001 | .987 | .993 | >.999 | >.999 | .005 | .896 | −.000 | .995 | — |
| lh | 6mp | .055 | .063 | .334 | .960 | >.999 | .036 | .544 | .993 | >.999 | >.999 | .048 | .169 | .021 | .732 | — |
| lh | 6v | .003 | .464 | .573 | >.999 | >.999 | .025 | .709 | .993 | >.999 | >.999 | −.006 | .888 | .027 | .698 | — |
| lh | p24pr | .045 | .101 | .350 | .989 | >.999 | −.008 | .902 | .993 | >.999 | >.999 | .052 | .132 | −.024 | .698 | — |
| lh | 33pr | .007 | .402 | .528 | >.999 | >.999 | .038 | .370 | .993 | >.999 | >.999 | −.006 | .829 | .040 | .361 | — |
| lh | a24pr | .028 | .192 | .411 | >.999 | >.999 | .032 | .551 | .993 | >.999 | >.999 | .020 | .536 | .026 | .638 | — |
| lh | p32pr | .036 | .129 | .365 | .998 | >.999 | .024 | .660 | .993 | >.999 | >.999 | .032 | .317 | .013 | .805 | — |
| lh | a24 | .056 | .055 | .334 | .953 | >.999 | −.006 | .919 | .993 | >.999 | >.999 | .064 | .059 | −.026 | .661 | — |
| lh | d32 | .026 | .245 | .441 | >.999 | >.999 | .041 | .534 | .993 | >.999 | >.999 | .015 | .699 | .036 | .592 | — |
| lh | 8BM | .075 | .058 | .334 | .776 | >.999 | −.016 | .838 | .993 | >.999 | >.999 | .088 | **.048** | −.044 | .595 | Matching |
| lh | p32 | −.003 | .544 | .635 | >.999 | >.999 | .036 | .407 | .993 | >.999 | >.999 | −.016 | .561 | .041 | .361 | — |
| lh | 10r | .044 | .084 | .334 | .990 | >.999 | .033 | .532 | .993 | >.999 | >.999 | .038 | .230 | .021 | .691 | — |
| lh | 47m | .019 | .266 | .441 | >.999 | >.999 | .044 | .386 | .993 | >.999 | >.999 | .006 | .853 | .042 | .421 | — |
| lh | 8Av | .088 | **.039** | .334 | .567 | >.999 | .072 | .394 | .993 | >.999 | >.999 | .073 | .127 | .050 | .571 | — |
| lh | 8Ad | .004 | .449 | .564 | >.999 | >.999 | .044 | .520 | .993 | >.999 | >.999 | −.011 | .786 | .047 | .500 | — |
| lh | 9m | .046 | .166 | .398 | .988 | >.999 | .016 | .844 | .993 | >.999 | >.999 | .045 | .330 | .002 | .979 | — |
| lh | 8BL | .096 | **.014** | .334 | .460 | >.999 | .035 | .628 | .993 | >.999 | >.999 | .094 | **.024** | .006 | .937 | Matching |
| lh | 9p | .029 | .233 | .431 | >.999 | >.999 | .061 | .369 | .993 | >.999 | >.999 | .010 | .787 | .058 | .409 | — |
| lh | 10d | .050 | .080 | .334 | .978 | >.999 | .024 | .672 | .993 | >.999 | >.999 | .047 | .173 | .010 | .869 | — |
| lh | 8C | .135 | **.003** | .279 | .091 | .988 | .144 | .088 | .993 | .955 | .957 | .100 | **.034** | .112 | .195 | Matching |
| lh | 44 | .089 | **.038** | .334 | .562 | >.999 | .119 | .164 | .993 | >.999 | >.999 | .057 | .235 | .101 | .251 | — |
| lh | 45 | .074 | .079 | .334 | .783 | >.999 | −.023 | .796 | .993 | >.999 | >.999 | .090 | .070 | −.052 | .576 | — |
| lh | 47l | .070 | .060 | .334 | .836 | >.999 | −.025 | .733 | .993 | >.999 | >.999 | .086 | **.042** | −.052 | .496 | Matching |
| lh | a47r | .073 | **.036** | .334 | .795 | >.999 | .045 | .512 | .993 | >.999 | >.999 | .065 | .094 | .025 | .726 | — |
| lh | 6r | .061 | .091 | .339 | .924 | >.999 | .120 | .127 | .993 | >.999 | >.999 | .026 | .557 | .112 | .164 | — |
| lh | IFJa | −.024 | .731 | .783 | >.999 | >.999 | .059 | .373 | .993 | >.999 | >.999 | −.048 | .211 | .074 | .280 | — |
| lh | IFJp | .021 | .263 | .441 | >.999 | >.999 | .059 | .285 | .993 | >.999 | >.999 | .003 | .930 | .058 | .309 | — |
| lh | IFSp | −.017 | .637 | .712 | >.999 | >.999 | .023 | .767 | .993 | >.999 | >.999 | −.027 | .539 | .031 | .696 | — |
| lh | IFSaᵃ | .178 | **<.001** | .072 | **.006** | .601 | .295 | **<.001** | .198 | **.009** | **.009** | .095 | .052 | .265 | **.003** | AK+ |
| lh | p9-46v | .057 | .124 | .364 | .948 | >.999 | .028 | .745 | .993 | >.999 | >.999 | .054 | .267 | .012 | .897 | — |
| lh | 46 | .033 | .220 | .425 | .999 | >.999 | −.003 | .972 | .993 | >.999 | >.999 | .038 | .380 | −.014 | .856 | — |
| lh | a9-46v | .054 | .103 | .351 | .963 | >.999 | .057 | .447 | .993 | >.999 | >.999 | .040 | .339 | .044 | .566 | — |
| lh | 9-46d | .023 | .300 | .446 | >.999 | >.999 | −.008 | .917 | .993 | >.999 | >.999 | .028 | .526 | −.017 | .836 | — |
| lh | 9a | .066 | .085 | .334 | .876 | >.999 | −.025 | .761 | .993 | >.999 | >.999 | .082 | .073 | −.051 | .544 | — |
| lh | 10v | .054 | .060 | .334 | .963 | >.999 | .011 | .853 | .993 | >.999 | >.999 | .056 | .098 | −.007 | .909 | — |
| lh | a10p | .005 | .436 | .560 | >.999 | >.999 | −.023 | .667 | .993 | >.999 | >.999 | .013 | .675 | −.027 | .618 | — |
| lh | 10pp | .047 | .061 | .334 | .985 | >.999 | .023 | .653 | .993 | >.999 | >.999 | .044 | .136 | .009 | .866 | — |
| lh | 11l | .040 | .145 | .392 | .996 | >.999 | −.020 | .757 | .993 | >.999 | >.999 | .051 | .162 | −.036 | .586 | — |
| lh | 13l | .024 | .259 | .441 | >.999 | >.999 | .036 | .560 | .993 | >.999 | >.999 | .014 | .704 | .032 | .623 | — |
| lh | OFC | .013 | .353 | .488 | >.999 | >.999 | −.089 | .125 | .993 | >.999 | >.999 | .046 | .182 | −.103 | .085 | — |
| lh | 47s | .013 | .350 | .488 | >.999 | >.999 | −.050 | .393 | .993 | >.999 | >.999 | .032 | .352 | −.060 | .318 | — |
| lh | LIPd | .047 | .060 | .334 | .984 | >.999 | .004 | .934 | .993 | >.999 | >.999 | .051 | .091 | −.012 | .821 | — |
| lh | 6a | .036 | .168 | .400 | .998 | >.999 | .074 | .240 | .993 | >.999 | >.999 | .014 | .700 | .070 | .284 | — |
| lh | i6-8 | .043 | .118 | .361 | .992 | >.999 | .100 | .098 | .993 | >.999 | >.999 | .013 | .723 | .096 | .121 | — |
| lh | s6-8 | .043 | .099 | .348 | .991 | >.999 | .081 | .141 | .993 | >.999 | >.999 | .020 | .545 | .075 | .187 | — |
| lh | 43 | .014 | .337 | .483 | >.999 | >.999 | .062 | .246 | .993 | >.999 | >.999 | −.006 | .843 | .064 | .243 | — |
| lh | OP4 | .061 | .062 | .334 | .920 | >.999 | .047 | .477 | .993 | >.999 | >.999 | .051 | .177 | .031 | .650 | — |
| lh | OP1 | .013 | .351 | .488 | >.999 | >.999 | .012 | .845 | .993 | >.999 | >.999 | .011 | .756 | .008 | .891 | — |
| lh | OP2-3 | −.010 | .641 | .715 | >.999 | >.999 | −.020 | .651 | .993 | >.999 | >.999 | −.004 | .873 | −.018 | .687 | — |
| lh | 52 | −.055 | .973 | .984 | >.999 | >.999 | −.084 | .064 | .993 | >.999 | >.999 | −.032 | .272 | −.074 | .115 | — |
| lh | RI | .015 | .314 | .457 | >.999 | >.999 | .009 | .861 | .993 | >.999 | >.999 | .013 | .653 | .004 | .932 | — |
| lh | PFcm | .071 | **.034** | .334 | .825 | >.999 | .053 | .396 | .993 | >.999 | >.999 | .060 | .100 | .034 | .596 | — |
| lh | PoI2 | .024 | .254 | .441 | >.999 | >.999 | −.046 | .447 | .993 | >.999 | >.999 | .043 | .234 | −.060 | .343 | — |
| lh | TA2 | .033 | .160 | .396 | .999 | >.999 | −.008 | .879 | .993 | >.999 | >.999 | .039 | .223 | −.020 | .709 | — |
| lh | FOP4 | .054 | .076 | .334 | .961 | >.999 | .131 | **.038** | .993 | .994 | .994 | .015 | .690 | .126 | .052 | — |
| lh | MI | −.023 | .722 | .779 | >.999 | >.999 | −.076 | .239 | .993 | >.999 | >.999 | .001 | .976 | −.077 | .251 | — |
| lh | Pir | −.015 | .680 | .739 | >.999 | >.999 | −.027 | .603 | .993 | >.999 | >.999 | −.007 | .826 | −.025 | .642 | — |
| lh | AVI | .052 | .082 | .334 | .971 | >.999 | .059 | .350 | .993 | >.999 | >.999 | .037 | .307 | .047 | .465 | — |
| lh | AAIC | −.021 | .730 | .783 | >.999 | >.999 | −.079 | .155 | .993 | >.999 | >.999 | .004 | .900 | −.081 | .161 | — |
| lh | FOP1 | .003 | .466 | .575 | >.999 | >.999 | .003 | .958 | .993 | >.999 | >.999 | .002 | .946 | .002 | .968 | — |
| lh | FOP3 | −.063 | .984 | .992 | >.999 | >.999 | −.018 | .688 | .993 | >.999 | >.999 | −.063 | **.027** | .001 | .976 | Matching |
| lh | FOP2 | .042 | .060 | .334 | .993 | >.999 | −.042 | .308 | .993 | >.999 | >.999 | .061 | **.021** | −.061 | .150 | Matching |
| lh | PFt | .019 | .272 | .441 | >.999 | >.999 | −.067 | .194 | .993 | >.999 | >.999 | .044 | .157 | −.081 | .127 | — |
| lh | AIP | .006 | .431 | .556 | >.999 | >.999 | −.028 | .640 | .993 | >.999 | >.999 | .016 | .656 | −.033 | .597 | — |
| lh | EC | .008 | .407 | .530 | >.999 | >.999 | −.040 | .524 | .993 | >.999 | >.999 | .023 | .530 | −.047 | .464 | — |
| lh | PreS | .020 | .295 | .442 | >.999 | >.999 | −.064 | .294 | .993 | >.999 | >.999 | .044 | .221 | −.078 | .214 | — |
| lh | H | .064 | **.034** | .334 | .901 | >.999 | .021 | .708 | .993 | >.999 | >.999 | .063 | .057 | .001 | .978 | — |
| lh | ProS | .024 | .187 | .404 | >.999 | >.999 | .053 | .211 | .993 | >.999 | >.999 | .008 | .760 | .051 | .248 | — |
| lh | PeEc | .039 | .187 | .404 | .996 | >.999 | −.018 | .817 | .993 | >.999 | >.999 | .049 | .250 | −.033 | .674 | — |
| lh | STGa | .019 | .312 | .457 | >.999 | >.999 | .037 | .580 | .993 | >.999 | >.999 | .008 | .830 | .035 | .617 | — |
| lh | PBelt | −.014 | .663 | .728 | >.999 | >.999 | −.042 | .453 | .993 | >.999 | >.999 | −.001 | .977 | −.042 | .469 | — |
| lh | A5 | .098 | **.031** | .334 | .426 | >.999 | .034 | .696 | .993 | >.999 | >.999 | .097 | **.044** | .003 | .970 | Matching |
| lh | PHA1 | .022 | .257 | .441 | >.999 | >.999 | −.140 | **.012** | .993 | .970 | .972 | .073 | **.031** | −.163 | **.005** | Match + AK− |
| lh | PHA3 | .011 | .377 | .507 | >.999 | >.999 | −.017 | .774 | .993 | >.999 | >.999 | .018 | .619 | −.023 | .718 | — |
| lh | STSda | .055 | .086 | .334 | .956 | >.999 | .045 | .509 | .993 | >.999 | >.999 | .046 | .245 | .031 | .658 | — |
| lh | STSdp | .050 | .114 | .361 | .978 | >.999 | .042 | .556 | .993 | >.999 | >.999 | .041 | .308 | .029 | .691 | — |
| lh | STSvp | .049 | .149 | .395 | .981 | >.999 | .041 | .621 | .993 | >.999 | >.999 | .040 | .395 | .029 | .735 | — |
| lh | TGd | .031 | .258 | .441 | >.999 | >.999 | .014 | .868 | .993 | >.999 | >.999 | .030 | .530 | .005 | .953 | — |
| lh | TE1a | .063 | .093 | .341 | .908 | >.999 | .024 | .773 | .993 | >.999 | >.999 | .061 | .179 | .005 | .955 | — |
| lh | TE1p | .026 | .292 | .442 | >.999 | >.999 | −.053 | .540 | .993 | >.999 | >.999 | .047 | .324 | −.068 | .447 | — |
| lh | TE2a | .014 | .367 | .499 | >.999 | >.999 | −.037 | .640 | .993 | >.999 | >.999 | .029 | .520 | −.046 | .567 | — |
| lh | TF | .024 | .285 | .442 | >.999 | >.999 | .002 | .983 | .993 | >.999 | >.999 | .027 | .532 | −.007 | .931 | — |
| lh | TE2p | .040 | .181 | .404 | .995 | >.999 | −.012 | .881 | .993 | >.999 | >.999 | .049 | .264 | −.027 | .735 | — |
| lh | PHT | .031 | .271 | .441 | >.999 | >.999 | −.059 | .519 | .993 | >.999 | >.999 | .054 | .279 | −.076 | .416 | — |
| lh | PH | .023 | .295 | .442 | >.999 | >.999 | −.027 | .716 | .993 | >.999 | >.999 | .035 | .410 | −.038 | .621 | — |
| lh | TPOJ1 | −.014 | .610 | .691 | >.999 | >.999 | −.035 | .655 | .993 | >.999 | >.999 | −.003 | .940 | −.034 | .673 | — |
| lh | TPOJ2 | .054 | .122 | .361 | .965 | >.999 | −.007 | .935 | .993 | >.999 | >.999 | .062 | .169 | −.026 | .750 | — |
| lh | TPOJ3 | .043 | .068 | .334 | .991 | >.999 | −.032 | .486 | .993 | >.999 | >.999 | .059 | **.040** | −.051 | .288 | Matching |
| lh | DVT | .056 | .072 | .334 | .955 | >.999 | .077 | .232 | .993 | >.999 | >.999 | .035 | .349 | .066 | .317 | — |
| lh | PGp | .025 | .260 | .441 | >.999 | >.999 | .042 | .540 | .993 | >.999 | >.999 | .013 | .742 | .038 | .589 | — |
| lh | IP2 | −.023 | .743 | .791 | >.999 | >.999 | −.036 | .526 | .993 | >.999 | >.999 | −.013 | .702 | −.032 | .583 | — |
| lh | IP1 | −.010 | .602 | .686 | >.999 | >.999 | −.010 | .870 | .993 | >.999 | >.999 | −.007 | .838 | −.008 | .903 | — |
| lh | IP0 | .045 | .097 | .348 | .989 | >.999 | −.073 | .211 | .993 | >.999 | >.999 | .076 | **.024** | −.097 | .106 | Matching |
| lh | PFop | −.026 | .766 | .811 | >.999 | >.999 | .002 | .980 | .993 | >.999 | >.999 | −.030 | .398 | .011 | .861 | — |
| lh | PF | .023 | .313 | .457 | >.999 | >.999 | .003 | .970 | .993 | >.999 | >.999 | .024 | .608 | −.004 | .961 | — |
| lh | PFm | .093 | **.024** | .334 | .501 | >.999 | .105 | .195 | .993 | >.999 | >.999 | .067 | .140 | .084 | .312 | — |
| lh | PGi | .064 | .091 | .339 | .895 | >.999 | .085 | .315 | .993 | >.999 | >.999 | .042 | .385 | .072 | .410 | — |
| lh | PGsᵃ | −.001 | .499 | .603 | >.999 | >.999 | .080 | .241 | .993 | >.999 | >.999 | −.029 | .459 | .089 | .207 | — |
| lh | V6A | −.033 | .874 | .892 | >.999 | >.999 | −.074 | .097 | .993 | >.999 | >.999 | −.010 | .717 | −.071 | .122 | — |
| lh | VMV1 | −.033 | .863 | .883 | >.999 | >.999 | .025 | .620 | .993 | >.999 | >.999 | −.045 | .137 | .039 | .447 | — |
| lh | VMV3 | .011 | .354 | .488 | >.999 | >.999 | .062 | .220 | .993 | >.999 | >.999 | −.009 | .774 | .064 | .214 | — |
| lh | PHA2 | .003 | .459 | .570 | >.999 | >.999 | −.024 | .520 | .993 | >.999 | >.999 | .011 | .648 | −.028 | .472 | — |
| lh | V4t | −.010 | .630 | .706 | >.999 | >.999 | −.095 | .057 | .993 | >.999 | >.999 | .022 | .480 | −.102 | **.047** | AK− |
| lh | FST | .016 | .344 | .488 | >.999 | >.999 | −.053 | .426 | .993 | >.999 | >.999 | .036 | .350 | −.065 | .350 | — |
| lh | V3CD | .035 | .160 | .396 | .999 | >.999 | .036 | .539 | .993 | >.999 | >.999 | .026 | .450 | .027 | .644 | — |
| lh | LO3 | −.073 | .992 | .997 | >.999 | >.999 | −.034 | .500 | .993 | >.999 | >.999 | −.069 | **.022** | −.013 | .810 | Matching |
| lh | VMV2 | −.033 | .898 | .913 | >.999 | >.999 | −.082 | **.044** | .993 | >.999 | >.999 | −.009 | .744 | −.079 | .060 | — |
| lh | 31pd | .018 | .272 | .441 | >.999 | >.999 | .006 | .904 | .993 | >.999 | >.999 | .018 | .547 | .000 | .995 | — |
| lh | 31a | .020 | .294 | .442 | >.999 | >.999 | −.034 | .595 | .993 | >.999 | >.999 | .034 | .359 | −.044 | .498 | — |
| lh | VVC | .051 | .115 | .361 | .975 | >.999 | .028 | .689 | .993 | >.999 | >.999 | .046 | .253 | .014 | .848 | — |
| lh | 25 | .030 | .160 | .396 | >.999 | >.999 | −.040 | .411 | .993 | >.999 | >.999 | .048 | .111 | −.055 | .272 | — |
| lh | s32 | .002 | .451 | .564 | >.999 | >.999 | −.000 | .988 | .993 | >.999 | >.999 | .002 | .917 | −.001 | .968 | — |
| lh | pOFC | .067 | **.041** | .334 | .867 | >.999 | −.008 | .900 | .993 | >.999 | >.999 | .077 | **.034** | −.032 | .623 | Matching |
| lh | PoI1 | −.005 | .554 | .641 | >.999 | >.999 | −.058 | .313 | .993 | >.999 | >.999 | .015 | .674 | −.062 | .292 | — |
| lh | Ig | .018 | .267 | .441 | >.999 | >.999 | .075 | .119 | .993 | >.999 | >.999 | −.006 | .851 | .077 | .118 | — |
| lh | FOP5 | −.035 | .811 | .846 | >.999 | >.999 | .056 | .400 | .993 | >.999 | >.999 | −.058 | .130 | .074 | .277 | — |
| lh | p10p | .034 | .158 | .396 | .999 | >.999 | −.008 | .888 | .993 | >.999 | >.999 | .041 | .220 | −.021 | .721 | — |
| lh | p47r | .106 | **.009** | .279 | .327 | >.999 | .112 | .133 | .993 | >.999 | >.999 | .079 | .066 | .088 | .259 | — |
| lh | TGv | −.018 | .663 | .728 | >.999 | >.999 | .020 | .782 | .993 | >.999 | >.999 | −.027 | .506 | .029 | .697 | — |
| lh | MBelt | .053 | .050 | .334 | .967 | >.999 | −.051 | .347 | .993 | >.999 | >.999 | .077 | **.015** | −.075 | .176 | Matching |
| lh | LBelt | .023 | .208 | .418 | >.999 | >.999 | .007 | .876 | .993 | >.999 | >.999 | .023 | .421 | −.000 | .996 | — |
| lh | A4 | .070 | .081 | .334 | .834 | >.999 | .035 | .684 | .993 | >.999 | >.999 | .066 | .172 | .014 | .873 | — |
| lh | STSva | .004 | .454 | .566 | >.999 | >.999 | −.008 | .895 | .993 | >.999 | >.999 | .007 | .842 | −.010 | .870 | — |
| lh | TE1m | .061 | .076 | .334 | .921 | >.999 | .033 | .644 | .993 | >.999 | >.999 | .056 | .167 | .015 | .837 | — |
| lh | PI | −.002 | .530 | .626 | >.999 | >.999 | −.069 | .233 | .993 | >.999 | >.999 | .022 | .533 | −.076 | .204 | — |
| lh | a32pr | .039 | .159 | .396 | .996 | >.999 | .009 | .894 | .993 | >.999 | >.999 | .040 | .288 | −.004 | .959 | — |
| lh | p24 | .005 | .449 | .564 | >.999 | >.999 | −.028 | .667 | .993 | >.999 | >.999 | .015 | .692 | −.032 | .623 | — |
| rh | V1 | .024 | .294 | .442 | >.999 | >.999 | −.025 | .754 | .993 | >.999 | >.999 | .035 | .423 | −.036 | .653 | — |
| rh | MST | .019 | .287 | .442 | >.999 | >.999 | −.031 | .562 | .993 | >.999 | >.999 | .032 | .330 | −.041 | .456 | — |
| rh | V6 | .013 | .341 | .488 | >.999 | >.999 | .054 | .309 | .993 | >.999 | >.999 | −.004 | .896 | .055 | .311 | — |
| rh | V2 | .040 | .193 | .411 | .995 | >.999 | .026 | .751 | .993 | >.999 | >.999 | .035 | .436 | .015 | .860 | — |
| rh | V3 | .036 | .206 | .418 | .998 | >.999 | .037 | .639 | .993 | >.999 | >.999 | .027 | .529 | .028 | .725 | — |
| rh | V4 | .026 | .273 | .441 | >.999 | >.999 | −.037 | .620 | .993 | >.999 | >.999 | .042 | .326 | −.050 | .516 | — |
| rh | V8 | .051 | .075 | .334 | .975 | >.999 | .020 | .734 | .993 | >.999 | >.999 | .050 | .146 | .004 | .942 | — |
| rh | 4 | .060 | .095 | .346 | .926 | >.999 | .025 | .751 | .993 | >.999 | >.999 | .058 | .191 | .007 | .932 | — |
| rh | 3b | .072 | **.047** | .334 | .812 | >.999 | .031 | .671 | .993 | >.999 | >.999 | .069 | .093 | .009 | .900 | — |
| rh | FEF | .054 | .081 | .334 | .961 | >.999 | .016 | .811 | .993 | >.999 | >.999 | .055 | .144 | −.001 | .983 | — |
| rh | PEF | .008 | .404 | .529 | >.999 | >.999 | .039 | .491 | .993 | >.999 | >.999 | −.005 | .891 | .040 | .489 | — |
| rh | 55b | .054 | .098 | .348 | .963 | >.999 | .016 | .814 | .993 | >.999 | >.999 | .054 | .180 | −.000 | .994 | — |
| rh | V3A | −.031 | .804 | .841 | >.999 | >.999 | −.022 | .718 | .993 | >.999 | >.999 | −.027 | .453 | −.014 | .825 | — |
| rh | RSC | .047 | .063 | .334 | .985 | >.999 | .032 | .527 | .993 | >.999 | >.999 | .041 | .171 | .019 | .717 | — |
| rh | POS2 | .015 | .356 | .489 | >.999 | >.999 | −.010 | .897 | .993 | >.999 | >.999 | .021 | .632 | −.017 | .832 | — |
| rh | V7 | −.005 | .545 | .635 | >.999 | >.999 | .053 | .334 | .993 | >.999 | >.999 | −.023 | .479 | .060 | .283 | — |
| rh | IPS1 | .003 | .471 | .577 | >.999 | >.999 | .013 | .823 | .993 | >.999 | >.999 | −.002 | .961 | .014 | .821 | — |
| rh | FFC | .071 | .058 | .334 | .819 | >.999 | .124 | .099 | .993 | .999 | .999 | .036 | .397 | .112 | .144 | — |
| rh | V3B | .003 | .452 | .564 | >.999 | >.999 | .078 | .066 | .993 | >.999 | >.999 | −.024 | .387 | .085 | .052 | — |
| rh | LO1 | −.031 | .842 | .874 | >.999 | >.999 | −.020 | .697 | .993 | >.999 | >.999 | −.027 | .372 | −.011 | .833 | — |
| rh | LO2 | .022 | .250 | .441 | >.999 | >.999 | .009 | .865 | .993 | >.999 | >.999 | .021 | .504 | .002 | .964 | — |
| rh | PIT | .024 | .267 | .441 | >.999 | >.999 | .094 | .154 | .993 | >.999 | >.999 | −.006 | .879 | .096 | .159 | — |
| rh | MT | −.001 | .503 | .606 | >.999 | >.999 | −.020 | .695 | .993 | >.999 | >.999 | .006 | .841 | −.022 | .675 | — |
| rh | A1 | −.007 | .576 | .658 | >.999 | >.999 | −.031 | .543 | .993 | >.999 | >.999 | .004 | .913 | −.032 | .541 | — |
| rh | PSL | .056 | .099 | .348 | .952 | >.999 | .023 | .761 | .993 | >.999 | >.999 | .054 | .203 | .006 | .938 | — |
| rh | SFL | .057 | .087 | .334 | .950 | >.999 | .094 | .192 | .993 | >.999 | >.999 | .030 | .462 | .084 | .256 | — |
| rh | PCV | .055 | .074 | .334 | .957 | >.999 | .018 | .785 | .993 | >.999 | >.999 | .055 | .139 | .001 | .992 | — |
| rh | STV | .092 | **.028** | .334 | .519 | >.999 | .044 | .590 | .993 | >.999 | >.999 | .086 | .063 | .017 | .837 | — |
| rh | 7Pm | −.009 | .604 | .686 | >.999 | >.999 | −.024 | .673 | .993 | >.999 | >.999 | −.002 | .947 | −.023 | .690 | — |
| rh | 7m | .055 | .071 | .334 | .960 | >.999 | .057 | .362 | .993 | >.999 | >.999 | .041 | .259 | .044 | .494 | — |
| rh | POS1 | .067 | **.044** | .334 | .871 | >.999 | .079 | .221 | .993 | >.999 | >.999 | .046 | .210 | .064 | .330 | — |
| rh | 23d | .017 | .278 | .441 | >.999 | >.999 | −.022 | .654 | .993 | >.999 | >.999 | .026 | .374 | −.030 | .549 | — |
| rh | v23ab | .007 | .395 | .521 | >.999 | >.999 | .013 | .764 | .993 | >.999 | >.999 | .003 | .913 | .012 | .787 | — |
| rh | d23ab | .027 | .214 | .420 | >.999 | >.999 | .043 | .458 | .993 | >.999 | >.999 | .015 | .651 | .038 | .523 | — |
| rh | 31pv | .027 | .208 | .418 | >.999 | >.999 | .036 | .513 | .993 | >.999 | >.999 | .017 | .607 | .031 | .587 | — |
| rh | 5m | .040 | .117 | .361 | .995 | >.999 | .060 | .288 | .993 | >.999 | >.999 | .024 | .475 | .052 | .367 | — |
| rh | 5mv | .005 | .441 | .563 | >.999 | >.999 | −.054 | .386 | .993 | >.999 | >.999 | .024 | .509 | −.062 | .338 | — |
| rh | 23c | −.005 | .544 | .635 | >.999 | >.999 | −.034 | .568 | .993 | >.999 | >.999 | .006 | .863 | −.035 | .556 | — |
| rh | 5L | .004 | .443 | .564 | >.999 | >.999 | −.002 | .967 | .993 | >.999 | >.999 | .006 | .863 | −.004 | .944 | — |
| rh | 24dd | .021 | .278 | .441 | >.999 | >.999 | −.006 | .917 | .993 | >.999 | >.999 | .025 | .480 | −.014 | .821 | — |
| rh | 24dv | .038 | .128 | .365 | .997 | >.999 | .105 | .059 | .993 | >.999 | >.999 | .006 | .859 | .103 | .073 | — |
| rh | 7AL | .024 | .238 | .435 | >.999 | >.999 | −.003 | .953 | .993 | >.999 | >.999 | .028 | .409 | −.012 | .831 | — |
| rh | SCEF | .023 | .272 | .441 | >.999 | >.999 | −.045 | .488 | .993 | >.999 | >.999 | .041 | .271 | −.058 | .383 | — |
| rh | 6ma | .094 | **.016** | .334 | .484 | >.999 | −.048 | .526 | .993 | >.999 | >.999 | .121 | **.004** | −.085 | .268 | Matching |
| rh | 7Am | .009 | .390 | .520 | >.999 | >.999 | .012 | .854 | .993 | >.999 | >.999 | .006 | .864 | .010 | .879 | — |
| rh | 7Pl | .015 | .307 | .454 | >.999 | >.999 | .050 | .314 | .993 | >.999 | >.999 | −.001 | .980 | .051 | .327 | — |
| rh | 7PC | .044 | .120 | .361 | .991 | >.999 | .006 | .924 | .993 | >.999 | >.999 | .047 | .203 | −.009 | .893 | — |
| rh | LIPv | .039 | .116 | .361 | .996 | >.999 | .018 | .741 | .993 | >.999 | >.999 | .037 | .253 | .007 | .910 | — |
| rh | VIP | .012 | .351 | .488 | >.999 | >.999 | .018 | .739 | .993 | >.999 | >.999 | .007 | .822 | .015 | .778 | — |
| rh | MIP | .058 | .081 | .334 | .944 | >.999 | .023 | .745 | .993 | >.999 | >.999 | .056 | .162 | .006 | .934 | — |
| rh | 1 | .040 | .176 | .404 | .995 | >.999 | .035 | .643 | .993 | >.999 | >.999 | .032 | .447 | .025 | .752 | — |
| rh | 2 | .066 | **.049** | .334 | .874 | >.999 | −.022 | .750 | .993 | >.999 | >.999 | .081 | **.036** | −.047 | .503 | Matching |
| rh | 3a | .055 | **.046** | .334 | .956 | >.999 | .024 | .659 | .993 | >.999 | >.999 | .053 | .103 | .008 | .893 | — |
| rh | 6d | .051 | .088 | .334 | .974 | >.999 | −.009 | .894 | .993 | >.999 | >.999 | .060 | .105 | −.027 | .680 | — |
| rh | 6mp | .064 | **.049** | .334 | .896 | >.999 | −.020 | .757 | .993 | >.999 | >.999 | .078 | **.035** | −.045 | .503 | Matching |
| rh | 6v | .045 | .145 | .392 | .989 | >.999 | .007 | .931 | .993 | >.999 | >.999 | .048 | .256 | −.008 | .909 | — |
| rh | p24pr | .016 | .293 | .442 | >.999 | >.999 | .030 | .554 | .993 | >.999 | >.999 | .008 | .802 | .027 | .603 | — |
| rh | 33pr | −.019 | .799 | .838 | >.999 | >.999 | −.027 | .405 | .993 | >.999 | >.999 | −.012 | .609 | −.024 | .485 | — |
| rh | a24pr | .027 | .202 | .418 | >.999 | >.999 | .030 | .570 | .993 | >.999 | >.999 | .019 | .547 | .024 | .659 | — |
| rh | p32pr | −.017 | .673 | .734 | >.999 | >.999 | −.002 | .973 | .993 | >.999 | >.999 | −.018 | .616 | .004 | .953 | — |
| rh | a24 | .000 | .496 | .601 | >.999 | >.999 | −.017 | .772 | .993 | >.999 | >.999 | .006 | .857 | −.019 | .755 | — |
| rh | d32 | .063 | .055 | .334 | .904 | >.999 | −.015 | .818 | .993 | >.999 | >.999 | .075 | **.049** | −.039 | .571 | Matching |
| rh | 8BM | .086 | **.029** | .334 | .609 | >.999 | −.007 | .928 | .993 | >.999 | >.999 | .097 | **.024** | −.037 | .638 | Matching |
| rh | p32 | .023 | .228 | .427 | >.999 | >.999 | −.009 | .853 | .993 | >.999 | >.999 | .029 | .344 | −.018 | .729 | — |
| rh | 10r | .035 | .144 | .392 | .999 | >.999 | .024 | .652 | .993 | >.999 | >.999 | .031 | .346 | .015 | .793 | — |
| rh | 47m | .017 | .287 | .442 | >.999 | >.999 | .057 | .255 | .993 | >.999 | >.999 | −.001 | .960 | .057 | .267 | — |
| rh | 8Av | .125 | **.008** | .279 | .151 | .999 | .090 | .310 | .993 | >.999 | >.999 | .107 | **.030** | .056 | .538 | Matching |
| rh | 8Ad | .105 | **.008** | .279 | .342 | >.999 | .041 | .568 | .993 | >.999 | >.999 | .102 | **.013** | .009 | .902 | Matching |
| rh | 9m | .056 | .122 | .361 | .951 | >.999 | .024 | .774 | .993 | >.999 | >.999 | .054 | .247 | .007 | .935 | — |
| rh | 8BL | .131 | **.005** | .279 | .113 | .994 | .032 | .712 | .993 | >.999 | >.999 | .134 | **.005** | −.010 | .906 | Matching |
| rh | 9p | .078 | **.022** | .334 | .724 | >.999 | .067 | .285 | .993 | >.999 | >.999 | .063 | .089 | .048 | .465 | — |
| rh | 10d | .074 | **.017** | .334 | .781 | >.999 | .048 | .401 | .993 | >.999 | >.999 | .066 | **.049** | .027 | .644 | Matching |
| rh | 8C | .052 | .153 | .396 | .973 | >.999 | .106 | .232 | .993 | >.999 | >.999 | .020 | .675 | .100 | .276 | — |
| rh | 44ᵃ | .147 | **.001** | .234 | **.047** | .934 | .083 | .329 | .993 | >.999 | >.999 | .135 | **.004** | .040 | .646 | Matching |
| rh | 45 | .082 | **.040** | .334 | .669 | >.999 | .032 | .691 | .993 | >.999 | >.999 | .080 | .074 | .007 | .934 | — |
| rh | 47l | .078 | **.020** | .334 | .726 | >.999 | .013 | .838 | .993 | >.999 | >.999 | .082 | **.026** | −.012 | .854 | Matching |
| rh | a47r | .043 | .128 | .365 | .992 | >.999 | −.009 | .882 | .993 | >.999 | >.999 | .051 | .168 | −.025 | .698 | — |
| rh | 6r | .070 | .066 | .334 | .831 | >.999 | .155 | **.049** | .993 | .869 | .871 | .024 | .594 | .147 | .070 | — |
| rh | IFJa | −.006 | .569 | .654 | >.999 | >.999 | −.031 | .555 | .993 | >.999 | >.999 | .004 | .901 | −.033 | .549 | — |
| rh | IFJp | −.067 | .997 | .997 | >.999 | >.999 | −.018 | .659 | .993 | >.999 | >.999 | −.068 | **.008** | .004 | .931 | Matching |
| rh | IFSp | .043 | .163 | .396 | .992 | >.999 | .129 | .090 | .993 | .996 | .996 | .003 | .947 | .128 | .103 | — |
| rh | IFSa | .047 | .149 | .395 | .985 | >.999 | .019 | .813 | .993 | >.999 | >.999 | .046 | .308 | .005 | .954 | — |
| rh | p9-46v | .081 | .057 | .334 | .679 | >.999 | .041 | .647 | .993 | >.999 | >.999 | .076 | .119 | .017 | .853 | — |
| rh | 46 | .081 | **.036** | .334 | .682 | >.999 | .039 | .614 | .993 | >.999 | >.999 | .077 | .077 | .015 | .854 | — |
| rh | a9-46v | .063 | .070 | .334 | .907 | >.999 | .002 | .974 | .993 | >.999 | >.999 | .069 | .088 | −.019 | .796 | — |
| rh | 9-46d | .014 | .366 | .499 | >.999 | >.999 | −.055 | .464 | .993 | >.999 | >.999 | .035 | .414 | −.066 | .395 | — |
| rh | 9a | .090 | **.021** | .334 | .541 | >.999 | .018 | .810 | .993 | >.999 | >.999 | .094 | **.028** | −.011 | .877 | Matching |
| rh | 10v | .065 | **.022** | .334 | .885 | >.999 | −.000 | .995 | .997 | >.999 | >.999 | .072 | **.020** | −.023 | .665 | Matching |
| rh | a10p | .037 | .128 | .365 | .998 | >.999 | −.008 | .880 | .993 | >.999 | >.999 | .044 | .175 | −.022 | .694 | — |
| rh | 10pp | .079 | **.008** | .279 | .713 | >.999 | .010 | .843 | .993 | >.999 | >.999 | .084 | **.007** | −.016 | .768 | Matching |
| rh | 11l | .035 | .177 | .404 | .999 | >.999 | −.027 | .678 | .993 | >.999 | >.999 | .048 | .196 | −.042 | .533 | — |
| rh | 13l | .026 | .213 | .420 | >.999 | >.999 | .036 | .492 | .993 | >.999 | >.999 | .016 | .622 | .032 | .564 | — |
| rh | OFC | .032 | .187 | .404 | >.999 | >.999 | −.044 | .456 | .993 | >.999 | >.999 | .051 | .152 | −.060 | .327 | — |
| rh | 47s | .024 | .246 | .441 | >.999 | >.999 | −.006 | .920 | .993 | >.999 | >.999 | .029 | .416 | −.015 | .813 | — |
| rh | LIPd | −.020 | .791 | .833 | >.999 | >.999 | −.086 | **.021** | .993 | >.999 | >.999 | .007 | .760 | −.088 | **.022** | AK− |
| rh | 6a | .030 | .218 | .425 | >.999 | >.999 | .005 | .937 | .993 | >.999 | >.999 | .032 | .403 | −.005 | .945 | — |
| rh | i6-8 | .041 | .156 | .396 | .994 | >.999 | −.004 | .948 | .993 | >.999 | >.999 | .047 | .239 | −.019 | .789 | — |
| rh | s6-8 | .047 | .109 | .359 | .984 | >.999 | .054 | .411 | .993 | >.999 | >.999 | .034 | .363 | .043 | .520 | — |
| rh | 43 | .034 | .176 | .404 | .999 | >.999 | .029 | .642 | .993 | >.999 | >.999 | .028 | .445 | .020 | .756 | — |
| rh | OP4 | .046 | .104 | .351 | .987 | >.999 | .039 | .535 | .993 | >.999 | >.999 | .038 | .291 | .027 | .675 | — |
| rh | OP1 | .014 | .335 | .483 | >.999 | >.999 | .031 | .569 | .993 | >.999 | >.999 | .005 | .883 | .030 | .600 | — |
| rh | OP2-3 | −.044 | .946 | .959 | >.999 | >.999 | −.071 | .108 | .993 | >.999 | >.999 | −.024 | .388 | −.063 | .164 | — |
| rh | 52 | .022 | .199 | .418 | >.999 | >.999 | .016 | .696 | .993 | >.999 | >.999 | .019 | .462 | .010 | .814 | — |
| rh | RI | −.011 | .667 | .729 | >.999 | >.999 | .013 | .732 | .993 | >.999 | >.999 | −.016 | .508 | .018 | .646 | — |
| rh | PFcm | .054 | .065 | .334 | .963 | >.999 | .049 | .402 | .993 | >.999 | >.999 | .043 | .216 | .036 | .554 | — |
| rh | PoI2 | −.040 | .859 | .883 | >.999 | >.999 | −.052 | .416 | .993 | >.999 | >.999 | −.027 | .478 | −.044 | .508 | — |
| rh | TA2 | .014 | .348 | .488 | >.999 | >.999 | −.057 | .344 | .993 | >.999 | >.999 | .035 | .324 | −.068 | .273 | — |
| rh | FOP4 | .062 | **.039** | .334 | .917 | >.999 | .039 | .499 | .993 | >.999 | >.999 | .055 | .107 | .022 | .710 | — |
| rh | MI | .031 | .209 | .418 | >.999 | >.999 | −.003 | .962 | .993 | >.999 | >.999 | .035 | .350 | −.014 | .832 | — |
| rh | Pir | −.002 | .520 | .616 | >.999 | >.999 | −.067 | .245 | .993 | >.999 | >.999 | .021 | .537 | −.073 | .216 | — |
| rh | AVI | .032 | .161 | .396 | >.999 | >.999 | −.032 | .560 | .993 | >.999 | >.999 | .047 | .148 | −.047 | .410 | — |
| rh | AAIC | .041 | .120 | .361 | .994 | >.999 | .004 | .943 | .993 | >.999 | >.999 | .044 | .197 | −.010 | .868 | — |
| rh | FOP1 | −.032 | .840 | .874 | >.999 | >.999 | −.064 | .222 | .993 | >.999 | >.999 | −.013 | .685 | −.060 | .268 | — |
| rh | FOP3 | .007 | .394 | .521 | >.999 | >.999 | .005 | .893 | .993 | >.999 | >.999 | .006 | .822 | .003 | .931 | — |
| rh | FOP2 | .016 | .278 | .441 | >.999 | >.999 | .077 | .073 | .993 | >.999 | >.999 | −.009 | .737 | .080 | .071 | — |
| rh | PFt | .027 | .225 | .425 | >.999 | >.999 | −.117 | **.048** | .993 | >.999 | >.999 | .071 | **.044** | −.139 | **.022** | Match + AK− |
| rh | AIP | .035 | .170 | .402 | .999 | >.999 | .092 | .135 | .993 | >.999 | >.999 | .007 | .849 | .090 | .157 | — |
| rh | EC | .021 | .299 | .446 | >.999 | >.999 | −.021 | .754 | .993 | >.999 | >.999 | .031 | .435 | −.031 | .658 | — |
| rh | PreS | .032 | .162 | .396 | >.999 | >.999 | .021 | .701 | .993 | >.999 | >.999 | .028 | .384 | .013 | .828 | — |
| rh | H | −.013 | .648 | .718 | >.999 | >.999 | −.046 | .395 | .993 | >.999 | >.999 | .001 | .969 | −.046 | .406 | — |
| rh | ProS | .020 | .255 | .441 | >.999 | >.999 | −.004 | .941 | .993 | >.999 | >.999 | .023 | .447 | −.011 | .827 | — |
| rh | PeEc | .038 | .181 | .404 | .997 | >.999 | .060 | .413 | .993 | >.999 | >.999 | .022 | .599 | .053 | .480 | — |
| rh | STGa | −.011 | .618 | .695 | >.999 | >.999 | .003 | .955 | .993 | >.999 | >.999 | −.014 | .694 | .008 | .903 | — |
| rh | PBelt | .058 | **.042** | .334 | .942 | >.999 | −.031 | .569 | .993 | >.999 | >.999 | .075 | **.023** | −.055 | .334 | Matching |
| rh | A5 | .061 | .104 | .351 | .921 | >.999 | −.003 | .973 | .993 | >.999 | >.999 | .069 | .147 | −.024 | .782 | — |
| rh | PHA1 | .036 | .186 | .404 | .998 | >.999 | −.005 | .948 | .993 | >.999 | >.999 | .041 | .287 | −.018 | .800 | — |
| rh | PHA3 | −.001 | .506 | .607 | >.999 | >.999 | .034 | .423 | .993 | >.999 | >.999 | −.012 | .646 | .038 | .389 | — |
| rh | STSda | .034 | .205 | .418 | .999 | >.999 | −.019 | .798 | .993 | >.999 | >.999 | .044 | .282 | −.033 | .665 | — |
| rh | STSdp | .101 | **.008** | .279 | .388 | >.999 | .161 | **.023** | .993 | .811 | .813 | .056 | .171 | .143 | .052 | — |
| rh | STSvp | .122 | **.006** | .279 | .171 | >.999 | .066 | .412 | .993 | >.999 | >.999 | .112 | **.014** | .031 | .706 | Matching |
| rh | TGd | .068 | .075 | .334 | .854 | >.999 | .081 | .321 | .993 | >.999 | >.999 | .048 | .298 | .066 | .431 | — |
| rh | TE1a | .065 | .070 | .334 | .888 | >.999 | .084 | .259 | .993 | >.999 | >.999 | .043 | .314 | .071 | .355 | — |
| rh | TE1p | .046 | .158 | .396 | .988 | >.999 | .055 | .493 | .993 | >.999 | >.999 | .032 | .484 | .045 | .586 | — |
| rh | TE2a | .082 | **.033** | .334 | .674 | >.999 | .044 | .555 | .993 | >.999 | >.999 | .075 | .074 | .020 | .792 | — |
| rh | TF | −.000 | .490 | .596 | >.999 | >.999 | −.024 | .735 | .993 | >.999 | >.999 | .008 | .844 | −.027 | .713 | — |
| rh | TE2p | .024 | .264 | .441 | >.999 | >.999 | −.028 | .681 | .993 | >.999 | >.999 | .037 | .342 | −.039 | .570 | — |
| rh | PHT | .089 | **.030** | .334 | .562 | >.999 | .119 | .141 | .993 | >.999 | >.999 | .057 | .215 | .101 | .226 | — |
| rh | PH | .077 | **.019** | .334 | .737 | >.999 | .137 | **.027** | .993 | .983 | .984 | .038 | .286 | .125 | **.049** | AK+ |
| rh | TPOJ1 | .119 | **.010** | .281 | .188 | >.999 | .087 | .324 | .993 | >.999 | >.999 | .102 | **.037** | .054 | .550 | Matching |
| rh | TPOJ2 | .008 | .424 | .550 | >.999 | >.999 | −.020 | .801 | .993 | >.999 | >.999 | .016 | .729 | −.025 | .758 | — |
| rh | TPOJ3 | .052 | .054 | .334 | .972 | >.999 | .113 | **.030** | .993 | >.999 | >.999 | .018 | .564 | .107 | **.045** | AK+ |
| rh | DVT | .018 | .324 | .470 | >.999 | >.999 | .035 | .607 | .993 | >.999 | >.999 | .008 | .848 | .032 | .640 | — |
| rh | PGp | .032 | .208 | .418 | >.999 | >.999 | .041 | .540 | .993 | >.999 | >.999 | .021 | .586 | .035 | .620 | — |
| rh | IP2 | .078 | **.020** | .334 | .733 | >.999 | −.035 | .582 | .993 | >.999 | >.999 | .098 | **.006** | −.065 | .308 | Matching |
| rh | IP1 | .041 | .139 | .384 | .995 | >.999 | .053 | .397 | .993 | >.999 | >.999 | .027 | .477 | .045 | .487 | — |
| rh | IP0 | .027 | .224 | .425 | >.999 | >.999 | .139 | **.021** | .993 | .975 | .977 | −.018 | .615 | .145 | **.020** | AK+ |
| rh | PFop | .025 | .238 | .435 | >.999 | >.999 | .004 | .950 | .993 | >.999 | >.999 | .026 | .455 | −.005 | .941 | — |
| rh | PF | .087 | **.022** | .334 | .584 | >.999 | .056 | .439 | .993 | >.999 | >.999 | .077 | .062 | .032 | .667 | — |
| rh | PFm | .079 | **.046** | .334 | .718 | >.999 | .108 | .175 | .993 | >.999 | >.999 | .050 | .276 | .093 | .261 | — |
| rh | PGi | .114 | **.009** | .279 | .235 | >.999 | .137 | .091 | .993 | .981 | .982 | .079 | .087 | .113 | .178 | — |
| rh | PGs | .034 | .209 | .418 | .999 | >.999 | .069 | .338 | .993 | >.999 | >.999 | .013 | .750 | .065 | .381 | — |
| rh | V6A | .017 | .277 | .441 | >.999 | >.999 | .023 | .629 | .993 | >.999 | >.999 | .011 | .710 | .020 | .688 | — |
| rh | VMV1 | .033 | .182 | .404 | >.999 | >.999 | −.066 | .282 | .993 | >.999 | >.999 | .059 | .094 | −.084 | .181 | — |
| rh | VMV3 | .002 | .471 | .577 | >.999 | >.999 | −.047 | .336 | .993 | >.999 | >.999 | .018 | .536 | −.052 | .296 | — |
| rh | PHA2 | −.073 | .997 | .997 | >.999 | >.999 | −.101 | **.021** | .993 | >.999 | >.999 | −.045 | .100 | −.087 | .051 | — |
| rh | V4t | .001 | .481 | .587 | >.999 | >.999 | −.027 | .596 | .993 | >.999 | >.999 | .011 | .731 | −.031 | .559 | — |
| rh | FST | −.022 | .741 | .791 | >.999 | >.999 | −.048 | .405 | .993 | >.999 | >.999 | −.008 | .819 | −.046 | .439 | — |
| rh | V3CD | .012 | .352 | .488 | >.999 | >.999 | .039 | .463 | .993 | >.999 | >.999 | −.001 | .985 | .039 | .477 | — |
| rh | LO3 | −.017 | .686 | .744 | >.999 | >.999 | −.060 | .318 | .993 | >.999 | >.999 | .002 | .966 | −.060 | .327 | — |
| rh | VMV2 | .005 | .428 | .554 | >.999 | >.999 | .045 | .288 | .993 | >.999 | >.999 | −.011 | .698 | .048 | .267 | — |
| rh | 31pd | .041 | .105 | .351 | .994 | >.999 | .031 | .577 | .993 | >.999 | >.999 | .035 | .277 | .020 | .732 | — |
| rh | 31a | .026 | .225 | .425 | >.999 | >.999 | −.002 | .968 | .993 | >.999 | >.999 | .030 | .392 | −.012 | .845 | — |
| rh | VVC | .048 | .112 | .361 | .983 | >.999 | .103 | .121 | .993 | >.999 | >.999 | .017 | .652 | .098 | .154 | — |
| rh | 25 | −.004 | .560 | .646 | >.999 | >.999 | −.004 | .927 | .993 | >.999 | >.999 | −.003 | .908 | −.003 | .946 | — |
| rh | s32 | .026 | .164 | .397 | >.999 | >.999 | −.011 | .797 | .993 | >.999 | >.999 | .032 | .217 | −.021 | .623 | — |
| rh | pOFC | .027 | .216 | .422 | >.999 | >.999 | .020 | .724 | .993 | >.999 | >.999 | .023 | .495 | .013 | .824 | — |
| rh | PoI1 | .021 | .255 | .441 | >.999 | >.999 | .077 | .129 | .993 | >.999 | >.999 | −.004 | .895 | .079 | .133 | — |
| rh | Ig | .019 | .234 | .431 | >.999 | >.999 | .029 | .478 | .993 | >.999 | >.999 | .011 | .668 | .025 | .548 | — |
| rh | FOP5 | −.004 | .540 | .635 | >.999 | >.999 | .015 | .790 | .993 | >.999 | >.999 | −.010 | .768 | .018 | .759 | — |
| rh | p10p | .032 | .187 | .404 | >.999 | >.999 | −.003 | .957 | .993 | >.999 | >.999 | .036 | .306 | −.014 | .810 | — |
| rh | p47r | .057 | .076 | .334 | .950 | >.999 | .017 | .790 | .993 | >.999 | >.999 | .057 | .134 | −.000 | .997 | — |
| rh | TGv | .010 | .393 | .521 | >.999 | >.999 | .008 | .911 | .993 | >.999 | >.999 | .009 | .828 | .005 | .943 | — |
| rh | MBelt | −.016 | .701 | .758 | >.999 | >.999 | −.036 | .453 | .993 | >.999 | >.999 | −.005 | .865 | −.035 | .488 | — |
| rh | LBelt | .055 | **.045** | .334 | .957 | >.999 | .038 | .477 | .993 | >.999 | >.999 | .048 | .134 | .023 | .678 | — |
| rh | A4 | .013 | .376 | .507 | >.999 | >.999 | −.060 | .418 | .993 | >.999 | >.999 | .035 | .406 | −.071 | .351 | — |
| rh | STSva | .037 | .180 | .404 | .997 | >.999 | .025 | .721 | .993 | >.999 | >.999 | .033 | .423 | .015 | .840 | — |
| rh | TE1m | .076 | **.043** | .334 | .761 | >.999 | .066 | .370 | .993 | >.999 | >.999 | .061 | .149 | .047 | .538 | — |
| rh | PI | .020 | .279 | .441 | >.999 | >.999 | −.063 | .265 | .993 | >.999 | >.999 | .044 | .196 | −.077 | .187 | — |
| rh | a32pr | .019 | .288 | .442 | >.999 | >.999 | −.020 | .738 | .993 | >.999 | >.999 | .028 | .415 | −.029 | .636 | — |
| rh | p24 | −.040 | .863 | .883 | >.999 | >.999 | −.005 | .932 | .993 | >.999 | >.999 | −.042 | .238 | .008 | .902 | — |

**Prospective ELS — Adjusted for sex, ART, age, SES (N = 89, 20,000 permutations, 360 parcels, r(|a − b|, mean) = .31)**

|  | | **Matching ρ(D, \|a − b\|)** | | | | | **Additive ρ(D, mean(a, b))** | | | | | **Joint model D ~ \|a − b\| + mean(a, b)** | | | |  |
| --- | --- | --- | --- | --- | --- | --- | --- | --- | --- | --- | --- | --- | --- | --- | --- | --- |
| **Hemi** | **Region** | **ρ** | **p** | **q** | **p FWE-P** | **p FWE-F** | **ρ** | **p** | **q** | **p FWE-P** | **p FWE-F** | **β \|a−b\|** | **p** | **β mean** | **p** | **Mechanism** |
| lh | V1 | .059 | .097 | .320 | .934 | >.999 | −.005 | .947 | .990 | >.999 | >.999 | .067 | .128 | −.027 | .748 | — |
| lh | MST | .005 | .425 | .546 | >.999 | >.999 | .032 | .397 | .990 | >.999 | >.999 | −.006 | .803 | .034 | .385 | — |
| lh | V6 | .020 | .272 | .431 | >.999 | >.999 | −.019 | .740 | .990 | >.999 | >.999 | .029 | .382 | −.028 | .633 | — |
| lh | V2 | .051 | .137 | .373 | .973 | >.999 | −.030 | .710 | .990 | >.999 | >.999 | .067 | .141 | −.051 | .540 | — |
| lh | V3 | .030 | .249 | .431 | >.999 | >.999 | −.025 | .750 | .990 | >.999 | >.999 | .042 | .344 | −.038 | .631 | — |
| lh | V4 | −.004 | .528 | .617 | >.999 | >.999 | −.053 | .505 | .990 | >.999 | >.999 | .014 | .758 | −.058 | .486 | — |
| lh | V8 | .018 | .322 | .468 | >.999 | >.999 | .002 | .973 | .990 | >.999 | >.999 | .019 | .629 | −.004 | .961 | — |
| lh | 4 | .046 | .161 | .379 | .987 | >.999 | −.021 | .797 | .990 | >.999 | >.999 | .059 | .201 | −.040 | .635 | — |
| lh | 3b | .037 | .184 | .397 | .998 | >.999 | .005 | .947 | .990 | >.999 | >.999 | .039 | .336 | −.008 | .918 | — |
| lh | FEF | .050 | .101 | .325 | .976 | >.999 | −.001 | .990 | .990 | >.999 | >.999 | .056 | .147 | −.018 | .788 | — |
| lh | PEF | −.024 | .757 | .804 | >.999 | >.999 | .013 | .817 | .990 | >.999 | >.999 | −.032 | .355 | .023 | .697 | — |
| lh | 55b | .037 | .205 | .414 | .997 | >.999 | −.027 | .725 | .990 | >.999 | >.999 | .051 | .246 | −.043 | .589 | — |
| lh | V3A | −.036 | .848 | .873 | >.999 | >.999 | −.046 | .427 | .990 | >.999 | >.999 | −.024 | .492 | −.039 | .517 | — |
| lh | RSC | −.026 | .782 | .823 | >.999 | >.999 | −.105 | **.048** | .990 | >.999 | >.999 | .008 | .795 | −.108 | **.049** | AK− |
| lh | POS2 | .062 | .070 | .309 | .910 | >.999 | .053 | .459 | .990 | >.999 | >.999 | .051 | .217 | .037 | .616 | — |
| lh | V7 | .015 | .294 | .434 | >.999 | >.999 | .057 | .221 | .990 | >.999 | >.999 | −.003 | .921 | .057 | .228 | — |
| lh | IPS1 | −.010 | .591 | .669 | >.999 | >.999 | −.071 | .287 | .990 | >.999 | >.999 | .014 | .719 | −.075 | .271 | — |
| lh | FFC | .062 | .092 | .319 | .913 | >.999 | −.056 | .489 | .990 | >.999 | >.999 | .088 | .051 | −.083 | .314 | — |
| lh | V3B | .046 | **.048** | .308 | .988 | >.999 | −.027 | .545 | .990 | >.999 | >.999 | .060 | **.028** | −.046 | .318 | Matching |
| lh | LO1 | −.001 | .507 | .604 | >.999 | >.999 | −.021 | .694 | .990 | >.999 | >.999 | .006 | .854 | −.022 | .677 | — |
| lh | LO2 | .042 | .127 | .363 | .994 | >.999 | .015 | .807 | .990 | >.999 | >.999 | .041 | .256 | .002 | .972 | — |
| lh | PIT | .049 | .105 | .327 | .980 | >.999 | .003 | .966 | .990 | >.999 | >.999 | .053 | .169 | −.014 | .844 | — |
| lh | MT | −.029 | .849 | .873 | >.999 | >.999 | −.034 | .449 | .990 | >.999 | >.999 | −.021 | .457 | −.028 | .549 | — |
| lh | A1 | .020 | .267 | .431 | >.999 | >.999 | −.058 | .273 | .990 | >.999 | >.999 | .042 | .188 | −.071 | .191 | — |
| lh | PSL | .000 | .495 | .594 | >.999 | >.999 | −.031 | .709 | .990 | >.999 | >.999 | .011 | .819 | −.034 | .686 | — |
| lh | SFL | .078 | .053 | .308 | .726 | >.999 | .091 | .254 | .990 | >.999 | >.999 | .054 | .236 | .074 | .368 | — |
| lh | PCV | .020 | .284 | .431 | >.999 | >.999 | −.018 | .767 | .990 | >.999 | >.999 | .029 | .426 | −.027 | .669 | — |
| lh | STV | .056 | .124 | .358 | .952 | >.999 | .048 | .568 | .990 | >.999 | >.999 | .046 | .335 | .034 | .695 | — |
| lh | 7Pm | .028 | .214 | .415 | >.999 | >.999 | −.029 | .617 | .990 | >.999 | >.999 | .041 | .234 | −.042 | .481 | — |
| lh | 7m | .044 | .152 | .378 | .990 | >.999 | .058 | .441 | .990 | >.999 | >.999 | .029 | .489 | .049 | .531 | — |
| lh | POS1 | .000 | .488 | .590 | >.999 | >.999 | −.053 | .426 | .990 | >.999 | >.999 | .019 | .625 | −.059 | .390 | — |
| lh | 23d | .029 | .219 | .415 | >.999 | >.999 | .078 | .200 | .990 | >.999 | >.999 | .005 | .901 | .076 | .221 | — |
| lh | v23ab | .043 | .081 | .309 | .992 | >.999 | .074 | .153 | .990 | >.999 | >.999 | .022 | .469 | .067 | .210 | — |
| lh | d23ab | .015 | .350 | .486 | >.999 | >.999 | .007 | .914 | .990 | >.999 | >.999 | .014 | .725 | .003 | .968 | — |
| lh | 31pv | .090 | **.004** | .249 | .537 | >.999 | .130 | **.017** | .990 | .995 | .995 | .055 | .095 | .112 | **.046** | AK+ |
| lh | 5m | .011 | .354 | .486 | >.999 | >.999 | .081 | .101 | .990 | >.999 | >.999 | −.016 | .600 | .086 | .090 | — |
| lh | 5mv | .053 | **.044** | .308 | .964 | >.999 | .016 | .748 | .990 | >.999 | >.999 | .054 | .080 | −.000 | .995 | — |
| lh | 23c | .046 | .084 | .309 | .988 | >.999 | −.032 | .567 | .990 | >.999 | >.999 | .062 | .058 | −.051 | .367 | — |
| lh | 5L | .024 | .235 | .423 | >.999 | >.999 | −.004 | .939 | .990 | >.999 | >.999 | .028 | .403 | −.013 | .824 | — |
| lh | 24dd | .035 | .179 | .390 | .998 | >.999 | .017 | .799 | .990 | >.999 | >.999 | .033 | .384 | .007 | .922 | — |
| lh | 24dv | .057 | **.037** | .308 | .945 | >.999 | .005 | .920 | .990 | >.999 | >.999 | .062 | **.045** | −.014 | .793 | Matching |
| lh | 7AL | −.013 | .637 | .705 | >.999 | >.999 | .001 | .988 | .990 | >.999 | >.999 | −.015 | .685 | .005 | .928 | — |
| lh | SCEF | .026 | .285 | .431 | >.999 | >.999 | .004 | .954 | .990 | >.999 | >.999 | .027 | .544 | −.004 | .959 | — |
| lh | 6ma | .026 | .270 | .431 | >.999 | >.999 | −.022 | .761 | .990 | >.999 | >.999 | .037 | .392 | −.034 | .659 | — |
| lh | 7Am | .013 | .371 | .494 | >.999 | >.999 | .035 | .606 | .990 | >.999 | >.999 | .002 | .957 | .034 | .619 | — |
| lh | 7Pl | .022 | .267 | .431 | >.999 | >.999 | .044 | .460 | .990 | >.999 | >.999 | .009 | .791 | .041 | .500 | — |
| lh | 7PC | .031 | .225 | .418 | >.999 | >.999 | .063 | .358 | .990 | >.999 | >.999 | .012 | .757 | .059 | .399 | — |
| lh | LIPv | −.003 | .523 | .614 | >.999 | >.999 | .040 | .530 | .990 | >.999 | >.999 | −.018 | .644 | .046 | .489 | — |
| lh | VIP | .028 | .241 | .429 | >.999 | >.999 | .076 | .270 | .990 | >.999 | >.999 | .005 | .909 | .074 | .293 | — |
| lh | MIP | .011 | .366 | .493 | >.999 | >.999 | −.025 | .688 | .990 | >.999 | >.999 | .022 | .554 | −.032 | .621 | — |
| lh | 1 | .045 | .154 | .378 | .989 | >.999 | −.006 | .941 | .990 | >.999 | >.999 | .052 | .228 | −.022 | .781 | — |
| lh | 2 | .071 | **.040** | .308 | .812 | >.999 | .058 | .388 | .990 | >.999 | >.999 | .059 | .131 | .040 | .563 | — |
| lh | 3a | −.010 | .603 | .675 | >.999 | >.999 | −.084 | .147 | .990 | >.999 | >.999 | .018 | .598 | −.090 | .133 | — |
| lh | 6d | .006 | .432 | .550 | >.999 | >.999 | .001 | .986 | .990 | >.999 | >.999 | .007 | .863 | −.001 | .989 | — |
| lh | 6mp | .054 | .063 | .308 | .961 | >.999 | .036 | .551 | .990 | >.999 | >.999 | .048 | .175 | .021 | .737 | — |
| lh | 6v | .001 | .478 | .590 | >.999 | >.999 | .026 | .703 | .990 | >.999 | >.999 | −.008 | .849 | .028 | .689 | — |
| lh | p24pr | .046 | .099 | .320 | .988 | >.999 | −.006 | .916 | .990 | >.999 | >.999 | .053 | .127 | −.023 | .709 | — |
| lh | 33pr | .006 | .409 | .532 | >.999 | >.999 | .040 | .333 | .990 | >.999 | >.999 | −.007 | .781 | .043 | .321 | — |
| lh | a24pr | .026 | .202 | .411 | >.999 | >.999 | .031 | .555 | .990 | >.999 | >.999 | .018 | .571 | .025 | .636 | — |
| lh | p32pr | .037 | .134 | .373 | .998 | >.999 | .024 | .663 | .990 | >.999 | >.999 | .032 | .316 | .013 | .810 | — |
| lh | a24 | .057 | .051 | .308 | .949 | >.999 | −.004 | .937 | .990 | >.999 | >.999 | .064 | .058 | −.024 | .680 | — |
| lh | d32 | .028 | .226 | .418 | >.999 | >.999 | .042 | .525 | .990 | >.999 | >.999 | .017 | .661 | .037 | .585 | — |
| lh | 8BM | .077 | .052 | .308 | .741 | >.999 | −.014 | .865 | .990 | >.999 | >.999 | .090 | **.047** | −.042 | .613 | Matching |
| lh | p32 | −.003 | .546 | .632 | >.999 | >.999 | .037 | .391 | .990 | >.999 | >.999 | −.016 | .552 | .042 | .345 | — |
| lh | 10r | .045 | .079 | .309 | .989 | >.999 | .035 | .498 | .990 | >.999 | >.999 | .038 | .230 | .023 | .666 | — |
| lh | 47m | .019 | .269 | .431 | >.999 | >.999 | .045 | .370 | .990 | >.999 | >.999 | .005 | .869 | .044 | .401 | — |
| lh | 8Av | .092 | **.033** | .308 | .509 | >.999 | .078 | .352 | .990 | >.999 | >.999 | .075 | .112 | .054 | .529 | — |
| lh | 8Ad | .005 | .445 | .558 | >.999 | >.999 | .046 | .489 | .990 | >.999 | >.999 | −.011 | .783 | .050 | .471 | — |
| lh | 9m | .049 | .154 | .378 | .980 | >.999 | .018 | .825 | .990 | >.999 | >.999 | .048 | .300 | .003 | .969 | — |
| lh | 8BL | .097 | **.013** | .308 | .434 | >.999 | .038 | .603 | .990 | >.999 | >.999 | .095 | **.021** | .008 | .912 | Matching |
| lh | 9p | .030 | .225 | .418 | >.999 | >.999 | .062 | .356 | .990 | >.999 | >.999 | .011 | .775 | .059 | .396 | — |
| lh | 10d | .051 | .072 | .309 | .972 | >.999 | .026 | .650 | .990 | >.999 | >.999 | .048 | .159 | .011 | .854 | — |
| lh | 8C | .138 | **.003** | .249 | .079 | .978 | .150 | .068 | .990 | .915 | .917 | .101 | **.027** | .118 | .161 | Matching |
| lh | 44 | .090 | **.034** | .308 | .543 | >.999 | .125 | .132 | .990 | .998 | .998 | .056 | .236 | .107 | .209 | — |
| lh | 45 | .077 | .066 | .309 | .730 | >.999 | −.019 | .832 | .990 | >.999 | >.999 | .092 | .059 | −.048 | .600 | — |
| lh | 47l | .070 | .058 | .308 | .832 | >.999 | −.022 | .771 | .990 | >.999 | >.999 | .085 | **.043** | −.048 | .526 | Matching |
| lh | a47r | .073 | **.035** | .308 | .785 | >.999 | .048 | .472 | .990 | >.999 | >.999 | .065 | .098 | .028 | .684 | — |
| lh | 6r | .062 | .084 | .309 | .908 | >.999 | .123 | .110 | .990 | .999 | >.999 | .026 | .545 | .115 | .147 | — |
| lh | IFJa | −.023 | .714 | .769 | >.999 | >.999 | .063 | .342 | .990 | >.999 | >.999 | −.047 | .221 | .077 | .253 | — |
| lh | IFJp | .023 | .248 | .431 | >.999 | >.999 | .061 | .266 | .990 | >.999 | >.999 | .004 | .895 | .060 | .293 | — |
| lh | IFSp | −.017 | .644 | .709 | >.999 | >.999 | .026 | .727 | .990 | >.999 | >.999 | −.028 | .520 | .035 | .652 | — |
| lh | IFSaᵃ | .181 | **<.001** | .108 | **.006** | .555 | .300 | **<.001** | .234 | **.009** | **.009** | .097 | **.045** | .269 | **.002** | Match + AK+ |
| lh | p9-46v | .059 | .118 | .347 | .936 | >.999 | .032 | .701 | .990 | >.999 | >.999 | .054 | .266 | .016 | .858 | — |
| lh | 46 | .035 | .206 | .414 | .998 | >.999 | .002 | .982 | .990 | >.999 | >.999 | .039 | .365 | −.011 | .888 | — |
| lh | a9-46v | .056 | .095 | .319 | .950 | >.999 | .059 | .415 | .990 | >.999 | >.999 | .042 | .320 | .046 | .536 | — |
| lh | 9-46d | .026 | .279 | .431 | >.999 | >.999 | −.004 | .961 | .990 | >.999 | >.999 | .030 | .499 | −.013 | .871 | — |
| lh | 9a | .069 | .070 | .309 | .841 | >.999 | −.021 | .794 | .990 | >.999 | >.999 | .084 | .061 | −.047 | .561 | — |
| lh | 10v | .057 | **.048** | .308 | .947 | >.999 | .012 | .828 | .990 | >.999 | >.999 | .059 | .076 | −.006 | .919 | — |
| lh | a10p | .005 | .437 | .552 | >.999 | >.999 | −.022 | .682 | .990 | >.999 | >.999 | .013 | .692 | −.026 | .634 | — |
| lh | 10pp | .045 | .072 | .309 | .990 | >.999 | .023 | .636 | .990 | >.999 | >.999 | .042 | .164 | .010 | .841 | — |
| lh | 11l | .038 | .154 | .378 | .997 | >.999 | −.018 | .768 | .990 | >.999 | >.999 | .048 | .184 | −.034 | .602 | — |
| lh | 13l | .025 | .246 | .429 | >.999 | >.999 | .036 | .562 | .990 | >.999 | >.999 | .015 | .681 | .031 | .624 | — |
| lh | OFC | .013 | .356 | .486 | >.999 | >.999 | −.088 | .126 | .990 | >.999 | >.999 | .045 | .190 | −.102 | .084 | — |
| lh | 47s | .012 | .358 | .486 | >.999 | >.999 | −.049 | .396 | .990 | >.999 | >.999 | .031 | .367 | −.059 | .319 | — |
| lh | LIPd | .048 | .059 | .308 | .984 | >.999 | .005 | .918 | .990 | >.999 | >.999 | .051 | .086 | −.011 | .827 | — |
| lh | 6a | .040 | .143 | .373 | .996 | >.999 | .076 | .229 | .990 | >.999 | >.999 | .018 | .635 | .070 | .279 | — |
| lh | i6-8 | .042 | .122 | .353 | .994 | >.999 | .101 | .097 | .990 | >.999 | >.999 | .012 | .746 | .097 | .119 | — |
| lh | s6-8 | .047 | .083 | .309 | .987 | >.999 | .084 | .127 | .990 | >.999 | >.999 | .023 | .488 | .076 | .174 | — |
| lh | 43 | .014 | .331 | .475 | >.999 | >.999 | .063 | .238 | .990 | >.999 | >.999 | −.007 | .836 | .065 | .235 | — |
| lh | OP4 | .064 | **.050** | .308 | .892 | >.999 | .047 | .474 | .990 | >.999 | >.999 | .055 | .150 | .030 | .658 | — |
| lh | OP1 | .013 | .351 | .486 | >.999 | >.999 | .009 | .873 | .990 | >.999 | >.999 | .011 | .751 | .006 | .923 | — |
| lh | OP2-3 | −.010 | .639 | .705 | >.999 | >.999 | −.019 | .677 | .990 | >.999 | >.999 | −.005 | .853 | −.017 | .707 | — |
| lh | 52 | −.055 | .972 | .983 | >.999 | >.999 | −.085 | .065 | .990 | >.999 | >.999 | −.032 | .270 | −.076 | .113 | — |
| lh | RI | .017 | .288 | .434 | >.999 | >.999 | .007 | .881 | .990 | >.999 | >.999 | .016 | .588 | .002 | .968 | — |
| lh | PFcm | .073 | **.026** | .308 | .797 | >.999 | .052 | .400 | .990 | >.999 | >.999 | .063 | .087 | .032 | .612 | — |
| lh | PoI2 | .024 | .244 | .429 | >.999 | >.999 | −.048 | .430 | .990 | >.999 | >.999 | .044 | .219 | −.062 | .324 | — |
| lh | TA2 | .034 | .147 | .378 | .999 | >.999 | −.009 | .865 | .990 | >.999 | >.999 | .041 | .197 | −.022 | .689 | — |
| lh | FOP4 | .056 | .068 | .309 | .953 | >.999 | .133 | **.032** | .990 | .990 | .991 | .016 | .679 | .129 | **.046** | AK+ |
| lh | MI | −.022 | .715 | .769 | >.999 | >.999 | −.075 | .245 | .990 | >.999 | >.999 | .002 | .959 | −.075 | .257 | — |
| lh | Pir | −.015 | .678 | .735 | >.999 | >.999 | −.029 | .588 | .990 | >.999 | >.999 | −.007 | .823 | −.026 | .628 | — |
| lh | AVI | .052 | .084 | .309 | .971 | >.999 | .061 | .333 | .990 | >.999 | >.999 | .036 | .328 | .050 | .445 | — |
| lh | AAIC | −.020 | .723 | .774 | >.999 | >.999 | −.080 | .153 | .990 | >.999 | >.999 | .006 | .863 | −.081 | .155 | — |
| lh | FOP1 | .002 | .472 | .586 | >.999 | >.999 | .002 | .976 | .990 | >.999 | >.999 | .001 | .962 | .001 | .985 | — |
| lh | FOP3 | −.065 | .987 | .995 | >.999 | >.999 | −.017 | .712 | .990 | >.999 | >.999 | −.066 | **.018** | .004 | .934 | Matching |
| lh | FOP2 | .039 | .071 | .309 | .996 | >.999 | −.042 | .302 | .990 | >.999 | >.999 | .058 | **.027** | −.061 | .148 | Matching |
| lh | PFt | .019 | .270 | .431 | >.999 | >.999 | −.070 | .170 | .990 | >.999 | >.999 | .045 | .148 | −.084 | .113 | — |
| lh | AIP | .007 | .413 | .535 | >.999 | >.999 | −.026 | .652 | .990 | >.999 | >.999 | .017 | .626 | −.032 | .600 | — |
| lh | EC | .008 | .406 | .532 | >.999 | >.999 | −.039 | .529 | .990 | >.999 | >.999 | .023 | .536 | −.046 | .473 | — |
| lh | PreS | .022 | .271 | .431 | >.999 | >.999 | −.065 | .285 | .990 | >.999 | >.999 | .047 | .189 | −.080 | .204 | — |
| lh | H | .063 | **.035** | .308 | .904 | >.999 | .021 | .713 | .990 | >.999 | >.999 | .063 | .063 | .001 | .986 | — |
| lh | ProS | .024 | .187 | .398 | >.999 | >.999 | .054 | .206 | .990 | >.999 | >.999 | .008 | .772 | .051 | .243 | — |
| lh | PeEc | .042 | .169 | .381 | .994 | >.999 | −.016 | .832 | .990 | >.999 | >.999 | .052 | .231 | −.032 | .682 | — |
| lh | STGa | .020 | .300 | .439 | >.999 | >.999 | .038 | .568 | .990 | >.999 | >.999 | .009 | .810 | .035 | .607 | — |
| lh | PBelt | −.010 | .612 | .682 | >.999 | >.999 | −.042 | .435 | .990 | >.999 | >.999 | .004 | .900 | −.044 | .434 | — |
| lh | A5 | .101 | **.024** | .308 | .379 | >.999 | .037 | .659 | .990 | >.999 | >.999 | .099 | **.039** | .006 | .945 | Matching |
| lh | PHA1 | .022 | .253 | .431 | >.999 | >.999 | −.140 | **.016** | .990 | .973 | .974 | .074 | **.032** | −.163 | **.006** | Match + AK− |
| lh | PHA3 | .012 | .371 | .494 | >.999 | >.999 | −.018 | .779 | .990 | >.999 | >.999 | .019 | .597 | −.024 | .712 | — |
| lh | STSda | .057 | .079 | .309 | .946 | >.999 | .046 | .502 | .990 | >.999 | >.999 | .047 | .229 | .031 | .660 | — |
| lh | STSdp | .058 | .076 | .309 | .941 | >.999 | .045 | .518 | .990 | >.999 | >.999 | .049 | .219 | .029 | .682 | — |
| lh | STSvp | .051 | .140 | .373 | .971 | >.999 | .044 | .592 | .990 | >.999 | >.999 | .042 | .365 | .031 | .710 | — |
| lh | TGd | .037 | .215 | .415 | .998 | >.999 | .016 | .849 | .990 | >.999 | >.999 | .035 | .448 | .005 | .950 | — |
| lh | TE1a | .065 | .084 | .309 | .882 | >.999 | .026 | .741 | .990 | >.999 | >.999 | .063 | .158 | .006 | .942 | — |
| lh | TE1p | .029 | .277 | .431 | >.999 | >.999 | −.051 | .549 | .990 | >.999 | >.999 | .050 | .300 | −.067 | .447 | — |
| lh | TE2a | .018 | .333 | .476 | >.999 | >.999 | −.034 | .653 | .990 | >.999 | >.999 | .032 | .462 | −.044 | .573 | — |
| lh | TF | .027 | .262 | .431 | >.999 | >.999 | .003 | .970 | .990 | >.999 | >.999 | .030 | .485 | −.006 | .935 | — |
| lh | TE2p | .042 | .170 | .381 | .993 | >.999 | −.009 | .901 | .990 | >.999 | >.999 | .050 | .247 | −.025 | .748 | — |
| lh | PHT | .037 | .230 | .421 | .997 | >.999 | −.057 | .525 | .990 | >.999 | >.999 | .061 | .222 | −.076 | .407 | — |
| lh | PH | .025 | .278 | .431 | >.999 | >.999 | −.026 | .727 | .990 | >.999 | >.999 | .036 | .385 | −.037 | .624 | — |
| lh | TPOJ1 | −.012 | .597 | .671 | >.999 | >.999 | −.031 | .686 | .990 | >.999 | >.999 | −.002 | .959 | −.031 | .702 | — |
| lh | TPOJ2 | .058 | .103 | .327 | .940 | >.999 | −.005 | .950 | .990 | >.999 | >.999 | .066 | .138 | −.025 | .755 | — |
| lh | TPOJ3 | .044 | .063 | .308 | .991 | >.999 | −.033 | .478 | .990 | >.999 | >.999 | .060 | **.035** | −.051 | .280 | Matching |
| lh | DVT | .057 | .067 | .309 | .948 | >.999 | .078 | .226 | .990 | >.999 | >.999 | .036 | .331 | .067 | .314 | — |
| lh | PGp | .027 | .256 | .431 | >.999 | >.999 | .044 | .523 | .990 | >.999 | >.999 | .014 | .716 | .039 | .577 | — |
| lh | IP2 | −.025 | .766 | .811 | >.999 | >.999 | −.035 | .539 | .990 | >.999 | >.999 | −.016 | .638 | −.030 | .610 | — |
| lh | IP1 | −.006 | .565 | .647 | >.999 | >.999 | −.007 | .903 | .990 | >.999 | >.999 | −.005 | .898 | −.006 | .926 | — |
| lh | IP0 | .043 | .105 | .327 | .992 | >.999 | −.073 | .214 | .990 | >.999 | >.999 | .073 | **.029** | −.095 | .109 | Matching |
| lh | PFop | −.022 | .730 | .780 | >.999 | >.999 | .002 | .978 | .990 | >.999 | >.999 | −.025 | .482 | .009 | .881 | — |
| lh | PF | .026 | .285 | .431 | >.999 | >.999 | .006 | .937 | .990 | >.999 | >.999 | .027 | .563 | −.002 | .982 | — |
| lh | PFm | .094 | **.024** | .308 | .477 | >.999 | .108 | .179 | .990 | >.999 | >.999 | .067 | .141 | .087 | .286 | — |
| lh | PGi | .069 | .080 | .309 | .840 | >.999 | .088 | .293 | .990 | >.999 | >.999 | .046 | .330 | .074 | .393 | — |
| lh | PGsᵃ | .000 | .489 | .590 | >.999 | >.999 | .085 | .211 | .990 | >.999 | >.999 | −.029 | .454 | .094 | .178 | — |
| lh | V6A | −.032 | .877 | .894 | >.999 | >.999 | −.073 | .102 | .990 | >.999 | >.999 | −.010 | .705 | −.069 | .132 | — |
| lh | VMV1 | −.031 | .845 | .873 | >.999 | >.999 | .024 | .637 | .990 | >.999 | >.999 | −.042 | .157 | .037 | .472 | — |
| lh | VMV3 | .010 | .361 | .488 | >.999 | >.999 | .062 | .218 | .990 | >.999 | >.999 | −.010 | .740 | .066 | .209 | — |
| lh | PHA2 | .001 | .490 | .590 | >.999 | >.999 | −.024 | .526 | .990 | >.999 | >.999 | .009 | .707 | −.026 | .487 | — |
| lh | V4t | −.008 | .597 | .671 | >.999 | >.999 | −.094 | .057 | .990 | >.999 | >.999 | .024 | .426 | −.102 | **.046** | AK− |
| lh | FST | .016 | .340 | .483 | >.999 | >.999 | −.053 | .436 | .990 | >.999 | >.999 | .036 | .353 | −.064 | .359 | — |
| lh | V3CD | .037 | .142 | .373 | .998 | >.999 | .036 | .542 | .990 | >.999 | >.999 | .029 | .401 | .027 | .652 | — |
| lh | LO3 | −.073 | .993 | .997 | >.999 | >.999 | −.035 | .485 | .990 | >.999 | >.999 | −.068 | **.025** | −.014 | .791 | Matching |
| lh | VMV2 | −.032 | .889 | .904 | >.999 | >.999 | −.082 | **.041** | .990 | >.999 | >.999 | −.007 | .797 | −.080 | .054 | — |
| lh | 31pd | .018 | .265 | .431 | >.999 | >.999 | .007 | .888 | .990 | >.999 | >.999 | .018 | .545 | .001 | .978 | — |
| lh | 31a | .021 | .283 | .431 | >.999 | >.999 | −.032 | .614 | .990 | >.999 | >.999 | .035 | .344 | −.043 | .511 | — |
| lh | VVC | .054 | .098 | .320 | .962 | >.999 | .028 | .693 | .990 | >.999 | >.999 | .050 | .216 | .013 | .863 | — |
| lh | 25 | .030 | .157 | .379 | >.999 | >.999 | −.040 | .424 | .990 | >.999 | >.999 | .047 | .114 | −.054 | .281 | — |
| lh | s32 | .002 | .460 | .574 | >.999 | >.999 | −.001 | .981 | .990 | >.999 | >.999 | .002 | .908 | −.001 | .963 | — |
| lh | pOFC | .068 | **.039** | .308 | .855 | >.999 | −.008 | .902 | .990 | >.999 | >.999 | .078 | **.033** | −.032 | .619 | Matching |
| lh | PoI1 | −.005 | .549 | .634 | >.999 | >.999 | −.059 | .299 | .990 | >.999 | >.999 | .015 | .660 | −.064 | .277 | — |
| lh | Ig | .019 | .254 | .431 | >.999 | >.999 | .074 | .123 | .990 | >.999 | >.999 | −.005 | .876 | .075 | .128 | — |
| lh | FOP5 | −.035 | .811 | .846 | >.999 | >.999 | .059 | .374 | .990 | >.999 | >.999 | −.059 | .123 | .078 | .255 | — |
| lh | p10p | .034 | .156 | .379 | .999 | >.999 | −.007 | .900 | .990 | >.999 | >.999 | .040 | .220 | −.020 | .729 | — |
| lh | p47r | .108 | **.009** | .249 | .296 | >.999 | .112 | .135 | .990 | >.999 | >.999 | .081 | .062 | .087 | .260 | — |
| lh | TGv | −.014 | .627 | .697 | >.999 | >.999 | .022 | .762 | .990 | >.999 | >.999 | −.023 | .573 | .029 | .691 | — |
| lh | MBelt | .053 | .052 | .308 | .965 | >.999 | −.050 | .348 | .990 | >.999 | >.999 | .076 | **.016** | −.074 | .177 | Matching |
| lh | LBelt | .022 | .217 | .415 | >.999 | >.999 | .006 | .890 | .990 | >.999 | >.999 | .022 | .431 | −.001 | .989 | — |
| lh | A4 | .076 | .063 | .308 | .746 | >.999 | .037 | .665 | .990 | >.999 | >.999 | .072 | .128 | .014 | .866 | — |
| lh | STSva | .002 | .484 | .590 | >.999 | >.999 | −.008 | .892 | .990 | >.999 | >.999 | .004 | .897 | −.009 | .876 | — |
| lh | TE1m | .065 | .063 | .308 | .890 | >.999 | .036 | .613 | .990 | >.999 | >.999 | .059 | .141 | .017 | .817 | — |
| lh | PI | −.002 | .517 | .610 | >.999 | >.999 | −.069 | .238 | .990 | >.999 | >.999 | .022 | .522 | −.076 | .209 | — |
| lh | a32pr | .041 | .149 | .378 | .995 | >.999 | .010 | .879 | .990 | >.999 | >.999 | .042 | .278 | −.003 | .964 | — |
| lh | p24 | .006 | .426 | .546 | >.999 | >.999 | −.026 | .676 | .990 | >.999 | >.999 | .016 | .678 | −.031 | .629 | — |
| rh | V1 | .028 | .259 | .431 | >.999 | >.999 | −.025 | .749 | .990 | >.999 | >.999 | .040 | .367 | −.038 | .639 | — |
| rh | MST | .018 | .293 | .434 | >.999 | >.999 | −.031 | .560 | .990 | >.999 | >.999 | .030 | .348 | −.040 | .459 | — |
| rh | V6 | .012 | .358 | .486 | >.999 | >.999 | .054 | .303 | .990 | >.999 | >.999 | −.006 | .861 | .056 | .302 | — |
| rh | V2 | .045 | .165 | .380 | .990 | >.999 | .026 | .746 | .990 | >.999 | >.999 | .041 | .372 | .013 | .872 | — |
| rh | V3 | .039 | .188 | .398 | .997 | >.999 | .038 | .622 | .990 | >.999 | >.999 | .030 | .497 | .029 | .716 | — |
| rh | V4 | .028 | .255 | .431 | >.999 | >.999 | −.036 | .632 | .990 | >.999 | >.999 | .044 | .306 | −.050 | .521 | — |
| rh | V8 | .054 | .062 | .308 | .962 | >.999 | .019 | .739 | .990 | >.999 | >.999 | .053 | .120 | .003 | .962 | — |
| rh | 4 | .062 | .089 | .315 | .912 | >.999 | .025 | .748 | .990 | >.999 | >.999 | .060 | .181 | .006 | .937 | — |
| rh | 3b | .073 | **.045** | .308 | .791 | >.999 | .030 | .664 | .990 | >.999 | >.999 | .070 | .083 | .008 | .908 | — |
| rh | FEF | .055 | .078 | .309 | .957 | >.999 | .017 | .799 | .990 | >.999 | >.999 | .055 | .143 | −.001 | .993 | — |
| rh | PEF | .010 | .380 | .501 | >.999 | >.999 | .039 | .483 | .990 | >.999 | >.999 | −.002 | .949 | .040 | .484 | — |
| rh | 55b | .056 | .089 | .315 | .950 | >.999 | .018 | .794 | .990 | >.999 | >.999 | .056 | .159 | .001 | .992 | — |
| rh | V3A | −.027 | .776 | .819 | >.999 | >.999 | −.022 | .713 | .990 | >.999 | >.999 | −.022 | .523 | −.015 | .808 | — |
| rh | RSC | .049 | .062 | .308 | .980 | >.999 | .031 | .529 | .990 | >.999 | >.999 | .043 | .159 | .018 | .733 | — |
| rh | POS2 | .017 | .345 | .486 | >.999 | >.999 | −.008 | .911 | .990 | >.999 | >.999 | .021 | .620 | −.015 | .845 | — |
| rh | V7 | −.002 | .517 | .610 | >.999 | >.999 | .055 | .316 | .990 | >.999 | >.999 | −.021 | .520 | .061 | .275 | — |
| rh | IPS1 | .006 | .423 | .546 | >.999 | >.999 | .014 | .804 | .990 | >.999 | >.999 | .002 | .960 | .014 | .816 | — |
| rh | FFC | .072 | .053 | .308 | .804 | >.999 | .126 | .093 | .990 | .997 | .998 | .036 | .401 | .115 | .138 | — |
| rh | V3B | .004 | .445 | .558 | >.999 | >.999 | .078 | .068 | .990 | >.999 | >.999 | −.023 | .388 | .085 | .051 | — |
| rh | LO1 | −.028 | .818 | .851 | >.999 | >.999 | −.019 | .696 | .990 | >.999 | >.999 | −.024 | .423 | −.012 | .818 | — |
| rh | LO2 | .023 | .235 | .423 | >.999 | >.999 | .009 | .870 | .990 | >.999 | >.999 | .022 | .490 | .002 | .970 | — |
| rh | PIT | .022 | .278 | .431 | >.999 | >.999 | .094 | .159 | .990 | >.999 | >.999 | −.008 | .838 | .096 | .160 | — |
| rh | MT | −.002 | .518 | .610 | >.999 | >.999 | −.021 | .692 | .990 | >.999 | >.999 | .005 | .883 | −.022 | .678 | — |
| rh | A1 | −.007 | .579 | .659 | >.999 | >.999 | −.032 | .536 | .990 | >.999 | >.999 | .003 | .914 | −.033 | .533 | — |
| rh | PSL | .059 | .084 | .309 | .931 | >.999 | .027 | .717 | .990 | >.999 | >.999 | .057 | .177 | .009 | .905 | — |
| rh | SFL | .058 | .089 | .315 | .943 | >.999 | .094 | .191 | .990 | >.999 | >.999 | .031 | .455 | .085 | .250 | — |
| rh | PCV | .057 | .070 | .309 | .949 | >.999 | .020 | .753 | .990 | >.999 | >.999 | .056 | .130 | .003 | .968 | — |
| rh | STV | .096 | **.023** | .308 | .444 | >.999 | .047 | .563 | .990 | >.999 | >.999 | .090 | **.047** | .019 | .823 | Matching |
| rh | 7Pm | −.005 | .557 | .641 | >.999 | >.999 | −.025 | .645 | .990 | >.999 | >.999 | .003 | .919 | −.026 | .642 | — |
| rh | 7m | .053 | .073 | .309 | .966 | >.999 | .057 | .360 | .990 | >.999 | >.999 | .039 | .280 | .045 | .489 | — |
| rh | POS1 | .068 | **.041** | .308 | .857 | >.999 | .079 | .221 | .990 | >.999 | >.999 | .047 | .206 | .065 | .334 | — |
| rh | 23d | .017 | .281 | .431 | >.999 | >.999 | −.021 | .669 | .990 | >.999 | >.999 | .026 | .383 | −.029 | .564 | — |
| rh | v23ab | .009 | .377 | .499 | >.999 | >.999 | .013 | .760 | .990 | >.999 | >.999 | .005 | .857 | .012 | .794 | — |
| rh | d23ab | .028 | .214 | .415 | >.999 | >.999 | .044 | .463 | .990 | >.999 | >.999 | .016 | .635 | .038 | .526 | — |
| rh | 31pv | .027 | .211 | .415 | >.999 | >.999 | .037 | .501 | .990 | >.999 | >.999 | .017 | .610 | .032 | .573 | — |
| rh | 5m | .041 | .118 | .347 | .995 | >.999 | .059 | .294 | .990 | >.999 | >.999 | .024 | .466 | .052 | .373 | — |
| rh | 5mv | .006 | .435 | .552 | >.999 | >.999 | −.054 | .391 | .990 | >.999 | >.999 | .025 | .496 | −.061 | .342 | — |
| rh | 23c | −.004 | .537 | .624 | >.999 | >.999 | −.032 | .595 | .990 | >.999 | >.999 | .007 | .840 | −.034 | .577 | — |
| rh | 5L | .003 | .457 | .571 | >.999 | >.999 | −.003 | .953 | .990 | >.999 | >.999 | .004 | .899 | −.005 | .933 | — |
| rh | 24dd | .021 | .279 | .431 | >.999 | >.999 | −.007 | .914 | .990 | >.999 | >.999 | .026 | .473 | −.015 | .818 | — |
| rh | 24dv | .040 | .118 | .347 | .996 | >.999 | .106 | .056 | .990 | >.999 | >.999 | .007 | .825 | .103 | .068 | — |
| rh | 7AL | .025 | .228 | .418 | >.999 | >.999 | −.005 | .934 | .990 | >.999 | >.999 | .029 | .381 | −.014 | .808 | — |
| rh | SCEF | .024 | .268 | .431 | >.999 | >.999 | −.043 | .508 | .990 | >.999 | >.999 | .042 | .271 | −.056 | .398 | — |
| rh | 6ma | .094 | **.017** | .308 | .486 | >.999 | −.047 | .529 | .990 | >.999 | >.999 | .120 | **.005** | −.084 | .261 | Matching |
| rh | 7Am | .011 | .377 | .499 | >.999 | >.999 | .013 | .839 | .990 | >.999 | >.999 | .008 | .829 | .010 | .873 | — |
| rh | 7Pl | .014 | .326 | .472 | >.999 | >.999 | .049 | .326 | .990 | >.999 | >.999 | −.002 | .948 | .049 | .335 | — |
| rh | 7PC | .046 | .111 | .333 | .988 | >.999 | .005 | .930 | .990 | >.999 | >.999 | .049 | .184 | −.010 | .876 | — |
| rh | LIPv | .043 | .085 | .311 | .992 | >.999 | .021 | .692 | .990 | >.999 | >.999 | .041 | .190 | .008 | .882 | — |
| rh | VIP | .012 | .343 | .486 | >.999 | >.999 | .019 | .723 | .990 | >.999 | >.999 | .007 | .813 | .016 | .759 | — |
| rh | MIP | .057 | .084 | .309 | .947 | >.999 | .024 | .742 | .990 | >.999 | >.999 | .055 | .174 | .007 | .929 | — |
| rh | 1 | .044 | .154 | .378 | .992 | >.999 | .034 | .648 | .990 | >.999 | >.999 | .037 | .393 | .022 | .767 | — |
| rh | 2 | .067 | **.048** | .308 | .860 | >.999 | −.023 | .735 | .990 | >.999 | >.999 | .083 | **.036** | −.049 | .488 | Matching |
| rh | 3a | .057 | **.041** | .308 | .946 | >.999 | .023 | .669 | .990 | >.999 | >.999 | .055 | .089 | .006 | .912 | — |
| rh | 6d | .050 | .098 | .320 | .976 | >.999 | −.009 | .888 | .990 | >.999 | >.999 | .058 | .113 | −.027 | .672 | — |
| rh | 6mp | .065 | **.047** | .308 | .885 | >.999 | −.021 | .744 | .990 | >.999 | >.999 | .080 | **.033** | −.046 | .490 | Matching |
| rh | 6v | .045 | .150 | .378 | .990 | >.999 | .006 | .931 | .990 | >.999 | >.999 | .047 | .257 | −.008 | .910 | — |
| rh | p24pr | .016 | .296 | .436 | >.999 | >.999 | .031 | .548 | .990 | >.999 | >.999 | .007 | .814 | .028 | .589 | — |
| rh | 33pr | −.019 | .797 | .834 | >.999 | >.999 | −.027 | .406 | .990 | >.999 | >.999 | −.011 | .621 | −.024 | .487 | — |
| rh | a24pr | .028 | .191 | .399 | >.999 | >.999 | .030 | .572 | .990 | >.999 | >.999 | .020 | .530 | .024 | .664 | — |
| rh | p32pr | −.017 | .675 | .735 | >.999 | >.999 | −.001 | .989 | .990 | >.999 | >.999 | −.018 | .603 | .005 | .938 | — |
| rh | a24 | .001 | .479 | .590 | >.999 | >.999 | −.017 | .762 | .990 | >.999 | >.999 | .007 | .827 | −.020 | .741 | — |
| rh | d32 | .060 | .062 | .308 | .931 | >.999 | −.014 | .820 | .990 | >.999 | >.999 | .071 | .059 | −.037 | .583 | — |
| rh | 8BM | .086 | **.028** | .308 | .599 | >.999 | −.004 | .958 | .990 | >.999 | >.999 | .097 | **.025** | −.034 | .661 | Matching |
| rh | p32 | .021 | .244 | .429 | >.999 | >.999 | −.008 | .876 | .990 | >.999 | >.999 | .026 | .400 | −.016 | .760 | — |
| rh | 10r | .036 | .141 | .373 | .998 | >.999 | .025 | .652 | .990 | >.999 | >.999 | .031 | .344 | .015 | .790 | — |
| rh | 47m | .016 | .301 | .439 | >.999 | >.999 | .058 | .247 | .990 | >.999 | >.999 | −.003 | .931 | .059 | .257 | — |
| rh | 8Av | .131 | **.007** | .249 | .111 | .994 | .095 | .276 | .990 | >.999 | >.999 | .112 | **.021** | .060 | .503 | Matching |
| rh | 8Ad | .105 | **.008** | .249 | .334 | >.999 | .042 | .554 | .990 | >.999 | >.999 | .101 | **.015** | .011 | .883 | Matching |
| rh | 9m | .060 | .107 | .327 | .930 | >.999 | .027 | .733 | .990 | >.999 | >.999 | .057 | .221 | .010 | .907 | — |
| rh | 8BL | .132 | **.006** | .249 | .103 | .992 | .035 | .677 | .990 | >.999 | >.999 | .135 | **.005** | −.007 | .934 | Matching |
| rh | 9p | .078 | **.020** | .308 | .720 | >.999 | .067 | .293 | .990 | >.999 | >.999 | .064 | .084 | .047 | .476 | — |
| rh | 10d | .075 | **.016** | .308 | .762 | >.999 | .048 | .389 | .990 | >.999 | >.999 | .067 | **.046** | .028 | .637 | Matching |
| rh | 8C | .054 | .143 | .373 | .960 | >.999 | .110 | .208 | .990 | >.999 | >.999 | .022 | .655 | .103 | .252 | — |
| rh | 44ᵃ | .148 | **.001** | .225 | **.047** | .930 | .088 | .292 | .990 | >.999 | >.999 | .134 | **.004** | .046 | .591 | Matching |
| rh | 45 | .084 | **.036** | .308 | .630 | >.999 | .035 | .660 | .990 | >.999 | >.999 | .081 | .066 | .010 | .907 | — |
| rh | 47l | .078 | **.020** | .308 | .719 | >.999 | .016 | .800 | .990 | >.999 | >.999 | .081 | **.024** | −.010 | .880 | Matching |
| rh | a47r | .043 | .128 | .364 | .992 | >.999 | −.008 | .898 | .990 | >.999 | >.999 | .051 | .174 | −.024 | .711 | — |
| rh | 6r | .072 | .056 | .308 | .800 | >.999 | .160 | **.043** | .990 | .821 | .823 | .025 | .579 | .152 | .062 | — |
| rh | IFJa | −.007 | .580 | .659 | >.999 | >.999 | −.030 | .568 | .990 | >.999 | >.999 | .003 | .930 | −.031 | .568 | — |
| rh | IFJp | −.067 | .996 | .997 | >.999 | >.999 | −.015 | .698 | .990 | >.999 | >.999 | −.069 | **.006** | .007 | .865 | Matching |
| rh | IFSp | .048 | .135 | .373 | .983 | >.999 | .130 | .080 | .990 | .994 | .994 | .008 | .857 | .128 | .093 | — |
| rh | IFSa | .049 | .144 | .373 | .979 | >.999 | .023 | .783 | .990 | >.999 | >.999 | .046 | .304 | .008 | .923 | — |
| rh | p9-46v | .084 | .052 | .308 | .626 | >.999 | .045 | .607 | .990 | >.999 | >.999 | .078 | .113 | .021 | .819 | — |
| rh | 46 | .085 | **.029** | .308 | .620 | >.999 | .042 | .574 | .990 | >.999 | >.999 | .079 | .061 | .017 | .826 | — |
| rh | a9-46v | .067 | .056 | .308 | .869 | >.999 | .005 | .942 | .990 | >.999 | >.999 | .072 | .073 | −.017 | .807 | — |
| rh | 9-46d | .015 | .356 | .486 | >.999 | >.999 | −.053 | .471 | .990 | >.999 | >.999 | .035 | .410 | −.064 | .399 | — |
| rh | 9a | .091 | **.020** | .308 | .529 | >.999 | .021 | .781 | .990 | >.999 | >.999 | .094 | **.026** | −.009 | .909 | Matching |
| rh | 10v | .066 | **.021** | .308 | .872 | >.999 | .001 | .989 | .990 | >.999 | >.999 | .073 | **.020** | −.022 | .678 | Matching |
| rh | a10p | .035 | .143 | .373 | .998 | >.999 | −.008 | .886 | .990 | >.999 | >.999 | .042 | .191 | −.021 | .704 | — |
| rh | 10pp | .078 | **.008** | .249 | .719 | >.999 | .011 | .835 | .990 | >.999 | >.999 | .083 | **.007** | −.015 | .775 | Matching |
| rh | 11l | .035 | .175 | .386 | .998 | >.999 | −.025 | .699 | .990 | >.999 | >.999 | .048 | .194 | −.040 | .547 | — |
| rh | 13l | .027 | .199 | .406 | >.999 | >.999 | .038 | .479 | .990 | >.999 | >.999 | .017 | .583 | .032 | .552 | — |
| rh | OFC | .031 | .189 | .398 | >.999 | >.999 | −.044 | .456 | .990 | >.999 | >.999 | .050 | .151 | −.060 | .327 | — |
| rh | 47s | .025 | .243 | .429 | >.999 | >.999 | −.005 | .931 | .990 | >.999 | >.999 | .030 | .411 | −.015 | .812 | — |
| rh | LIPd | −.020 | .793 | .833 | >.999 | >.999 | −.087 | **.021** | .990 | >.999 | >.999 | .008 | .756 | −.089 | **.021** | AK− |
| rh | 6a | .030 | .218 | .415 | >.999 | >.999 | .008 | .899 | .990 | >.999 | >.999 | .031 | .421 | −.002 | .981 | — |
| rh | i6-8 | .040 | .159 | .379 | .995 | >.999 | −.003 | .965 | .990 | >.999 | >.999 | .046 | .248 | −.017 | .806 | — |
| rh | s6-8 | .048 | .107 | .327 | .982 | >.999 | .055 | .406 | .990 | >.999 | >.999 | .035 | .367 | .044 | .516 | — |
| rh | 43 | .034 | .173 | .385 | .999 | >.999 | .030 | .631 | .990 | >.999 | >.999 | .028 | .447 | .021 | .738 | — |
| rh | OP4 | .049 | .093 | .319 | .980 | >.999 | .039 | .529 | .990 | >.999 | >.999 | .041 | .255 | .026 | .679 | — |
| rh | OP1 | .014 | .331 | .475 | >.999 | >.999 | .029 | .595 | .990 | >.999 | >.999 | .005 | .880 | .028 | .626 | — |
| rh | OP2-3 | −.046 | .949 | .962 | >.999 | >.999 | −.071 | .112 | .990 | >.999 | >.999 | −.026 | .359 | −.063 | .174 | — |
| rh | 52 | .022 | .198 | .406 | >.999 | >.999 | .015 | .706 | .990 | >.999 | >.999 | .019 | .457 | .009 | .830 | — |
| rh | RI | −.010 | .652 | .716 | >.999 | >.999 | .012 | .749 | .990 | >.999 | >.999 | −.015 | .543 | .016 | .662 | — |
| rh | PFcm | .054 | .059 | .308 | .961 | >.999 | .048 | .400 | .990 | >.999 | >.999 | .043 | .198 | .034 | .558 | — |
| rh | PoI2 | −.040 | .851 | .873 | >.999 | >.999 | −.052 | .419 | .990 | >.999 | >.999 | −.026 | .491 | −.044 | .511 | — |
| rh | TA2 | .014 | .350 | .486 | >.999 | >.999 | −.057 | .341 | .990 | >.999 | >.999 | .035 | .321 | −.068 | .268 | — |
| rh | FOP4 | .062 | **.040** | .308 | .913 | >.999 | .041 | .471 | .990 | >.999 | >.999 | .054 | .113 | .024 | .684 | — |
| rh | MI | .031 | .208 | .414 | >.999 | >.999 | −.003 | .965 | .990 | >.999 | >.999 | .035 | .343 | −.014 | .833 | — |
| rh | Pir | −.001 | .501 | .599 | >.999 | >.999 | −.068 | .244 | .990 | >.999 | >.999 | .023 | .504 | −.075 | .214 | — |
| rh | AVI | .036 | .135 | .373 | .998 | >.999 | −.030 | .576 | .990 | >.999 | >.999 | .050 | .118 | −.046 | .405 | — |
| rh | AAIC | .043 | .110 | .332 | .992 | >.999 | .004 | .951 | .990 | >.999 | >.999 | .047 | .170 | −.011 | .851 | — |
| rh | FOP1 | −.030 | .825 | .856 | >.999 | >.999 | −.065 | .214 | .990 | >.999 | >.999 | −.011 | .728 | −.061 | .254 | — |
| rh | FOP3 | .006 | .395 | .519 | >.999 | >.999 | .005 | .898 | .990 | >.999 | >.999 | .005 | .843 | .004 | .931 | — |
| rh | FOP2 | .014 | .294 | .434 | >.999 | >.999 | .077 | .071 | .990 | >.999 | >.999 | −.011 | .680 | .080 | .066 | — |
| rh | PFt | .030 | .194 | .401 | >.999 | >.999 | −.118 | **.046** | .990 | >.999 | >.999 | .074 | **.033** | −.141 | **.020** | Match + AK− |
| rh | AIP | .036 | .169 | .381 | .998 | >.999 | .094 | .128 | .990 | >.999 | >.999 | .007 | .859 | .092 | .149 | — |
| rh | EC | .020 | .294 | .434 | >.999 | >.999 | −.021 | .767 | .990 | >.999 | >.999 | .030 | .450 | −.030 | .674 | — |
| rh | PreS | .033 | .161 | .379 | >.999 | >.999 | .020 | .705 | .990 | >.999 | >.999 | .029 | .368 | .011 | .842 | — |
| rh | H | −.014 | .662 | .725 | >.999 | >.999 | −.045 | .402 | .990 | >.999 | >.999 | −.000 | .996 | −.045 | .418 | — |
| rh | ProS | .018 | .272 | .431 | >.999 | >.999 | −.002 | .964 | .990 | >.999 | >.999 | .021 | .494 | −.009 | .861 | — |
| rh | PeEc | .039 | .173 | .385 | .996 | >.999 | .062 | .393 | .990 | >.999 | >.999 | .022 | .601 | .055 | .460 | — |
| rh | STGa | −.010 | .602 | .675 | >.999 | >.999 | .003 | .959 | .990 | >.999 | >.999 | −.012 | .728 | .007 | .910 | — |
| rh | PBelt | .057 | **.046** | .308 | .948 | >.999 | −.031 | .580 | .990 | >.999 | >.999 | .074 | **.027** | −.054 | .348 | Matching |
| rh | A5 | .064 | .094 | .319 | .898 | >.999 | −.001 | .986 | .990 | >.999 | >.999 | .071 | .131 | −.024 | .783 | — |
| rh | PHA1 | .036 | .177 | .389 | .998 | >.999 | −.006 | .936 | .990 | >.999 | >.999 | .042 | .279 | −.019 | .784 | — |
| rh | PHA3 | .001 | .487 | .590 | >.999 | >.999 | .034 | .423 | .990 | >.999 | >.999 | −.011 | .678 | .037 | .389 | — |
| rh | STSda | .036 | .194 | .401 | .998 | >.999 | −.018 | .803 | .990 | >.999 | >.999 | .046 | .256 | −.032 | .662 | — |
| rh | STSdp | .103 | **.006** | .249 | .349 | >.999 | .163 | **.020** | .990 | .776 | .778 | .058 | .153 | .145 | **.046** | AK+ |
| rh | STSvp | .123 | **.005** | .249 | .155 | .999 | .066 | .417 | .990 | >.999 | >.999 | .114 | **.013** | .031 | .716 | Matching |
| rh | TGd | .074 | .062 | .308 | .781 | >.999 | .082 | .304 | .990 | >.999 | >.999 | .053 | .242 | .066 | .422 | — |
| rh | TE1a | .068 | .064 | .308 | .857 | >.999 | .086 | .246 | .990 | >.999 | >.999 | .045 | .280 | .072 | .344 | — |
| rh | TE1p | .045 | .163 | .380 | .989 | >.999 | .056 | .494 | .990 | >.999 | >.999 | .031 | .501 | .046 | .580 | — |
| rh | TE2a | .083 | **.032** | .308 | .646 | >.999 | .045 | .554 | .990 | >.999 | >.999 | .076 | .070 | .021 | .792 | — |
| rh | TF | −.002 | .509 | .605 | >.999 | >.999 | −.022 | .760 | .990 | >.999 | >.999 | .005 | .894 | −.024 | .750 | — |
| rh | TE2p | .027 | .237 | .425 | >.999 | >.999 | −.027 | .695 | .990 | >.999 | >.999 | .040 | .304 | −.039 | .576 | — |
| rh | PHT | .089 | **.030** | .308 | .552 | >.999 | .120 | .140 | .990 | >.999 | >.999 | .057 | .213 | .102 | .230 | — |
| rh | PH | .077 | **.021** | .308 | .739 | >.999 | .139 | **.024** | .990 | .977 | .978 | .037 | .300 | .127 | **.044** | AK+ |
| rh | TPOJ1 | .124 | **.009** | .249 | .153 | .999 | .087 | .317 | .990 | >.999 | >.999 | .107 | **.029** | .054 | .552 | Matching |
| rh | TPOJ2 | .009 | .410 | .532 | >.999 | >.999 | −.017 | .831 | .990 | >.999 | >.999 | .016 | .714 | −.022 | .789 | — |
| rh | TPOJ3 | .055 | **.039** | .308 | .958 | >.999 | .114 | **.026** | .990 | >.999 | >.999 | .021 | .496 | .107 | **.043** | AK+ |
| rh | DVT | .021 | .294 | .434 | >.999 | >.999 | .037 | .590 | .990 | >.999 | >.999 | .011 | .788 | .033 | .636 | — |
| rh | PGp | .036 | .183 | .397 | .998 | >.999 | .044 | .512 | .990 | >.999 | >.999 | .024 | .532 | .036 | .599 | — |
| rh | IP2 | .078 | **.020** | .308 | .719 | >.999 | −.034 | .594 | .990 | >.999 | >.999 | .099 | **.008** | −.065 | .324 | Matching |
| rh | IP1 | .042 | .136 | .373 | .994 | >.999 | .055 | .389 | .990 | >.999 | >.999 | .027 | .468 | .046 | .482 | — |
| rh | IP0 | .026 | .232 | .422 | >.999 | >.999 | .139 | **.023** | .990 | .975 | .976 | −.019 | .585 | .145 | **.021** | AK+ |
| rh | PFop | .028 | .218 | .415 | >.999 | >.999 | .003 | .959 | .990 | >.999 | >.999 | .030 | .401 | −.006 | .920 | — |
| rh | PF | .089 | **.018** | .308 | .560 | >.999 | .057 | .431 | .990 | >.999 | >.999 | .079 | .057 | .032 | .664 | — |
| rh | PFm | .080 | **.045** | .308 | .687 | >.999 | .109 | .176 | .990 | >.999 | >.999 | .051 | .262 | .093 | .261 | — |
| rh | PGi | .113 | **.008** | .249 | .240 | >.999 | .141 | .081 | .990 | .968 | .969 | .077 | .091 | .117 | .159 | — |
| rh | PGs | .032 | .225 | .418 | >.999 | >.999 | .070 | .332 | .990 | >.999 | >.999 | .011 | .799 | .067 | .370 | — |
| rh | V6A | .019 | .256 | .431 | >.999 | >.999 | .024 | .605 | .990 | >.999 | >.999 | .013 | .667 | .021 | .679 | — |
| rh | VMV1 | .035 | .167 | .381 | .999 | >.999 | −.066 | .277 | .990 | >.999 | >.999 | .061 | .086 | −.085 | .173 | — |
| rh | VMV3 | .001 | .487 | .590 | >.999 | >.999 | −.046 | .345 | .990 | >.999 | >.999 | .017 | .576 | −.052 | .305 | — |
| rh | PHA2 | −.073 | .997 | .997 | >.999 | >.999 | −.102 | **.017** | .990 | >.999 | >.999 | −.046 | .099 | −.088 | **.050** | AK− |
| rh | V4t | .001 | .482 | .590 | >.999 | >.999 | −.026 | .614 | .990 | >.999 | >.999 | .010 | .753 | −.029 | .583 | — |
| rh | FST | −.024 | .755 | .804 | >.999 | >.999 | −.048 | .411 | .990 | >.999 | >.999 | −.010 | .763 | −.045 | .456 | — |
| rh | V3CD | .011 | .353 | .486 | >.999 | >.999 | .039 | .463 | .990 | >.999 | >.999 | −.001 | .982 | .039 | .477 | — |
| rh | LO3 | −.016 | .667 | .728 | >.999 | >.999 | −.059 | .323 | .990 | >.999 | >.999 | .003 | .934 | −.060 | .330 | — |
| rh | VMV2 | .005 | .430 | .549 | >.999 | >.999 | .045 | .282 | .990 | >.999 | >.999 | −.011 | .694 | .048 | .261 | — |
| rh | 31pd | .043 | .095 | .319 | .993 | >.999 | .029 | .594 | .990 | >.999 | >.999 | .037 | .247 | .018 | .756 | — |
| rh | 31a | .028 | .212 | .415 | >.999 | >.999 | −.002 | .976 | .990 | >.999 | >.999 | .032 | .353 | −.012 | .848 | — |
| rh | VVC | .050 | .103 | .327 | .976 | >.999 | .104 | .119 | .990 | >.999 | >.999 | .019 | .618 | .098 | .153 | — |
| rh | 25 | −.006 | .579 | .659 | >.999 | >.999 | −.005 | .902 | .990 | >.999 | >.999 | −.005 | .858 | −.004 | .936 | — |
| rh | s32 | .026 | .158 | .379 | >.999 | >.999 | −.011 | .788 | .990 | >.999 | >.999 | .033 | .202 | −.021 | .612 | — |
| rh | pOFC | .028 | .207 | .414 | >.999 | >.999 | .020 | .728 | .990 | >.999 | >.999 | .024 | .478 | .013 | .831 | — |
| rh | PoI1 | .019 | .268 | .431 | >.999 | >.999 | .077 | .129 | .990 | >.999 | >.999 | −.005 | .864 | .079 | .132 | — |
| rh | Ig | .019 | .227 | .418 | >.999 | >.999 | .029 | .468 | .990 | >.999 | >.999 | .011 | .671 | .026 | .532 | — |
| rh | FOP5 | −.003 | .533 | .621 | >.999 | >.999 | .017 | .761 | .990 | >.999 | >.999 | −.009 | .782 | .020 | .730 | — |
| rh | p10p | .035 | .163 | .380 | .998 | >.999 | .001 | .989 | .990 | >.999 | >.999 | .038 | .278 | −.011 | .852 | — |
| rh | p47r | .060 | .062 | .308 | .930 | >.999 | .022 | .738 | .990 | >.999 | >.999 | .059 | .120 | .003 | .963 | — |
| rh | TGv | .014 | .351 | .486 | >.999 | >.999 | .008 | .905 | .990 | >.999 | >.999 | .013 | .732 | .004 | .955 | — |
| rh | MBelt | −.016 | .708 | .765 | >.999 | >.999 | −.038 | .436 | .990 | >.999 | >.999 | −.005 | .860 | −.036 | .474 | — |
| rh | LBelt | .055 | **.047** | .308 | .957 | >.999 | .038 | .472 | .990 | >.999 | >.999 | .048 | .136 | .023 | .667 | — |
| rh | A4 | .014 | .367 | .493 | >.999 | >.999 | −.060 | .417 | .990 | >.999 | >.999 | .036 | .390 | −.071 | .347 | — |
| rh | STSva | .037 | .186 | .398 | .998 | >.999 | .027 | .704 | .990 | >.999 | >.999 | .032 | .433 | .017 | .814 | — |
| rh | TE1m | .077 | **.042** | .308 | .741 | >.999 | .067 | .357 | .990 | >.999 | >.999 | .062 | .141 | .048 | .530 | — |
| rh | PI | .021 | .262 | .431 | >.999 | >.999 | −.062 | .272 | .990 | >.999 | >.999 | .045 | .183 | −.076 | .193 | — |
| rh | a32pr | .020 | .283 | .431 | >.999 | >.999 | −.019 | .739 | .990 | >.999 | >.999 | .029 | .411 | −.028 | .637 | — |
| rh | p24 | −.041 | .867 | .887 | >.999 | >.999 | −.004 | .952 | .990 | >.999 | >.999 | −.044 | .220 | .010 | .873 | — |

**Retrospective ELS — Unadjusted (N = 85, 20,000 permutations, 360 parcels, r(|a − b|, mean) = .65)**

|  | | **Matching ρ(D, \|a − b\|)** | | | | | **Additive ρ(D, mean(a, b))** | | | | | **Joint model D ~ \|a − b\| + mean(a, b)** | | | |  |
| --- | --- | --- | --- | --- | --- | --- | --- | --- | --- | --- | --- | --- | --- | --- | --- | --- |
| **Hemi** | **Region** | **ρ** | **p** | **q** | **p FWE-P** | **p FWE-F** | **ρ** | **p** | **q** | **p FWE-P** | **p FWE-F** | **β \|a−b\|** | **p** | **β mean** | **p** | **Mechanism** |
| lh | V1 | −.053 | .826 | .975 | >.999 | >.999 | .006 | .941 | .996 | >.999 | >.999 | −.098 | .105 | .069 | .469 | — |
| lh | MST | .019 | .257 | .975 | >.999 | >.999 | .039 | .316 | .996 | >.999 | >.999 | −.010 | .766 | .046 | .326 | — |
| lh | V6 | −.013 | .621 | .975 | >.999 | >.999 | .006 | .923 | .996 | >.999 | >.999 | −.028 | .542 | .024 | .724 | — |
| lh | V2 | .007 | .443 | .975 | >.999 | >.999 | .051 | .535 | .996 | >.999 | >.999 | −.045 | .476 | .081 | .417 | — |
| lh | V3 | .013 | .406 | .975 | >.999 | >.999 | .075 | .351 | .996 | >.999 | >.999 | −.060 | .326 | .113 | .232 | — |
| lh | V4 | −.066 | .872 | .981 | >.999 | >.999 | .014 | .871 | .996 | >.999 | >.999 | −.129 | **.039** | .097 | .317 | Matching |
| lh | V8 | .026 | .301 | .975 | >.999 | >.999 | .047 | .504 | .996 | >.999 | >.999 | −.006 | .906 | .051 | .542 | — |
| lh | 4 | .011 | .417 | .975 | >.999 | >.999 | .040 | .625 | .996 | >.999 | >.999 | −.026 | .684 | .056 | .560 | — |
| lh | 3b | .007 | .437 | .975 | >.999 | >.999 | −.016 | .824 | .996 | >.999 | >.999 | .029 | .600 | −.034 | .687 | — |
| lh | FEF | .053 | .137 | .975 | .992 | >.999 | .019 | .781 | .996 | >.999 | >.999 | .071 | .179 | −.027 | .739 | — |
| lh | PEF | .003 | .466 | .975 | >.999 | >.999 | −.002 | .971 | .996 | >.999 | >.999 | .008 | .872 | −.007 | .917 | — |
| lh | 55b | −.038 | .751 | .975 | >.999 | >.999 | −.009 | .914 | .996 | >.999 | >.999 | −.056 | .349 | .028 | .768 | — |
| lh | V3A | .031 | .240 | .975 | >.999 | >.999 | .086 | .153 | .996 | >.999 | >.999 | −.042 | .381 | .114 | .115 | — |
| lh | RSC | .006 | .435 | .975 | >.999 | >.999 | −.049 | .392 | .996 | >.999 | >.999 | .065 | .149 | −.091 | .178 | — |
| lh | POS2 | −.046 | .807 | .975 | >.999 | >.999 | −.025 | .737 | .996 | >.999 | >.999 | −.051 | .357 | .008 | .923 | — |
| lh | V7 | −.008 | .592 | .975 | >.999 | >.999 | .003 | .956 | .996 | >.999 | >.999 | −.017 | .660 | .014 | .807 | — |
| lh | IPS1 | −.015 | .624 | .975 | >.999 | >.999 | .012 | .862 | .996 | >.999 | >.999 | −.040 | .448 | .037 | .636 | — |
| lh | FFC | −.017 | .600 | .975 | >.999 | >.999 | −.006 | .943 | .996 | >.999 | >.999 | −.022 | .725 | .009 | .929 | — |
| lh | V3B | .011 | .374 | .975 | >.999 | >.999 | .013 | .766 | .996 | >.999 | >.999 | .004 | .916 | .010 | .843 | — |
| lh | LO1 | .065 | .057 | .975 | .969 | >.999 | .094 | .081 | .996 | >.999 | >.999 | .007 | .877 | .090 | .166 | — |
| lh | LO2 | .006 | .449 | .975 | >.999 | >.999 | .011 | .859 | .996 | >.999 | >.999 | −.003 | .960 | .013 | .858 | — |
| lh | PIT | −.091 | .971 | .987 | >.999 | >.999 | −.020 | .770 | .996 | >.999 | >.999 | −.135 | **.009** | .068 | .402 | Matching |
| lh | MT | .057 | **.049** | .975 | .987 | >.999 | .072 | .103 | .996 | >.999 | >.999 | .017 | .643 | .061 | .258 | — |
| lh | A1 | .011 | .388 | .975 | >.999 | >.999 | .042 | .443 | .996 | >.999 | >.999 | −.028 | .532 | .060 | .356 | — |
| lh | PSL | −.081 | .914 | .984 | >.999 | >.999 | −.077 | .355 | .996 | >.999 | >.999 | −.053 | .400 | −.042 | .671 | — |
| lh | SFL | −.037 | .722 | .975 | >.999 | >.999 | −.026 | .756 | .996 | >.999 | >.999 | −.034 | .596 | −.004 | .967 | — |
| lh | PCV | .006 | .442 | .975 | >.999 | >.999 | −.006 | .920 | .996 | >.999 | >.999 | .018 | .703 | −.018 | .806 | — |
| lh | STV | −.084 | .919 | .984 | >.999 | >.999 | −.092 | .277 | .996 | >.999 | >.999 | −.043 | .514 | −.065 | .520 | — |
| lh | 7Pm | −.004 | .542 | .975 | >.999 | >.999 | −.023 | .715 | .996 | >.999 | >.999 | .018 | .711 | −.034 | .638 | — |
| lh | 7m | .035 | .263 | .975 | >.999 | >.999 | −.033 | .674 | .996 | >.999 | >.999 | .097 | .099 | −.095 | .297 | — |
| lh | POS1 | −.006 | .541 | .975 | >.999 | >.999 | .033 | .623 | .996 | >.999 | >.999 | −.047 | .374 | .063 | .434 | — |
| lh | 23d | −.005 | .543 | .975 | >.999 | >.999 | −.001 | .989 | .996 | >.999 | >.999 | −.008 | .872 | .004 | .952 | — |
| lh | v23ab | −.022 | .714 | .975 | >.999 | >.999 | −.118 | **.024** | .996 | >.999 | >.999 | .094 | **.025** | −.179 | **.004** | Match + AK− |
| lh | d23ab | .033 | .251 | .975 | >.999 | >.999 | .003 | .966 | .996 | >.999 | >.999 | .054 | .310 | −.032 | .693 | — |
| lh | 31pv | −.053 | .905 | .984 | >.999 | >.999 | −.143 | **.009** | .996 | .968 | .971 | .067 | .133 | −.186 | **.004** | AK− |
| lh | 5m | −.007 | .578 | .975 | >.999 | >.999 | .007 | .887 | .996 | >.999 | >.999 | −.021 | .615 | .021 | .733 | — |
| lh | 5mv | −.032 | .795 | .975 | >.999 | >.999 | −.011 | .827 | .996 | >.999 | >.999 | −.042 | .311 | .016 | .793 | — |
| lh | 23c | −.011 | .598 | .975 | >.999 | >.999 | .008 | .887 | .996 | >.999 | >.999 | −.028 | .537 | .026 | .700 | — |
| lh | 5L | −.024 | .713 | .975 | >.999 | >.999 | −.050 | .379 | .996 | >.999 | >.999 | .014 | .764 | −.060 | .383 | — |
| lh | 24dd | −.014 | .611 | .975 | >.999 | >.999 | −.015 | .816 | .996 | >.999 | >.999 | −.007 | .888 | −.011 | .890 | — |
| lh | 24dv | −.054 | .921 | .984 | >.999 | >.999 | −.042 | .415 | .996 | >.999 | >.999 | −.045 | .281 | −.013 | .833 | — |
| lh | 7AL | −.044 | .837 | .979 | >.999 | >.999 | −.025 | .688 | .996 | >.999 | >.999 | −.048 | .324 | .007 | .930 | — |
| lh | SCEF | −.007 | .548 | .975 | >.999 | >.999 | .049 | .538 | .996 | >.999 | >.999 | −.067 | .283 | .092 | .330 | — |
| lh | 6ma | −.039 | .759 | .975 | >.999 | >.999 | −.045 | .554 | .996 | >.999 | >.999 | −.018 | .763 | −.033 | .712 | — |
| lh | 7Am | .010 | .416 | .975 | >.999 | >.999 | .017 | .804 | .996 | >.999 | >.999 | −.001 | .979 | .018 | .828 | — |
| lh | 7Pl | .034 | .217 | .975 | >.999 | >.999 | .038 | .513 | .996 | >.999 | >.999 | .016 | .737 | .028 | .689 | — |
| lh | 7PC | −.004 | .529 | .975 | >.999 | >.999 | .041 | .550 | .996 | >.999 | >.999 | −.053 | .327 | .076 | .359 | — |
| lh | LIPv | −.056 | .882 | .984 | >.999 | >.999 | −.013 | .842 | .996 | >.999 | >.999 | −.082 | .113 | .040 | .606 | — |
| lh | VIP | −.070 | .919 | .984 | >.999 | >.999 | −.051 | .459 | .996 | >.999 | >.999 | −.064 | .241 | −.010 | .907 | — |
| lh | MIP | −.027 | .711 | .975 | >.999 | >.999 | −.007 | .916 | .996 | >.999 | >.999 | −.039 | .442 | .018 | .811 | — |
| lh | 1 | .001 | .481 | .975 | >.999 | >.999 | .029 | .707 | .996 | >.999 | >.999 | −.031 | .603 | .049 | .591 | — |
| lh | 2 | −.088 | .965 | .987 | >.999 | >.999 | −.040 | .560 | .996 | >.999 | >.999 | −.107 | **.046** | .029 | .722 | Matching |
| lh | 3a | .046 | .149 | .975 | .998 | >.999 | .046 | .444 | .996 | >.999 | >.999 | .028 | .565 | .028 | .692 | — |
| lh | 6d | −.007 | .550 | .975 | >.999 | >.999 | .077 | .261 | .996 | >.999 | >.999 | −.098 | .070 | .140 | .085 | — |
| lh | 6mp | −.026 | .727 | .975 | >.999 | >.999 | −.001 | .982 | .996 | >.999 | >.999 | −.044 | .366 | .027 | .707 | — |
| lh | 6v | .038 | .216 | .975 | >.999 | >.999 | .122 | .073 | .996 | >.999 | >.999 | −.069 | .195 | .166 | **.040** | AK+ |
| lh | p24pr | .011 | .395 | .975 | >.999 | >.999 | −.023 | .689 | .996 | >.999 | >.999 | .045 | .331 | −.053 | .453 | — |
| lh | 33pr | .003 | .454 | .975 | >.999 | >.999 | .045 | .310 | .996 | >.999 | >.999 | −.044 | .228 | .073 | .161 | — |
| lh | a24pr | −.042 | .862 | .980 | >.999 | >.999 | −.008 | .875 | .996 | >.999 | >.999 | −.063 | .144 | .032 | .608 | — |
| lh | p32pr | .011 | .392 | .975 | >.999 | >.999 | .009 | .864 | .996 | >.999 | >.999 | .009 | .835 | .003 | .959 | — |
| lh | a24 | .042 | .168 | .975 | >.999 | >.999 | .075 | .198 | .996 | >.999 | >.999 | −.012 | .797 | .083 | .237 | — |
| lh | d32 | .014 | .378 | .975 | >.999 | >.999 | −.046 | .482 | .996 | >.999 | >.999 | .076 | .143 | −.096 | .225 | — |
| lh | 8BM | −.088 | .941 | .984 | >.999 | >.999 | −.158 | **.046** | .996 | .861 | .866 | .025 | .688 | −.174 | .067 | — |
| lh | p32 | −.019 | .717 | .975 | >.999 | >.999 | −.015 | .735 | .996 | >.999 | >.999 | −.016 | .657 | −.004 | .933 | — |
| lh | 10r | −.040 | .841 | .979 | >.999 | >.999 | −.029 | .597 | .996 | >.999 | >.999 | −.037 | .400 | −.005 | .941 | — |
| lh | 47m | −.039 | .848 | .979 | >.999 | >.999 | .002 | .975 | .996 | >.999 | >.999 | −.070 | .092 | .047 | .452 | — |
| lh | 8Av | −.033 | .694 | .975 | >.999 | >.999 | −.054 | .535 | .996 | >.999 | >.999 | .003 | .962 | −.056 | .584 | — |
| lh | 8Ad | .005 | .456 | .975 | >.999 | >.999 | .025 | .706 | .996 | >.999 | >.999 | −.020 | .710 | .038 | .636 | — |
| lh | 9m | .007 | .440 | .975 | >.999 | >.999 | .032 | .705 | .996 | >.999 | >.999 | −.023 | .725 | .047 | .641 | — |
| lh | 8BL | −.004 | .526 | .975 | >.999 | >.999 | −.014 | .844 | .996 | >.999 | >.999 | .009 | .871 | −.020 | .814 | — |
| lh | 9p | −.013 | .598 | .975 | >.999 | >.999 | .033 | .632 | .996 | >.999 | >.999 | −.058 | .274 | .070 | .390 | — |
| lh | 10d | .023 | .295 | .975 | >.999 | >.999 | .044 | .460 | .996 | >.999 | >.999 | −.009 | .839 | .050 | .482 | — |
| lh | 8C | −.056 | .818 | .975 | >.999 | >.999 | −.054 | .535 | .996 | >.999 | >.999 | −.036 | .577 | −.030 | .764 | — |
| lh | 44 | .048 | .220 | .975 | .997 | >.999 | .133 | .122 | .996 | .993 | .994 | −.066 | .317 | .176 | .083 | — |
| lh | 45 | −.020 | .607 | .975 | >.999 | >.999 | −.042 | .643 | .996 | >.999 | >.999 | .013 | .856 | −.050 | .640 | — |
| lh | 47l | −.025 | .666 | .975 | >.999 | >.999 | −.007 | .929 | .996 | >.999 | >.999 | −.035 | .555 | .016 | .864 | — |
| lh | a47r | .012 | .396 | .975 | >.999 | >.999 | .076 | .278 | .996 | >.999 | >.999 | −.063 | .251 | .116 | .161 | — |
| lh | 6r | .021 | .349 | .975 | >.999 | >.999 | .091 | .249 | .996 | >.999 | >.999 | −.065 | .290 | .132 | .158 | — |
| lh | IFJa | −.010 | .582 | .975 | >.999 | >.999 | −.003 | .966 | .996 | >.999 | >.999 | −.015 | .780 | .007 | .936 | — |
| lh | IFJp | −.038 | .825 | .975 | >.999 | >.999 | −.012 | .831 | .996 | >.999 | >.999 | −.053 | .242 | .022 | .744 | — |
| lh | IFSp | .019 | .360 | .975 | >.999 | >.999 | .037 | .627 | .996 | >.999 | >.999 | −.007 | .905 | .042 | .642 | — |
| lh | IFSaᵃ | −.092 | .928 | .984 | >.999 | >.999 | −.053 | .553 | .996 | >.999 | >.999 | −.099 | .147 | .011 | .916 | — |
| lh | p9-46v | −.026 | .656 | .975 | >.999 | >.999 | −.084 | .331 | .996 | >.999 | >.999 | .048 | .460 | −.115 | .257 | — |
| lh | 46 | .014 | .384 | .975 | >.999 | >.999 | .067 | .373 | .996 | >.999 | >.999 | −.050 | .396 | .099 | .273 | — |
| lh | a9-46v | .015 | .386 | .975 | >.999 | >.999 | .019 | .797 | .996 | >.999 | >.999 | .005 | .936 | .016 | .860 | — |
| lh | 9-46d | −.013 | .578 | .975 | >.999 | >.999 | −.015 | .849 | .996 | >.999 | >.999 | −.005 | .935 | −.012 | .899 | — |
| lh | 9a | −.033 | .702 | .975 | >.999 | >.999 | −.019 | .823 | .996 | >.999 | >.999 | −.036 | .572 | .004 | .963 | — |
| lh | 10v | −.013 | .612 | .975 | >.999 | >.999 | .052 | .366 | .996 | >.999 | >.999 | −.080 | .081 | .104 | .130 | — |
| lh | a10p | −.023 | .715 | .975 | >.999 | >.999 | .004 | .943 | .996 | >.999 | >.999 | −.045 | .307 | .033 | .620 | — |
| lh | 10pp | −.045 | .889 | .984 | >.999 | >.999 | −.007 | .890 | .996 | >.999 | >.999 | −.070 | .087 | .038 | .523 | — |
| lh | 11l | −.039 | .796 | .975 | >.999 | >.999 | .019 | .773 | .996 | >.999 | >.999 | −.088 | .079 | .075 | .321 | — |
| lh | 13l | .009 | .416 | .975 | >.999 | >.999 | .090 | .153 | .996 | >.999 | >.999 | −.084 | .088 | .145 | .051 | — |
| lh | OFC | .036 | .203 | .975 | >.999 | >.999 | .062 | .298 | .996 | >.999 | >.999 | −.006 | .893 | .066 | .350 | — |
| lh | 47s | −.010 | .585 | .975 | >.999 | >.999 | .058 | .327 | .996 | >.999 | >.999 | −.081 | .083 | .110 | .118 | — |
| lh | LIPd | .012 | .365 | .975 | >.999 | >.999 | −.051 | .300 | .996 | >.999 | >.999 | .078 | .053 | −.102 | .086 | — |
| lh | 6a | −.023 | .680 | .975 | >.999 | >.999 | −.005 | .941 | .996 | >.999 | >.999 | −.034 | .498 | .017 | .823 | — |
| lh | i6-8 | .013 | .382 | .975 | >.999 | >.999 | .028 | .650 | .996 | >.999 | >.999 | −.008 | .867 | .033 | .649 | — |
| lh | s6-8 | −.025 | .726 | .975 | >.999 | >.999 | −.037 | .507 | .996 | >.999 | >.999 | −.001 | .978 | −.036 | .588 | — |
| lh | 43 | −.016 | .651 | .975 | >.999 | >.999 | .013 | .809 | .996 | >.999 | >.999 | −.043 | .329 | .041 | .529 | — |
| lh | OP4 | −.043 | .814 | .975 | >.999 | >.999 | .034 | .612 | .996 | >.999 | >.999 | −.112 | **.033** | .106 | .180 | Matching |
| lh | OP1 | −.026 | .729 | .975 | >.999 | >.999 | .022 | .707 | .996 | >.999 | >.999 | −.070 | .130 | .068 | .336 | — |
| lh | OP2-3 | −.039 | .886 | .984 | >.999 | >.999 | −.020 | .649 | .996 | >.999 | >.999 | −.046 | .211 | .010 | .851 | — |
| lh | 52 | .039 | .137 | .975 | >.999 | >.999 | .089 | .058 | .996 | >.999 | >.999 | −.032 | .418 | .110 | .054 | — |
| lh | RI | .027 | .238 | .975 | >.999 | >.999 | .062 | .221 | .996 | >.999 | >.999 | −.022 | .596 | .076 | .208 | — |
| lh | PFcm | .004 | .465 | .975 | >.999 | >.999 | .034 | .593 | .996 | >.999 | >.999 | −.031 | .534 | .054 | .475 | — |
| lh | PoI2 | .012 | .400 | .975 | >.999 | >.999 | .031 | .618 | .996 | >.999 | >.999 | −.014 | .776 | .040 | .588 | — |
| lh | TA2 | −.031 | .772 | .975 | >.999 | >.999 | −.002 | .969 | .996 | >.999 | >.999 | −.050 | .256 | .030 | .646 | — |
| lh | FOP4 | .024 | .304 | .975 | >.999 | >.999 | .052 | .421 | .996 | >.999 | >.999 | −.017 | .740 | .063 | .413 | — |
| lh | MI | .041 | .193 | .975 | >.999 | >.999 | .083 | .208 | .996 | >.999 | >.999 | −.021 | .678 | .096 | .217 | — |
| lh | Pir | .030 | .225 | .975 | >.999 | >.999 | .085 | .115 | .996 | >.999 | >.999 | −.042 | .334 | .112 | .083 | — |
| lh | AVI | .012 | .400 | .975 | >.999 | >.999 | .018 | .789 | .996 | >.999 | >.999 | .000 | .996 | .018 | .816 | — |
| lh | AAIC | .018 | .328 | .975 | >.999 | >.999 | .025 | .664 | .996 | >.999 | >.999 | .004 | .929 | .022 | .743 | — |
| lh | FOP1 | .002 | .477 | .975 | >.999 | >.999 | .056 | .313 | .996 | >.999 | >.999 | −.059 | .182 | .095 | .154 | — |
| lh | FOP3 | .029 | .205 | .975 | >.999 | >.999 | .032 | .508 | .996 | >.999 | >.999 | .014 | .723 | .023 | .688 | — |
| lh | FOP2 | .033 | .159 | .975 | >.999 | >.999 | .069 | .111 | .996 | >.999 | >.999 | −.019 | .603 | .081 | .116 | — |
| lh | PFt | −.020 | .686 | .975 | >.999 | >.999 | .040 | .461 | .996 | >.999 | >.999 | −.079 | .067 | .091 | .158 | — |
| lh | AIP | −.119 | .998 | .998 | >.999 | >.999 | −.101 | .089 | .996 | >.999 | >.999 | −.093 | **.044** | −.041 | .564 | Matching |
| lh | EC | .010 | .417 | .975 | >.999 | >.999 | .068 | .281 | .996 | >.999 | >.999 | −.059 | .238 | .106 | .156 | — |
| lh | PreS | −.021 | .675 | .975 | >.999 | >.999 | −.005 | .942 | .996 | >.999 | >.999 | −.031 | .518 | .016 | .831 | — |
| lh | H | −.058 | .911 | .984 | >.999 | >.999 | −.057 | .331 | .996 | >.999 | >.999 | −.036 | .427 | −.033 | .636 | — |
| lh | ProS | −.008 | .590 | .975 | >.999 | >.999 | .012 | .794 | .996 | >.999 | >.999 | −.027 | .474 | .029 | .585 | — |
| lh | PeEc | −.083 | .937 | .984 | >.999 | >.999 | −.033 | .665 | .996 | >.999 | >.999 | −.106 | .071 | .036 | .697 | — |
| lh | STGa | −.022 | .673 | .975 | >.999 | >.999 | −.006 | .936 | .996 | >.999 | >.999 | −.031 | .557 | .014 | .860 | — |
| lh | PBelt | .043 | .153 | .975 | .999 | >.999 | .063 | .269 | .996 | >.999 | >.999 | .004 | .929 | .060 | .376 | — |
| lh | A5 | −.057 | .819 | .975 | >.999 | >.999 | −.034 | .709 | .996 | >.999 | >.999 | −.061 | .360 | .006 | .951 | — |
| lh | PHA1 | .051 | .123 | .975 | .995 | >.999 | .077 | .190 | .996 | >.999 | >.999 | .001 | .986 | .077 | .280 | — |
| lh | PHA3 | −.036 | .775 | .975 | >.999 | >.999 | −.003 | .962 | .996 | >.999 | >.999 | −.058 | .248 | .034 | .656 | — |
| lh | STSda | −.017 | .628 | .975 | >.999 | >.999 | −.033 | .630 | .996 | >.999 | >.999 | .008 | .886 | −.038 | .636 | — |
| lh | STSdp | −.045 | .807 | .975 | >.999 | >.999 | −.006 | .933 | .996 | >.999 | >.999 | −.071 | .208 | .040 | .645 | — |
| lh | STSvp | −.133 | .990 | .995 | >.999 | >.999 | −.115 | .165 | .996 | >.999 | >.999 | −.101 | .112 | −.050 | .616 | — |
| lh | TGd | −.018 | .611 | .975 | >.999 | >.999 | .046 | .584 | .996 | >.999 | >.999 | −.083 | .200 | .100 | .318 | — |
| lh | TE1a | −.026 | .667 | .975 | >.999 | >.999 | .033 | .679 | .996 | >.999 | >.999 | −.082 | .191 | .086 | .377 | — |
| lh | TE1p | −.008 | .540 | .975 | >.999 | >.999 | −.033 | .699 | .996 | >.999 | >.999 | .024 | .716 | −.049 | .635 | — |
| lh | TE2a | −.018 | .617 | .975 | >.999 | >.999 | .051 | .514 | .996 | >.999 | >.999 | −.089 | .141 | .109 | .244 | — |
| lh | TF | −.045 | .794 | .975 | >.999 | >.999 | −.045 | .554 | .996 | >.999 | >.999 | −.028 | .638 | −.027 | .765 | — |
| lh | TE2p | −.025 | .663 | .975 | >.999 | >.999 | −.038 | .623 | .996 | >.999 | >.999 | .000 | .994 | −.039 | .677 | — |
| lh | PHT | −.087 | .914 | .984 | >.999 | >.999 | −.099 | .277 | .996 | >.999 | >.999 | −.040 | .572 | −.073 | .495 | — |
| lh | PH | .005 | .457 | .975 | >.999 | >.999 | .045 | .551 | .996 | >.999 | >.999 | −.041 | .483 | .071 | .420 | — |
| lh | TPOJ1 | −.020 | .630 | .975 | >.999 | >.999 | −.023 | .773 | .996 | >.999 | >.999 | −.009 | .884 | −.017 | .849 | — |
| lh | TPOJ2 | −.033 | .710 | .975 | >.999 | >.999 | −.024 | .761 | .996 | >.999 | >.999 | −.030 | .634 | −.005 | .954 | — |
| lh | TPOJ3 | .016 | .318 | .975 | >.999 | >.999 | .077 | .108 | .996 | >.999 | >.999 | −.057 | .145 | .114 | **.044** | AK+ |
| lh | DVT | −.034 | .760 | .975 | >.999 | >.999 | −.034 | .607 | .996 | >.999 | >.999 | −.021 | .674 | −.020 | .797 | — |
| lh | PGp | .012 | .396 | .975 | >.999 | >.999 | .047 | .499 | .996 | >.999 | >.999 | −.032 | .562 | .068 | .418 | — |
| lh | IP2 | .004 | .454 | .975 | >.999 | >.999 | −.046 | .408 | .996 | >.999 | >.999 | .059 | .189 | −.084 | .207 | — |
| lh | IP1 | .016 | .358 | .975 | >.999 | >.999 | .012 | .842 | .996 | >.999 | >.999 | .015 | .763 | .003 | .973 | — |
| lh | IP0 | −.067 | .941 | .984 | >.999 | >.999 | −.069 | .242 | .996 | >.999 | >.999 | −.038 | .422 | −.044 | .529 | — |
| lh | PFop | −.041 | .824 | .975 | >.999 | >.999 | .004 | .944 | .996 | >.999 | >.999 | −.075 | .108 | .053 | .455 | — |
| lh | PF | −.055 | .818 | .975 | >.999 | >.999 | −.029 | .725 | .996 | >.999 | >.999 | −.062 | .326 | .011 | .909 | — |
| lh | PFm | −.072 | .894 | .984 | >.999 | >.999 | −.072 | .380 | .996 | >.999 | >.999 | −.044 | .484 | −.044 | .653 | — |
| lh | PGi | −.115 | .972 | .987 | >.999 | >.999 | −.110 | .200 | .996 | >.999 | >.999 | −.075 | .249 | −.062 | .549 | — |
| lh | PGsᵃ | −.075 | .932 | .984 | >.999 | >.999 | −.063 | .370 | .996 | >.999 | >.999 | −.059 | .282 | −.025 | .762 | — |
| lh | V6A | −.022 | .738 | .975 | >.999 | >.999 | .028 | .549 | .996 | >.999 | >.999 | −.069 | .071 | .073 | .189 | — |
| lh | VMV1 | .004 | .462 | .975 | >.999 | >.999 | .045 | .362 | .996 | >.999 | >.999 | −.045 | .280 | .074 | .213 | — |
| lh | VMV3 | −.029 | .778 | .975 | >.999 | >.999 | −.028 | .584 | .996 | >.999 | >.999 | −.019 | .661 | −.016 | .791 | — |
| lh | PHA2 | .030 | .152 | .975 | >.999 | >.999 | .065 | .094 | .996 | >.999 | >.999 | −.020 | .563 | .077 | .099 | — |
| lh | V4t | −.026 | .750 | .975 | >.999 | >.999 | .021 | .680 | .996 | >.999 | >.999 | −.067 | .102 | .064 | .283 | — |
| lh | FST | .003 | .472 | .975 | >.999 | >.999 | .103 | .129 | .996 | >.999 | >.999 | −.109 | **.038** | .174 | **.032** | Match + AK+ |
| lh | V3CD | −.010 | .589 | .975 | >.999 | >.999 | .014 | .807 | .996 | >.999 | >.999 | −.033 | .485 | .036 | .621 | — |
| lh | LO3 | .052 | .090 | .975 | .994 | >.999 | .083 | .104 | .996 | >.999 | >.999 | −.004 | .923 | .086 | .160 | — |
| lh | VMV2 | −.040 | .894 | .984 | >.999 | >.999 | −.034 | .414 | .996 | >.999 | >.999 | −.030 | .405 | −.015 | .762 | — |
| lh | 31pd | .020 | .281 | .975 | >.999 | >.999 | .002 | .958 | .996 | >.999 | >.999 | .032 | .410 | −.018 | .751 | — |
| lh | 31a | −.040 | .799 | .975 | >.999 | >.999 | −.129 | **.043** | .996 | .998 | .998 | .075 | .135 | −.177 | **.020** | AK− |
| lh | VVC | −.009 | .562 | .975 | >.999 | >.999 | .036 | .614 | .996 | >.999 | >.999 | −.056 | .323 | .072 | .403 | — |
| lh | 25 | .045 | .114 | .975 | .998 | >.999 | .013 | .799 | .996 | >.999 | >.999 | .064 | .116 | −.029 | .631 | — |
| lh | s32 | −.040 | .960 | .987 | >.999 | >.999 | −.049 | .090 | .996 | >.999 | >.999 | −.015 | .590 | −.039 | .275 | — |
| lh | pOFC | −.031 | .748 | .975 | >.999 | >.999 | −.015 | .818 | .996 | >.999 | >.999 | −.038 | .447 | .010 | .897 | — |
| lh | PoI1 | −.017 | .658 | .975 | >.999 | >.999 | .002 | .978 | .996 | >.999 | >.999 | −.031 | .503 | .022 | .752 | — |
| lh | Ig | −.013 | .641 | .975 | >.999 | >.999 | .018 | .710 | .996 | >.999 | >.999 | −.043 | .286 | .046 | .433 | — |
| lh | FOP5 | .014 | .386 | .975 | >.999 | >.999 | −.008 | .901 | .996 | >.999 | >.999 | .033 | .533 | −.030 | .712 | — |
| lh | p10p | −.021 | .695 | .975 | >.999 | >.999 | −.028 | .607 | .996 | >.999 | >.999 | −.005 | .911 | −.025 | .706 | — |
| lh | p47r | −.058 | .860 | .980 | >.999 | >.999 | −.064 | .390 | .996 | >.999 | >.999 | −.027 | .636 | −.047 | .599 | — |
| lh | TGv | −.036 | .746 | .975 | >.999 | >.999 | .004 | .953 | .996 | >.999 | >.999 | −.067 | .243 | .048 | .582 | — |
| lh | MBelt | .031 | .223 | .975 | >.999 | >.999 | .002 | .966 | .996 | >.999 | >.999 | .051 | .248 | −.031 | .635 | — |
| lh | LBelt | −.053 | .946 | .984 | >.999 | >.999 | −.074 | .098 | .996 | >.999 | >.999 | −.009 | .799 | −.068 | .204 | — |
| lh | A4 | .037 | .271 | .975 | >.999 | >.999 | .092 | .285 | .996 | >.999 | >.999 | −.039 | .551 | .117 | .251 | — |
| lh | STSva | −.085 | .977 | .987 | >.999 | >.999 | −.001 | .990 | .996 | >.999 | >.999 | −.146 | **.002** | .094 | .181 | Matching |
| lh | TE1m | −.035 | .741 | .975 | >.999 | >.999 | −.003 | .966 | .996 | >.999 | >.999 | −.056 | .316 | .033 | .698 | — |
| lh | PI | .028 | .261 | .975 | >.999 | >.999 | .035 | .554 | .996 | >.999 | >.999 | .009 | .846 | .029 | .677 | — |
| lh | a32pr | .039 | .213 | .975 | >.999 | >.999 | .052 | .443 | .996 | >.999 | >.999 | .009 | .862 | .046 | .563 | — |
| lh | p24 | .023 | .312 | .975 | >.999 | >.999 | −.007 | .908 | .996 | >.999 | >.999 | .047 | .345 | −.038 | .624 | — |
| rh | V1 | −.010 | .563 | .975 | >.999 | >.999 | .037 | .640 | .996 | >.999 | >.999 | −.059 | .330 | .075 | .422 | — |
| rh | MST | .002 | .477 | .975 | >.999 | >.999 | .007 | .898 | .996 | >.999 | >.999 | −.004 | .933 | .009 | .886 | — |
| rh | V6 | .007 | .441 | .975 | >.999 | >.999 | .053 | .335 | .996 | >.999 | >.999 | −.047 | .273 | .083 | .196 | — |
| rh | V2 | −.007 | .544 | .975 | >.999 | >.999 | .035 | .671 | .996 | >.999 | >.999 | −.051 | .410 | .068 | .484 | — |
| rh | V3 | .015 | .392 | .975 | >.999 | >.999 | .060 | .439 | .996 | >.999 | >.999 | −.042 | .490 | .087 | .346 | — |
| rh | V4 | .012 | .407 | .975 | >.999 | >.999 | .050 | .509 | .996 | >.999 | >.999 | −.035 | .551 | .073 | .422 | — |
| rh | V8 | −.091 | .985 | .993 | >.999 | >.999 | −.081 | .178 | .996 | >.999 | >.999 | −.067 | .149 | −.038 | .600 | — |
| rh | 4 | .039 | .245 | .975 | >.999 | >.999 | .053 | .500 | .996 | >.999 | >.999 | .008 | .891 | .048 | .611 | — |
| rh | 3b | .015 | .385 | .975 | >.999 | >.999 | .015 | .838 | .996 | >.999 | >.999 | .009 | .877 | .009 | .918 | — |
| rh | FEF | −.038 | .786 | .975 | >.999 | >.999 | .016 | .810 | .996 | >.999 | >.999 | −.083 | .110 | .070 | .382 | — |
| rh | PEF | −.057 | .909 | .984 | >.999 | >.999 | .023 | .701 | .996 | >.999 | >.999 | −.123 | **.007** | .102 | .141 | Matching |
| rh | 55b | −.038 | .764 | .975 | >.999 | >.999 | .004 | .960 | .996 | >.999 | >.999 | −.069 | .211 | .048 | .564 | — |
| rh | V3A | −.016 | .642 | .975 | >.999 | >.999 | −.011 | .861 | .996 | >.999 | >.999 | −.016 | .735 | −.000 | .996 | — |
| rh | RSC | −.018 | .679 | .975 | >.999 | >.999 | −.081 | .099 | .996 | >.999 | >.999 | .059 | .151 | −.120 | **.044** | AK− |
| rh | POS2 | −.006 | .530 | .975 | >.999 | >.999 | −.013 | .868 | .996 | >.999 | >.999 | .004 | .937 | −.016 | .862 | — |
| rh | V7 | −.008 | .572 | .975 | >.999 | >.999 | .023 | .683 | .996 | >.999 | >.999 | −.039 | .388 | .048 | .475 | — |
| rh | IPS1 | .006 | .448 | .975 | >.999 | >.999 | .008 | .887 | .996 | >.999 | >.999 | .001 | .987 | .007 | .912 | — |
| rh | FFC | −.006 | .532 | .975 | >.999 | >.999 | .029 | .704 | .996 | >.999 | >.999 | −.043 | .467 | .057 | .535 | — |
| rh | V3B | −.050 | .940 | .984 | >.999 | >.999 | −.007 | .880 | .996 | >.999 | >.999 | −.079 | **.030** | .045 | .391 | Matching |
| rh | LO1 | .073 | **.028** | .975 | .936 | >.999 | .056 | .275 | .996 | >.999 | >.999 | .063 | .125 | .015 | .809 | — |
| rh | LO2 | −.016 | .660 | .975 | >.999 | >.999 | −.024 | .648 | .996 | >.999 | >.999 | −.001 | .983 | −.024 | .711 | — |
| rh | PIT | .022 | .326 | .975 | >.999 | >.999 | .044 | .511 | .996 | >.999 | >.999 | −.011 | .832 | .051 | .518 | — |
| rh | MT | −.012 | .612 | .975 | >.999 | >.999 | .026 | .625 | .996 | >.999 | >.999 | −.049 | .253 | .057 | .360 | — |
| rh | A1 | .063 | .059 | .975 | .975 | >.999 | .012 | .818 | .996 | >.999 | >.999 | .094 | **.026** | −.049 | .435 | Matching |
| rh | PSL | −.047 | .800 | .975 | >.999 | >.999 | −.007 | .921 | .996 | >.999 | >.999 | −.072 | .222 | .039 | .663 | — |
| rh | SFL | .007 | .442 | .975 | >.999 | >.999 | .039 | .599 | .996 | >.999 | >.999 | −.032 | .574 | .059 | .500 | — |
| rh | PCV | −.095 | .975 | .987 | >.999 | >.999 | −.149 | **.023** | .996 | .935 | .940 | .003 | .954 | −.151 | .054 | — |
| rh | STV | −.030 | .683 | .975 | >.999 | >.999 | .014 | .865 | .996 | >.999 | >.999 | −.067 | .293 | .058 | .553 | — |
| rh | 7Pm | −.024 | .724 | .975 | >.999 | >.999 | −.038 | .499 | .996 | >.999 | >.999 | .000 | .997 | −.038 | .568 | — |
| rh | 7m | −.071 | .942 | .984 | >.999 | >.999 | −.107 | .091 | .996 | >.999 | >.999 | −.003 | .962 | −.105 | .165 | — |
| rh | POS1 | −.027 | .721 | .975 | >.999 | >.999 | −.036 | .582 | .996 | >.999 | >.999 | −.007 | .882 | −.031 | .691 | — |
| rh | 23d | .012 | .354 | .975 | >.999 | >.999 | .039 | .421 | .996 | >.999 | >.999 | −.023 | .574 | .054 | .359 | — |
| rh | v23ab | .029 | .191 | .975 | >.999 | >.999 | .035 | .416 | .996 | >.999 | >.999 | .010 | .781 | .028 | .584 | — |
| rh | d23ab | .007 | .435 | .975 | >.999 | >.999 | −.027 | .657 | .996 | >.999 | >.999 | .041 | .389 | −.053 | .463 | — |
| rh | 31pv | −.014 | .635 | .975 | >.999 | >.999 | −.002 | .974 | .996 | >.999 | >.999 | −.022 | .625 | .012 | .854 | — |
| rh | 5m | −.017 | .656 | .975 | >.999 | >.999 | −.038 | .508 | .996 | >.999 | >.999 | .014 | .763 | −.047 | .492 | — |
| rh | 5mv | −.008 | .561 | .975 | >.999 | >.999 | .041 | .515 | .996 | >.999 | >.999 | −.059 | .229 | .079 | .288 | — |
| rh | 23c | −.010 | .579 | .975 | >.999 | >.999 | −.016 | .784 | .996 | >.999 | >.999 | .001 | .990 | −.016 | .818 | — |
| rh | 5L | −.004 | .532 | .975 | >.999 | >.999 | −.038 | .498 | .996 | >.999 | >.999 | .035 | .431 | −.061 | .362 | — |
| rh | 24dd | .034 | .228 | .975 | >.999 | >.999 | .041 | .506 | .996 | >.999 | >.999 | .013 | .790 | .032 | .651 | — |
| rh | 24dv | .017 | .342 | .975 | >.999 | >.999 | .054 | .337 | .996 | >.999 | >.999 | −.031 | .503 | .074 | .270 | — |
| rh | 7AL | −.007 | .560 | .975 | >.999 | >.999 | −.019 | .731 | .996 | >.999 | >.999 | .010 | .835 | −.026 | .706 | — |
| rh | SCEF | −.009 | .572 | .975 | >.999 | >.999 | .020 | .760 | .996 | >.999 | >.999 | −.038 | .454 | .044 | .561 | — |
| rh | 6ma | −.032 | .716 | .975 | >.999 | >.999 | .035 | .648 | .996 | >.999 | >.999 | −.093 | .109 | .095 | .292 | — |
| rh | 7Am | −.012 | .599 | .975 | >.999 | >.999 | −.028 | .673 | .996 | >.999 | >.999 | .009 | .855 | −.034 | .664 | — |
| rh | 7Pl | .008 | .419 | .975 | >.999 | >.999 | −.000 | >.999 | >.999 | >.999 | >.999 | .014 | .745 | −.009 | .885 | — |
| rh | 7PC | −.075 | .951 | .987 | >.999 | >.999 | −.063 | .318 | .996 | >.999 | >.999 | −.058 | .253 | −.026 | .731 | — |
| rh | LIPv | −.029 | .763 | .975 | >.999 | >.999 | .002 | .967 | .996 | >.999 | >.999 | −.053 | .219 | .036 | .574 | — |
| rh | VIP | −.032 | .784 | .975 | >.999 | >.999 | −.037 | .490 | .996 | >.999 | >.999 | −.014 | .754 | −.028 | .665 | — |
| rh | MIP | −.005 | .534 | .975 | >.999 | >.999 | .003 | .970 | .996 | >.999 | >.999 | −.012 | .837 | .011 | .899 | — |
| rh | 1 | .045 | .204 | .975 | .998 | >.999 | .073 | .336 | .996 | >.999 | >.999 | −.004 | .950 | .075 | .402 | — |
| rh | 2 | −.097 | .973 | .987 | >.999 | >.999 | −.122 | .081 | .996 | >.999 | >.999 | −.031 | .563 | −.101 | .220 | — |
| rh | 3a | −.036 | .815 | .975 | >.999 | >.999 | −.011 | .846 | .996 | >.999 | >.999 | −.049 | .257 | .021 | .734 | — |
| rh | 6d | .033 | .240 | .975 | >.999 | >.999 | .057 | .375 | .996 | >.999 | >.999 | −.007 | .891 | .062 | .422 | — |
| rh | 6mp | −.001 | .500 | .975 | >.999 | >.999 | .023 | .727 | .996 | >.999 | >.999 | −.027 | .586 | .041 | .607 | — |
| rh | 6v | .008 | .434 | .975 | >.999 | >.999 | .049 | .508 | .996 | >.999 | >.999 | −.040 | .481 | .075 | .397 | — |
| rh | p24pr | .053 | .086 | .975 | .993 | >.999 | .073 | .156 | .996 | >.999 | >.999 | .009 | .829 | .067 | .279 | — |
| rh | 33pr | .003 | .453 | .975 | >.999 | >.999 | .003 | .926 | .996 | >.999 | >.999 | .002 | .949 | .002 | .967 | — |
| rh | a24pr | −.040 | .844 | .979 | >.999 | >.999 | −.071 | .183 | .996 | >.999 | >.999 | .009 | .831 | −.077 | .228 | — |
| rh | p32pr | −.003 | .525 | .975 | >.999 | >.999 | .021 | .725 | .996 | >.999 | >.999 | −.030 | .532 | .041 | .569 | — |
| rh | a24 | .073 | **.050** | .975 | .938 | >.999 | .081 | .172 | .996 | >.999 | >.999 | .035 | .459 | .058 | .409 | — |
| rh | d32 | .053 | .144 | .975 | .993 | >.999 | −.004 | .954 | .996 | >.999 | >.999 | .095 | .071 | −.065 | .422 | — |
| rh | 8BM | −.010 | .569 | .975 | >.999 | >.999 | −.012 | .880 | .996 | >.999 | >.999 | −.004 | .950 | −.010 | .918 | — |
| rh | p32 | −.031 | .796 | .975 | >.999 | >.999 | −.051 | .322 | .996 | >.999 | >.999 | .004 | .931 | −.054 | .386 | — |
| rh | 10r | −.013 | .627 | .975 | >.999 | >.999 | .021 | .706 | .996 | >.999 | >.999 | −.046 | .302 | .051 | .445 | — |
| rh | 47m | .008 | .406 | .975 | >.999 | >.999 | .038 | .462 | .996 | >.999 | >.999 | −.028 | .508 | .056 | .368 | — |
| rh | 8Av | −.028 | .655 | .975 | >.999 | >.999 | −.057 | .529 | .996 | >.999 | >.999 | .015 | .830 | −.066 | .535 | — |
| rh | 8Ad | .033 | .266 | .975 | >.999 | >.999 | .065 | .370 | .996 | >.999 | >.999 | −.017 | .773 | .076 | .381 | — |
| rh | 9m | −.034 | .706 | .975 | >.999 | >.999 | .011 | .897 | .996 | >.999 | >.999 | −.071 | .276 | .056 | .570 | — |
| rh | 8BL | −.026 | .650 | .975 | >.999 | >.999 | .006 | .945 | .996 | >.999 | >.999 | −.052 | .437 | .040 | .704 | — |
| rh | 9p | .009 | .413 | .975 | >.999 | >.999 | .006 | .925 | .996 | >.999 | >.999 | .009 | .851 | .000 | >.999 | — |
| rh | 10d | −.002 | .514 | .975 | >.999 | >.999 | .009 | .874 | .996 | >.999 | >.999 | −.014 | .765 | .018 | .794 | — |
| rh | 8C | −.009 | .540 | .975 | >.999 | >.999 | −.058 | .522 | .996 | >.999 | >.999 | .048 | .482 | −.089 | .407 | — |
| rh | 44ᵃ | .009 | .428 | .975 | >.999 | >.999 | .038 | .656 | .996 | >.999 | >.999 | −.026 | .692 | .055 | .586 | — |
| rh | 45 | −.020 | .625 | .975 | >.999 | >.999 | .040 | .622 | .996 | >.999 | >.999 | −.079 | .200 | .091 | .340 | — |
| rh | 47l | .019 | .341 | .975 | >.999 | >.999 | .095 | .144 | .996 | >.999 | >.999 | −.073 | .152 | .142 | .065 | — |
| rh | a47r | .002 | .481 | .975 | >.999 | >.999 | .012 | .859 | .996 | >.999 | >.999 | −.010 | .841 | .019 | .811 | — |
| rh | 6r | −.014 | .597 | .975 | >.999 | >.999 | .001 | .994 | .997 | >.999 | >.999 | −.026 | .680 | .017 | .859 | — |
| rh | IFJa | .035 | .193 | .975 | >.999 | >.999 | .037 | .500 | .996 | >.999 | >.999 | .018 | .674 | .025 | .705 | — |
| rh | IFJp | .040 | .106 | .975 | >.999 | >.999 | .041 | .331 | .996 | >.999 | >.999 | .023 | .515 | .026 | .608 | — |
| rh | IFSp | −.046 | .796 | .975 | >.999 | >.999 | .057 | .446 | .996 | >.999 | >.999 | −.142 | **.014** | .149 | .092 | Matching |
| rh | IFSa | −.100 | .959 | .987 | >.999 | >.999 | −.090 | .266 | .996 | >.999 | >.999 | −.072 | .249 | −.043 | .651 | — |
| rh | p9-46v | −.022 | .624 | .975 | >.999 | >.999 | −.060 | .503 | .996 | >.999 | >.999 | .029 | .676 | −.079 | .460 | — |
| rh | 46 | −.005 | .520 | .975 | >.999 | >.999 | .005 | .950 | .996 | >.999 | >.999 | −.013 | .827 | .013 | .886 | — |
| rh | a9-46v | −.023 | .666 | .975 | >.999 | >.999 | −.023 | .753 | .996 | >.999 | >.999 | −.015 | .800 | −.013 | .880 | — |
| rh | 9-46d | −.038 | .752 | .975 | >.999 | >.999 | −.061 | .425 | .996 | >.999 | >.999 | .003 | .963 | −.062 | .487 | — |
| rh | 9a | .026 | .313 | .975 | >.999 | >.999 | .063 | .403 | .996 | >.999 | >.999 | −.025 | .660 | .079 | .376 | — |
| rh | 10v | .031 | .212 | .975 | >.999 | >.999 | .095 | .070 | .996 | >.999 | >.999 | −.052 | .222 | .129 | **.038** | AK+ |
| rh | a10p | .022 | .291 | .975 | >.999 | >.999 | .069 | .206 | .996 | >.999 | >.999 | −.038 | .396 | .093 | .153 | — |
| rh | 10pp | .008 | .418 | .975 | >.999 | >.999 | .047 | .380 | .996 | >.999 | >.999 | −.039 | .360 | .073 | .255 | — |
| rh | 11l | −.004 | .524 | .975 | >.999 | >.999 | .064 | .340 | .996 | >.999 | >.999 | −.077 | .138 | .113 | .149 | — |
| rh | 13l | .027 | .250 | .975 | >.999 | >.999 | .095 | .084 | .996 | >.999 | >.999 | −.059 | .182 | .133 | **.042** | AK+ |
| rh | OFC | .008 | .421 | .975 | >.999 | >.999 | .043 | .473 | .996 | >.999 | >.999 | −.034 | .473 | .066 | .361 | — |
| rh | 47s | .051 | .136 | .975 | .995 | >.999 | .080 | .202 | .996 | >.999 | >.999 | −.002 | .967 | .082 | .278 | — |
| rh | LIPd | −.015 | .696 | .975 | >.999 | >.999 | −.000 | .990 | .996 | >.999 | >.999 | −.025 | .450 | .016 | .731 | — |
| rh | 6a | −.073 | .932 | .984 | >.999 | >.999 | −.052 | .449 | .996 | >.999 | >.999 | −.068 | .197 | −.008 | .928 | — |
| rh | i6-8 | −.023 | .665 | .975 | >.999 | >.999 | −.069 | .325 | .996 | >.999 | >.999 | .038 | .490 | −.093 | .265 | — |
| rh | s6-8 | −.001 | .506 | .975 | >.999 | >.999 | .076 | .248 | .996 | >.999 | >.999 | −.087 | .093 | .132 | .089 | — |
| rh | 43 | .066 | .080 | .975 | .966 | >.999 | .161 | **.010** | .996 | .827 | .832 | −.066 | .192 | .203 | **.007** | AK+ |
| rh | OP4 | −.015 | .632 | .975 | >.999 | >.999 | .034 | .595 | .996 | >.999 | >.999 | −.063 | .206 | .074 | .318 | — |
| rh | OP1 | −.016 | .655 | .975 | >.999 | >.999 | .009 | .880 | .996 | >.999 | >.999 | −.037 | .409 | .032 | .623 | — |
| rh | OP2-3 | .036 | .147 | .975 | >.999 | >.999 | .025 | .578 | .996 | >.999 | >.999 | .033 | .387 | .004 | .948 | — |
| rh | 52 | −.024 | .778 | .975 | >.999 | >.999 | −.023 | .576 | .996 | >.999 | >.999 | −.016 | .653 | −.013 | .794 | — |
| rh | RI | −.054 | .969 | .987 | >.999 | >.999 | −.025 | .495 | .996 | >.999 | >.999 | −.065 | **.045** | .017 | .705 | Matching |
| rh | PFcm | −.067 | .946 | .984 | >.999 | >.999 | .020 | .735 | .996 | >.999 | >.999 | −.137 | **.003** | .109 | .122 | Matching |
| rh | PoI2 | .074 | .062 | .975 | .927 | >.999 | .117 | .070 | .996 | >.999 | >.999 | −.002 | .962 | .119 | .128 | — |
| rh | TA2 | .024 | .298 | .975 | >.999 | >.999 | .044 | .476 | .996 | >.999 | >.999 | −.008 | .877 | .049 | .505 | — |
| rh | FOP4 | −.042 | .835 | .979 | >.999 | >.999 | −.042 | .465 | .996 | >.999 | >.999 | −.025 | .592 | −.026 | .706 | — |
| rh | MI | .019 | .346 | .975 | >.999 | >.999 | .018 | .792 | .996 | >.999 | >.999 | .012 | .814 | .010 | .905 | — |
| rh | Pir | .025 | .288 | .975 | >.999 | >.999 | .067 | .266 | .996 | >.999 | >.999 | −.032 | .497 | .088 | .223 | — |
| rh | AVI | .003 | .462 | .975 | >.999 | >.999 | .056 | .313 | .996 | >.999 | >.999 | −.057 | .200 | .093 | .159 | — |
| rh | AAIC | .014 | .372 | .975 | >.999 | >.999 | .064 | .285 | .996 | >.999 | >.999 | −.046 | .324 | .094 | .187 | — |
| rh | FOP1 | −.027 | .751 | .975 | >.999 | >.999 | −.029 | .582 | .996 | >.999 | >.999 | −.014 | .747 | −.020 | .745 | — |
| rh | FOP3 | −.019 | .737 | .975 | >.999 | >.999 | .023 | .554 | .996 | >.999 | >.999 | −.059 | .081 | .062 | .195 | — |
| rh | FOP2 | −.038 | .870 | .981 | >.999 | >.999 | −.020 | .652 | .996 | >.999 | >.999 | −.042 | .258 | .007 | .889 | — |
| rh | PFt | −.044 | .834 | .979 | >.999 | >.999 | −.033 | .586 | .996 | >.999 | >.999 | −.038 | .425 | −.008 | .909 | — |
| rh | AIP | −.064 | .918 | .984 | >.999 | >.999 | −.070 | .270 | .996 | >.999 | >.999 | −.033 | .518 | −.048 | .522 | — |
| rh | EC | .079 | .062 | .975 | .899 | >.999 | .113 | .097 | .996 | >.999 | >.999 | .010 | .844 | .106 | .197 | — |
| rh | PreS | .027 | .251 | .975 | >.999 | >.999 | −.041 | .459 | .996 | >.999 | >.999 | .091 | **.038** | −.100 | .126 | Matching |
| rh | H | −.001 | .503 | .975 | >.999 | >.999 | −.012 | .833 | .996 | >.999 | >.999 | .012 | .787 | −.019 | .769 | — |
| rh | ProS | −.030 | .789 | .975 | >.999 | >.999 | .011 | .829 | .996 | >.999 | >.999 | −.063 | .118 | .052 | .385 | — |
| rh | PeEc | −.005 | .532 | .975 | >.999 | >.999 | .033 | .649 | .996 | >.999 | >.999 | −.046 | .418 | .063 | .470 | — |
| rh | STGa | .071 | .063 | .975 | .947 | >.999 | .146 | **.016** | .996 | .949 | .952 | −.041 | .402 | .173 | **.017** | AK+ |
| rh | PBelt | −.007 | .559 | .975 | >.999 | >.999 | .048 | .398 | .996 | >.999 | >.999 | −.066 | .146 | .091 | .182 | — |
| rh | A5 | .005 | .458 | .975 | >.999 | >.999 | .029 | .730 | .996 | >.999 | >.999 | −.024 | .714 | .044 | .661 | — |
| rh | PHA1 | .087 | **.042** | .975 | .841 | >.999 | .092 | .166 | .996 | >.999 | >.999 | .046 | .377 | .062 | .434 | — |
| rh | PHA3 | .019 | .279 | .975 | >.999 | >.999 | .044 | .297 | .996 | >.999 | >.999 | −.017 | .641 | .055 | .278 | — |
| rh | STSda | −.007 | .548 | .975 | >.999 | >.999 | .019 | .790 | .996 | >.999 | >.999 | −.034 | .551 | .041 | .632 | — |
| rh | STSdp | −.137 | .995 | .998 | >.999 | >.999 | −.112 | .131 | .996 | >.999 | >.999 | −.110 | **.050** | −.041 | .647 | Matching |
| rh | STSvp | −.012 | .577 | .975 | >.999 | >.999 | −.039 | .642 | .996 | >.999 | >.999 | .022 | .735 | −.053 | .593 | — |
| rh | TGd | .016 | .385 | .975 | >.999 | >.999 | .116 | .161 | .996 | >.999 | >.999 | −.101 | .104 | .181 | .062 | — |
| rh | TE1a | −.026 | .674 | .975 | >.999 | >.999 | .013 | .869 | .996 | >.999 | >.999 | −.059 | .315 | .051 | .572 | — |
| rh | TE1p | −.013 | .574 | .975 | >.999 | >.999 | −.016 | .848 | .996 | >.999 | >.999 | −.004 | .945 | −.013 | .891 | — |
| rh | TE2a | −.045 | .786 | .975 | >.999 | >.999 | −.002 | .984 | .996 | >.999 | >.999 | −.076 | .200 | .047 | .605 | — |
| rh | TF | .004 | .460 | .975 | >.999 | >.999 | .003 | .963 | .996 | >.999 | >.999 | .004 | .943 | .001 | .992 | — |
| rh | TE2p | −.044 | .809 | .975 | >.999 | >.999 | −.028 | .686 | .996 | >.999 | >.999 | −.044 | .413 | .001 | .993 | — |
| rh | PHT | .003 | .480 | .975 | >.999 | >.999 | .021 | .800 | .996 | >.999 | >.999 | −.019 | .768 | .033 | .739 | — |
| rh | PH | −.016 | .637 | .975 | >.999 | >.999 | −.011 | .862 | .996 | >.999 | >.999 | −.016 | .737 | −.001 | .994 | — |
| rh | TPOJ1 | −.020 | .618 | .975 | >.999 | >.999 | −.018 | .837 | .996 | >.999 | >.999 | −.015 | .829 | −.008 | .936 | — |
| rh | TPOJ2 | .025 | .333 | .975 | >.999 | >.999 | .024 | .764 | .996 | >.999 | >.999 | .015 | .805 | .014 | .878 | — |
| rh | TPOJ3 | −.003 | .522 | .975 | >.999 | >.999 | .047 | .390 | .996 | >.999 | >.999 | −.057 | .193 | .084 | .196 | — |
| rh | DVT | −.021 | .659 | .975 | >.999 | >.999 | −.044 | .525 | .996 | >.999 | >.999 | .012 | .826 | −.051 | .528 | — |
| rh | PGp | .030 | .278 | .975 | >.999 | >.999 | .018 | .799 | .996 | >.999 | >.999 | .032 | .558 | −.003 | .970 | — |
| rh | IP2 | −.071 | .936 | .984 | >.999 | >.999 | −.052 | .426 | .996 | >.999 | >.999 | −.064 | .208 | −.011 | .892 | — |
| rh | IP1 | −.013 | .614 | .975 | >.999 | >.999 | −.022 | .723 | .996 | >.999 | >.999 | .002 | .976 | −.023 | .760 | — |
| rh | IP0 | −.003 | .529 | .975 | >.999 | >.999 | −.012 | .849 | .996 | >.999 | >.999 | .007 | .884 | −.016 | .825 | — |
| rh | PFop | −.049 | .866 | .980 | >.999 | >.999 | −.002 | .974 | .996 | >.999 | >.999 | −.082 | .088 | .051 | .481 | — |
| rh | PF | −.026 | .689 | .975 | >.999 | >.999 | −.018 | .809 | .996 | >.999 | >.999 | −.025 | .669 | −.002 | .980 | — |
| rh | PFm | −.063 | .860 | .980 | >.999 | >.999 | −.091 | .264 | .996 | >.999 | >.999 | −.007 | .911 | −.087 | .372 | — |
| rh | PGi | .020 | .364 | .975 | >.999 | >.999 | −.026 | .751 | .996 | >.999 | >.999 | .064 | .317 | −.068 | .491 | — |
| rh | PGs | −.026 | .682 | .975 | >.999 | >.999 | −.116 | .121 | .996 | >.999 | >.999 | .084 | .145 | −.170 | .052 | — |
| rh | V6A | −.002 | .514 | .975 | >.999 | >.999 | .006 | .902 | .996 | >.999 | >.999 | −.010 | .806 | .012 | .836 | — |
| rh | VMV1 | .016 | .355 | .975 | >.999 | >.999 | .040 | .515 | .996 | >.999 | >.999 | −.017 | .735 | .051 | .486 | — |
| rh | VMV3 | .017 | .323 | .975 | >.999 | >.999 | .044 | .380 | .996 | >.999 | >.999 | −.020 | .625 | .056 | .343 | — |
| rh | PHA2 | −.035 | .848 | .979 | >.999 | >.999 | −.030 | .506 | .996 | >.999 | >.999 | −.027 | .480 | −.013 | .815 | — |
| rh | V4t | −.042 | .862 | .980 | >.999 | >.999 | .007 | .890 | .996 | >.999 | >.999 | −.081 | .056 | .060 | .340 | — |
| rh | FST | −.024 | .707 | .975 | >.999 | >.999 | −.082 | .157 | .996 | >.999 | >.999 | .051 | .276 | −.115 | .096 | — |
| rh | V3CD | −.015 | .637 | .975 | >.999 | >.999 | −.012 | .823 | .996 | >.999 | >.999 | −.011 | .795 | −.005 | .941 | — |
| rh | LO3 | −.052 | .879 | .984 | >.999 | >.999 | −.049 | .411 | .996 | >.999 | >.999 | −.034 | .481 | −.027 | .705 | — |
| rh | VMV2 | .029 | .185 | .975 | >.999 | >.999 | .022 | .616 | .996 | >.999 | >.999 | .026 | .476 | .005 | .925 | — |
| rh | 31pd | .064 | .060 | .975 | .972 | >.999 | .015 | .779 | .996 | >.999 | >.999 | .093 | **.034** | −.045 | .502 | Matching |
| rh | 31a | −.030 | .739 | .975 | >.999 | >.999 | −.049 | .420 | .996 | >.999 | >.999 | .003 | .946 | −.051 | .480 | — |
| rh | VVC | .001 | .490 | .975 | >.999 | >.999 | .027 | .691 | .996 | >.999 | >.999 | −.029 | .589 | .046 | .568 | — |
| rh | 25 | .073 | **.017** | .975 | .936 | >.999 | .070 | .129 | .996 | >.999 | >.999 | .048 | .210 | .039 | .480 | — |
| rh | s32 | .004 | .452 | .975 | >.999 | >.999 | .037 | .375 | .996 | >.999 | >.999 | −.035 | .318 | .060 | .234 | — |
| rh | pOFC | −.012 | .603 | .975 | >.999 | >.999 | .071 | .214 | .996 | >.999 | >.999 | −.099 | **.030** | .135 | **.045** | Match + AK+ |
| rh | PoI1 | .030 | .226 | .975 | >.999 | >.999 | .026 | .625 | .996 | >.999 | >.999 | .022 | .599 | .011 | .860 | — |
| rh | Ig | .019 | .279 | .975 | >.999 | >.999 | .004 | .928 | .996 | >.999 | >.999 | .028 | .433 | −.014 | .776 | — |
| rh | FOP5 | .018 | .334 | .975 | >.999 | >.999 | .048 | .403 | .996 | >.999 | >.999 | −.023 | .610 | .063 | .354 | — |
| rh | p10p | .025 | .285 | .975 | >.999 | >.999 | .011 | .857 | .996 | >.999 | >.999 | .030 | .538 | −.008 | .913 | — |
| rh | p47r | −.018 | .631 | .975 | >.999 | >.999 | .001 | .990 | .996 | >.999 | >.999 | −.032 | .548 | .021 | .790 | — |
| rh | TGv | −.043 | .806 | .975 | >.999 | >.999 | −.013 | .848 | .996 | >.999 | >.999 | −.060 | .261 | .026 | .751 | — |
| rh | MBelt | −.008 | .586 | .975 | >.999 | >.999 | .035 | .469 | .996 | >.999 | >.999 | −.053 | .193 | .069 | .235 | — |
| rh | LBelt | −.027 | .756 | .975 | >.999 | >.999 | .001 | .981 | .996 | >.999 | >.999 | −.049 | .258 | .033 | .612 | — |
| rh | A4 | .099 | **.037** | .975 | .716 | >.999 | .177 | **.016** | .996 | .631 | .637 | −.027 | .634 | .194 | **.026** | AK+ |
| rh | STSva | .008 | .431 | .975 | >.999 | >.999 | .056 | .443 | .996 | >.999 | >.999 | −.048 | .400 | .087 | .317 | — |
| rh | TE1m | .066 | .119 | .975 | .965 | >.999 | .137 | .068 | .996 | .986 | .988 | −.038 | .515 | .162 | .070 | — |
| rh | PI | .008 | .422 | .975 | >.999 | >.999 | .003 | .965 | .996 | >.999 | >.999 | .011 | .805 | −.005 | .944 | — |
| rh | a32pr | −.048 | .863 | .980 | >.999 | >.999 | −.064 | .282 | .996 | >.999 | >.999 | −.011 | .818 | −.057 | .426 | — |
| rh | p24 | .047 | .155 | .975 | .997 | >.999 | .031 | .628 | .996 | >.999 | >.999 | .046 | .347 | .001 | .992 | — |

**Retrospective ELS — Adjusted for sex, ART, age, SES (N = 85, 20,000 permutations, 360 parcels, r(|a − b|, mean) = .65)**

|  | | **Matching ρ(D, \|a − b\|)** | | | | | **Additive ρ(D, mean(a, b))** | | | | | **Joint model D ~ \|a − b\| + mean(a, b)** | | | |  |
| --- | --- | --- | --- | --- | --- | --- | --- | --- | --- | --- | --- | --- | --- | --- | --- | --- |
| **Hemi** | **Region** | **ρ** | **p** | **q** | **p FWE-P** | **p FWE-F** | **ρ** | **p** | **q** | **p FWE-P** | **p FWE-F** | **β \|a−b\|** | **p** | **β mean** | **p** | **Mechanism** |
| lh | V1 | −.047 | .793 | .964 | >.999 | >.999 | .010 | .903 | .996 | >.999 | >.999 | −.091 | .125 | .069 | .458 | — |
| lh | MST | .019 | .253 | .964 | >.999 | >.999 | .040 | .296 | .996 | >.999 | >.999 | −.011 | .745 | .047 | .302 | — |
| lh | V6 | −.013 | .613 | .964 | >.999 | >.999 | .006 | .920 | .996 | >.999 | >.999 | −.029 | .527 | .025 | .716 | — |
| lh | V2 | .017 | .375 | .964 | >.999 | >.999 | .058 | .482 | .996 | >.999 | >.999 | −.034 | .578 | .080 | .406 | — |
| lh | V3 | .020 | .362 | .964 | >.999 | >.999 | .079 | .317 | .996 | >.999 | >.999 | −.054 | .369 | .114 | .222 | — |
| lh | V4 | −.060 | .848 | .973 | >.999 | >.999 | .018 | .828 | .996 | >.999 | >.999 | −.123 | **.044** | .098 | .309 | Matching |
| lh | V8 | .029 | .273 | .964 | >.999 | >.999 | .049 | .482 | .996 | >.999 | >.999 | −.004 | .943 | .051 | .533 | — |
| lh | 4 | .016 | .388 | .964 | >.999 | >.999 | .042 | .604 | .996 | >.999 | >.999 | −.020 | .747 | .055 | .564 | — |
| lh | 3b | .012 | .400 | .964 | >.999 | >.999 | −.013 | .855 | .996 | >.999 | >.999 | .035 | .525 | −.036 | .681 | — |
| lh | FEF | .048 | .164 | .964 | .996 | >.999 | .017 | .800 | .996 | >.999 | >.999 | .063 | .223 | −.024 | .767 | — |
| lh | PEF | .004 | .465 | .964 | >.999 | >.999 | −.002 | .975 | .996 | >.999 | >.999 | .008 | .862 | −.007 | .922 | — |
| lh | 55b | −.038 | .743 | .964 | >.999 | >.999 | −.008 | .922 | .996 | >.999 | >.999 | −.056 | .348 | .028 | .761 | — |
| lh | V3A | .036 | .202 | .964 | >.999 | >.999 | .090 | .132 | .996 | >.999 | >.999 | −.037 | .430 | .114 | .106 | — |
| lh | RSC | .008 | .426 | .964 | >.999 | >.999 | −.048 | .388 | .996 | >.999 | >.999 | .066 | .136 | −.091 | .169 | — |
| lh | POS2 | −.047 | .806 | .964 | >.999 | >.999 | −.025 | .739 | .996 | >.999 | >.999 | −.053 | .339 | .010 | .909 | — |
| lh | V7 | −.008 | .590 | .964 | >.999 | >.999 | .003 | .942 | .996 | >.999 | >.999 | −.017 | .657 | .015 | .794 | — |
| lh | IPS1 | −.013 | .597 | .964 | >.999 | >.999 | .014 | .839 | .996 | >.999 | >.999 | −.037 | .478 | .038 | .632 | — |
| lh | FFC | −.012 | .570 | .964 | >.999 | >.999 | −.003 | .971 | .996 | >.999 | >.999 | −.017 | .781 | .008 | .930 | — |
| lh | V3B | .011 | .374 | .964 | >.999 | >.999 | .014 | .757 | .996 | >.999 | >.999 | .004 | .917 | .011 | .834 | — |
| lh | LO1 | .064 | .058 | .964 | .973 | >.999 | .094 | .082 | .996 | >.999 | >.999 | .005 | .916 | .091 | .155 | — |
| lh | LO2 | .004 | .467 | .964 | >.999 | >.999 | .011 | .864 | .996 | >.999 | >.999 | −.006 | .912 | .014 | .848 | — |
| lh | PIT | −.090 | .971 | .989 | >.999 | >.999 | −.018 | .784 | .996 | >.999 | >.999 | −.134 | **.010** | .068 | .392 | Matching |
| lh | MT | .056 | .051 | .964 | .990 | >.999 | .072 | .114 | .996 | >.999 | >.999 | .016 | .668 | .062 | .262 | — |
| lh | A1 | .012 | .377 | .964 | >.999 | >.999 | .042 | .439 | .996 | >.999 | >.999 | −.027 | .532 | .060 | .353 | — |
| lh | PSL | −.078 | .904 | .988 | >.999 | >.999 | −.075 | .369 | .996 | >.999 | >.999 | −.051 | .416 | −.042 | .672 | — |
| lh | SFL | −.036 | .716 | .964 | >.999 | >.999 | −.025 | .762 | .996 | >.999 | >.999 | −.034 | .590 | −.003 | .974 | — |
| lh | PCV | .006 | .437 | .964 | >.999 | >.999 | −.006 | .914 | .996 | >.999 | >.999 | .018 | .708 | −.018 | .801 | — |
| lh | STV | −.081 | .909 | .988 | >.999 | >.999 | −.090 | .289 | .996 | >.999 | >.999 | −.038 | .549 | −.065 | .513 | — |
| lh | 7Pm | −.003 | .531 | .964 | >.999 | >.999 | −.022 | .703 | .996 | >.999 | >.999 | .019 | .687 | −.034 | .626 | — |
| lh | 7m | .029 | .295 | .964 | >.999 | >.999 | −.035 | .644 | .996 | >.999 | >.999 | .089 | .119 | −.092 | .301 | — |
| lh | POS1 | −.003 | .516 | .964 | >.999 | >.999 | .035 | .606 | .996 | >.999 | >.999 | −.043 | .411 | .063 | .431 | — |
| lh | 23d | −.007 | .557 | .964 | >.999 | >.999 | −.001 | .985 | .996 | >.999 | >.999 | −.010 | .832 | .005 | .942 | — |
| lh | v23ab | −.024 | .734 | .964 | >.999 | >.999 | −.119 | **.024** | .996 | >.999 | >.999 | .091 | **.030** | −.178 | **.005** | Match + AK− |
| lh | d23ab | .034 | .242 | .964 | >.999 | >.999 | .004 | .960 | .996 | >.999 | >.999 | .054 | .298 | −.031 | .701 | — |
| lh | 31pv | −.052 | .898 | .986 | >.999 | >.999 | −.142 | **.010** | .996 | .972 | .974 | .068 | .131 | −.186 | **.005** | AK− |
| lh | 5m | −.010 | .603 | .964 | >.999 | >.999 | .006 | .900 | .996 | >.999 | >.999 | −.024 | .551 | .022 | .718 | — |
| lh | 5mv | −.030 | .789 | .964 | >.999 | >.999 | −.010 | .837 | .996 | >.999 | >.999 | −.041 | .327 | .016 | .795 | — |
| lh | 23c | −.011 | .597 | .964 | >.999 | >.999 | .008 | .887 | .996 | >.999 | >.999 | −.027 | .542 | .026 | .697 | — |
| lh | 5L | −.026 | .726 | .964 | >.999 | >.999 | −.051 | .370 | .996 | >.999 | >.999 | .013 | .784 | −.059 | .381 | — |
| lh | 24dd | −.015 | .610 | .964 | >.999 | >.999 | −.016 | .817 | .996 | >.999 | >.999 | −.008 | .872 | −.010 | .899 | — |
| lh | 24dv | −.055 | .926 | .988 | >.999 | >.999 | −.043 | .415 | .996 | >.999 | >.999 | −.047 | .259 | −.012 | .845 | — |
| lh | 7AL | −.041 | .813 | .964 | >.999 | >.999 | −.023 | .715 | .996 | >.999 | >.999 | −.045 | .355 | .006 | .940 | — |
| lh | SCEF | −.013 | .579 | .964 | >.999 | >.999 | .047 | .557 | .996 | >.999 | >.999 | −.074 | .210 | .095 | .311 | — |
| lh | 6ma | −.038 | .747 | .964 | >.999 | >.999 | −.044 | .562 | .996 | >.999 | >.999 | −.016 | .788 | −.034 | .709 | — |
| lh | 7Am | .012 | .404 | .964 | >.999 | >.999 | .018 | .795 | .996 | >.999 | >.999 | .000 | .996 | .018 | .829 | — |
| lh | 7Pl | .032 | .219 | .964 | >.999 | >.999 | .038 | .512 | .996 | >.999 | >.999 | .013 | .776 | .029 | .668 | — |
| lh | 7PC | −.004 | .530 | .964 | >.999 | >.999 | .042 | .553 | .996 | >.999 | >.999 | −.054 | .320 | .077 | .355 | — |
| lh | LIPv | −.052 | .863 | .979 | >.999 | >.999 | −.011 | .868 | .996 | >.999 | >.999 | −.078 | .126 | .040 | .609 | — |
| lh | VIP | −.068 | .917 | .988 | >.999 | >.999 | −.050 | .464 | .996 | >.999 | >.999 | −.062 | .244 | −.010 | .906 | — |
| lh | MIP | −.027 | .720 | .964 | >.999 | >.999 | −.006 | .933 | .996 | >.999 | >.999 | −.041 | .418 | .021 | .786 | — |
| lh | 1 | .003 | .466 | .964 | >.999 | >.999 | .030 | .696 | .996 | >.999 | >.999 | −.028 | .636 | .048 | .601 | — |
| lh | 2 | −.084 | .956 | .988 | >.999 | >.999 | −.038 | .587 | .996 | >.999 | >.999 | −.102 | .055 | .028 | .738 | — |
| lh | 3a | .051 | .128 | .964 | .994 | >.999 | .048 | .414 | .996 | >.999 | >.999 | .034 | .475 | .026 | .709 | — |
| lh | 6d | −.005 | .529 | .964 | >.999 | >.999 | .078 | .256 | .996 | >.999 | >.999 | −.095 | .070 | .140 | .082 | — |
| lh | 6mp | −.028 | .728 | .964 | >.999 | >.999 | −.002 | .975 | .996 | >.999 | >.999 | −.045 | .351 | .027 | .705 | — |
| lh | 6v | .035 | .238 | .964 | >.999 | >.999 | .120 | .080 | .996 | >.999 | >.999 | −.073 | .170 | .167 | **.039** | AK+ |
| lh | p24pr | .010 | .399 | .964 | >.999 | >.999 | −.023 | .690 | .996 | >.999 | >.999 | .043 | .351 | −.052 | .460 | — |
| lh | 33pr | .001 | .488 | .964 | >.999 | >.999 | .044 | .302 | .996 | >.999 | >.999 | −.048 | .183 | .075 | .143 | — |
| lh | a24pr | −.045 | .874 | .983 | >.999 | >.999 | −.010 | .858 | .996 | >.999 | >.999 | −.066 | .119 | .033 | .598 | — |
| lh | p32pr | .011 | .393 | .964 | >.999 | >.999 | .009 | .864 | .996 | >.999 | >.999 | .008 | .847 | .004 | .949 | — |
| lh | a24 | .041 | .162 | .964 | .998 | >.999 | .076 | .194 | .996 | >.999 | >.999 | −.014 | .765 | .084 | .222 | — |
| lh | d32 | .016 | .364 | .964 | >.999 | >.999 | −.045 | .500 | .996 | >.999 | >.999 | .077 | .134 | −.095 | .228 | — |
| lh | 8BM | −.089 | .942 | .988 | >.999 | >.999 | −.158 | **.046** | .996 | .858 | .863 | .023 | .699 | −.174 | .063 | — |
| lh | p32 | −.019 | .717 | .964 | >.999 | >.999 | −.015 | .745 | .996 | >.999 | >.999 | −.017 | .656 | −.004 | .936 | — |
| lh | 10r | −.040 | .847 | .973 | >.999 | >.999 | −.028 | .594 | .996 | >.999 | >.999 | −.038 | .378 | −.004 | .948 | — |
| lh | 47m | −.042 | .860 | .979 | >.999 | >.999 | .001 | .981 | .996 | >.999 | >.999 | −.073 | .082 | .048 | .429 | — |
| lh | 8Av | −.032 | .690 | .964 | >.999 | >.999 | −.053 | .533 | .996 | >.999 | >.999 | .004 | .955 | −.056 | .586 | — |
| lh | 8Ad | .004 | .457 | .964 | >.999 | >.999 | .026 | .701 | .996 | >.999 | >.999 | −.021 | .685 | .039 | .619 | — |
| lh | 9m | .010 | .420 | .964 | >.999 | >.999 | .034 | .685 | .996 | >.999 | >.999 | −.021 | .742 | .047 | .633 | — |
| lh | 8BL | −.005 | .518 | .964 | >.999 | >.999 | −.014 | .845 | .996 | >.999 | >.999 | .008 | .888 | −.019 | .822 | — |
| lh | 9p | −.012 | .583 | .964 | >.999 | >.999 | .033 | .629 | .996 | >.999 | >.999 | −.058 | .277 | .071 | .383 | — |
| lh | 10d | .023 | .289 | .964 | >.999 | >.999 | .045 | .445 | .996 | >.999 | >.999 | −.011 | .823 | .052 | .460 | — |
| lh | 8C | −.056 | .821 | .964 | >.999 | >.999 | −.053 | .526 | .996 | >.999 | >.999 | −.038 | .558 | −.029 | .770 | — |
| lh | 44 | .045 | .231 | .964 | .997 | >.999 | .134 | .114 | .996 | .992 | .993 | −.072 | .262 | .181 | .070 | — |
| lh | 45 | −.020 | .601 | .964 | >.999 | >.999 | −.041 | .649 | .996 | >.999 | >.999 | .012 | .855 | −.049 | .641 | — |
| lh | 47l | −.029 | .695 | .964 | >.999 | >.999 | −.008 | .913 | .996 | >.999 | >.999 | −.042 | .469 | .019 | .830 | — |
| lh | a47r | .011 | .399 | .964 | >.999 | >.999 | .076 | .278 | .996 | >.999 | >.999 | −.066 | .219 | .118 | .150 | — |
| lh | 6r | .022 | .345 | .964 | >.999 | >.999 | .092 | .242 | .996 | >.999 | >.999 | −.065 | .283 | .134 | .148 | — |
| lh | IFJa | −.010 | .576 | .964 | >.999 | >.999 | −.002 | .980 | .996 | >.999 | >.999 | −.015 | .774 | .008 | .923 | — |
| lh | IFJp | −.037 | .814 | .964 | >.999 | >.999 | −.011 | .844 | .996 | >.999 | >.999 | −.052 | .244 | .023 | .732 | — |
| lh | IFSp | .017 | .372 | .964 | >.999 | >.999 | .037 | .632 | .996 | >.999 | >.999 | −.011 | .848 | .044 | .623 | — |
| lh | IFSaᵃ | −.091 | .922 | .988 | >.999 | >.999 | −.052 | .559 | .996 | >.999 | >.999 | −.098 | .142 | .011 | .913 | — |
| lh | p9-46v | −.028 | .659 | .964 | >.999 | >.999 | −.085 | .325 | .996 | >.999 | >.999 | .047 | .474 | −.115 | .259 | — |
| lh | 46 | .015 | .380 | .964 | >.999 | >.999 | .068 | .370 | .996 | >.999 | >.999 | −.051 | .380 | .101 | .255 | — |
| lh | a9-46v | .016 | .373 | .964 | >.999 | >.999 | .021 | .781 | .996 | >.999 | >.999 | .005 | .929 | .017 | .850 | — |
| lh | 9-46d | −.012 | .571 | .964 | >.999 | >.999 | −.014 | .858 | .996 | >.999 | >.999 | −.005 | .926 | −.010 | .912 | — |
| lh | 9a | −.033 | .705 | .964 | >.999 | >.999 | −.018 | .826 | .996 | >.999 | >.999 | −.038 | .537 | .007 | .941 | — |
| lh | 10v | −.010 | .579 | .964 | >.999 | >.999 | .054 | .334 | .996 | >.999 | >.999 | −.077 | .083 | .104 | .122 | — |
| lh | a10p | −.025 | .727 | .964 | >.999 | >.999 | .003 | .950 | .996 | >.999 | >.999 | −.047 | .287 | .034 | .616 | — |
| lh | 10pp | −.050 | .912 | .988 | >.999 | >.999 | −.009 | .855 | .996 | >.999 | >.999 | −.076 | .059 | .041 | .490 | — |
| lh | 11l | −.042 | .817 | .964 | >.999 | >.999 | .018 | .782 | .996 | >.999 | >.999 | −.092 | .062 | .077 | .310 | — |
| lh | 13l | .012 | .394 | .964 | >.999 | >.999 | .091 | .150 | .996 | >.999 | >.999 | −.081 | .096 | .144 | .054 | — |
| lh | OFC | .035 | .208 | .964 | >.999 | >.999 | .062 | .296 | .996 | >.999 | >.999 | −.009 | .850 | .067 | .344 | — |
| lh | 47s | −.011 | .589 | .964 | >.999 | >.999 | .057 | .332 | .996 | >.999 | >.999 | −.082 | .079 | .110 | .115 | — |
| lh | LIPd | .011 | .379 | .964 | >.999 | >.999 | −.051 | .311 | .996 | >.999 | >.999 | .077 | .057 | −.101 | .090 | — |
| lh | 6a | −.019 | .657 | .964 | >.999 | >.999 | −.002 | .969 | .996 | >.999 | >.999 | −.029 | .558 | .016 | .826 | — |
| lh | i6-8 | .011 | .402 | .964 | >.999 | >.999 | .028 | .655 | .996 | >.999 | >.999 | −.012 | .802 | .035 | .630 | — |
| lh | s6-8 | −.022 | .700 | .964 | >.999 | >.999 | −.035 | .518 | .996 | >.999 | >.999 | .001 | .981 | −.036 | .575 | — |
| lh | 43 | −.017 | .662 | .964 | >.999 | >.999 | .013 | .816 | .996 | >.999 | >.999 | −.043 | .323 | .041 | .531 | — |
| lh | OP4 | −.039 | .785 | .964 | >.999 | >.999 | .037 | .590 | .996 | >.999 | >.999 | −.107 | **.041** | .106 | .185 | Matching |
| lh | OP1 | −.027 | .735 | .964 | >.999 | >.999 | .022 | .693 | .996 | >.999 | >.999 | −.071 | .118 | .068 | .321 | — |
| lh | OP2-3 | −.041 | .888 | .986 | >.999 | >.999 | −.020 | .649 | .996 | >.999 | >.999 | −.048 | .196 | .011 | .835 | — |
| lh | 52 | .039 | .139 | .964 | .999 | >.999 | .090 | .061 | .996 | >.999 | >.999 | −.032 | .421 | .110 | .052 | — |
| lh | RI | .031 | .201 | .964 | >.999 | >.999 | .064 | .210 | .996 | >.999 | >.999 | −.017 | .676 | .075 | .214 | — |
| lh | PFcm | .007 | .431 | .964 | >.999 | >.999 | .035 | .573 | .996 | >.999 | >.999 | −.027 | .582 | .053 | .476 | — |
| lh | PoI2 | .012 | .389 | .964 | >.999 | >.999 | .031 | .620 | .996 | >.999 | >.999 | −.014 | .789 | .040 | .589 | — |
| lh | TA2 | −.027 | .744 | .964 | >.999 | >.999 | −.001 | .985 | .996 | >.999 | >.999 | −.045 | .302 | .028 | .658 | — |
| lh | FOP4 | .025 | .295 | .964 | >.999 | >.999 | .052 | .411 | .996 | >.999 | >.999 | −.016 | .755 | .063 | .409 | — |
| lh | MI | .041 | .193 | .964 | .998 | >.999 | .084 | .199 | .996 | >.999 | >.999 | −.022 | .668 | .098 | .203 | — |
| lh | Pir | .030 | .221 | .964 | >.999 | >.999 | .085 | .115 | .996 | >.999 | >.999 | −.042 | .327 | .112 | .078 | — |
| lh | AVI | .010 | .411 | .964 | >.999 | >.999 | .017 | .785 | .996 | >.999 | >.999 | −.002 | .968 | .018 | .807 | — |
| lh | AAIC | .020 | .309 | .964 | >.999 | >.999 | .026 | .656 | .996 | >.999 | >.999 | .006 | .891 | .022 | .753 | — |
| lh | FOP1 | .001 | .494 | .964 | >.999 | >.999 | .056 | .316 | .996 | >.999 | >.999 | −.061 | .166 | .095 | .147 | — |
| lh | FOP3 | .024 | .252 | .964 | >.999 | >.999 | .030 | .525 | .996 | >.999 | >.999 | .008 | .842 | .025 | .656 | — |
| lh | FOP2 | .029 | .184 | .964 | >.999 | >.999 | .067 | .120 | .996 | >.999 | >.999 | −.025 | .507 | .083 | .108 | — |
| lh | PFt | −.020 | .683 | .964 | >.999 | >.999 | .040 | .456 | .996 | >.999 | >.999 | −.078 | .068 | .090 | .157 | — |
| lh | AIP | −.119 | .998 | .998 | >.999 | >.999 | −.100 | .088 | .996 | >.999 | >.999 | −.093 | **.045** | −.040 | .572 | Matching |
| lh | EC | .009 | .416 | .964 | >.999 | >.999 | .068 | .290 | .996 | >.999 | >.999 | −.059 | .239 | .106 | .160 | — |
| lh | PreS | −.018 | .651 | .964 | >.999 | >.999 | −.003 | .961 | .996 | >.999 | >.999 | −.028 | .563 | .015 | .834 | — |
| lh | H | −.059 | .921 | .988 | >.999 | >.999 | −.058 | .318 | .996 | >.999 | >.999 | −.037 | .417 | −.033 | .628 | — |
| lh | ProS | −.008 | .593 | .964 | >.999 | >.999 | .012 | .795 | .996 | >.999 | >.999 | −.027 | .471 | .029 | .588 | — |
| lh | PeEc | −.080 | .930 | .988 | >.999 | >.999 | −.031 | .686 | .996 | >.999 | >.999 | −.102 | .077 | .035 | .700 | — |
| lh | STGa | −.021 | .656 | .964 | >.999 | >.999 | −.005 | .943 | .996 | >.999 | >.999 | −.030 | .568 | .014 | .861 | — |
| lh | PBelt | .051 | .102 | .964 | .994 | >.999 | .066 | .228 | .996 | >.999 | >.999 | .015 | .738 | .057 | .385 | — |
| lh | A5 | −.057 | .811 | .964 | >.999 | >.999 | −.033 | .712 | .996 | >.999 | >.999 | −.062 | .350 | .008 | .942 | — |
| lh | PHA1 | .051 | .118 | .964 | .994 | >.999 | .078 | .192 | .996 | >.999 | >.999 | .002 | .970 | .077 | .279 | — |
| lh | PHA3 | −.034 | .765 | .964 | >.999 | >.999 | −.002 | .974 | .996 | >.999 | >.999 | −.056 | .267 | .034 | .655 | — |
| lh | STSda | −.015 | .610 | .964 | >.999 | >.999 | −.032 | .637 | .996 | >.999 | >.999 | .010 | .848 | −.039 | .632 | — |
| lh | STSdp | −.037 | .758 | .964 | >.999 | >.999 | −.001 | .984 | .996 | >.999 | >.999 | −.061 | .251 | .038 | .648 | — |
| lh | STSvp | −.132 | .989 | .994 | >.999 | >.999 | −.114 | .172 | .996 | >.999 | >.999 | −.099 | .117 | −.050 | .621 | — |
| lh | TGd | −.011 | .560 | .964 | >.999 | >.999 | .050 | .546 | .996 | >.999 | >.999 | −.074 | .242 | .099 | .324 | — |
| lh | TE1a | −.024 | .645 | .964 | >.999 | >.999 | .035 | .671 | .996 | >.999 | >.999 | −.080 | .192 | .087 | .366 | — |
| lh | TE1p | −.005 | .517 | .964 | >.999 | >.999 | −.032 | .715 | .996 | >.999 | >.999 | .027 | .679 | −.049 | .632 | — |
| lh | TE2a | −.013 | .573 | .964 | >.999 | >.999 | .054 | .493 | .996 | >.999 | >.999 | −.083 | .166 | .108 | .248 | — |
| lh | TF | −.042 | .774 | .964 | >.999 | >.999 | −.043 | .572 | .996 | >.999 | >.999 | −.025 | .671 | −.027 | .768 | — |
| lh | TE2p | −.023 | .643 | .964 | >.999 | >.999 | −.037 | .632 | .996 | >.999 | >.999 | .002 | .968 | −.039 | .670 | — |
| lh | PHT | −.081 | .895 | .986 | >.999 | >.999 | −.096 | .286 | .996 | >.999 | >.999 | −.032 | .640 | −.076 | .477 | — |
| lh | PH | .007 | .438 | .964 | >.999 | >.999 | .046 | .537 | .996 | >.999 | >.999 | −.039 | .501 | .071 | .418 | — |
| lh | TPOJ1 | −.019 | .618 | .964 | >.999 | >.999 | −.022 | .777 | .996 | >.999 | >.999 | −.008 | .893 | −.016 | .860 | — |
| lh | TPOJ2 | −.028 | .680 | .964 | >.999 | >.999 | −.022 | .784 | .996 | >.999 | >.999 | −.024 | .692 | −.006 | .946 | — |
| lh | TPOJ3 | .017 | .310 | .964 | >.999 | >.999 | .077 | .109 | .996 | >.999 | >.999 | −.056 | .155 | .113 | **.047** | AK+ |
| lh | DVT | −.033 | .756 | .964 | >.999 | >.999 | −.033 | .615 | .996 | >.999 | >.999 | −.021 | .678 | −.019 | .796 | — |
| lh | PGp | .013 | .392 | .964 | >.999 | >.999 | .048 | .493 | .996 | >.999 | >.999 | −.031 | .567 | .069 | .414 | — |
| lh | IP2 | −.000 | .502 | .964 | >.999 | >.999 | −.048 | .389 | .996 | >.999 | >.999 | .053 | .237 | −.083 | .215 | — |
| lh | IP1 | .020 | .328 | .964 | >.999 | >.999 | .015 | .812 | .996 | >.999 | >.999 | .018 | .706 | .003 | .966 | — |
| lh | IP0 | −.070 | .949 | .988 | >.999 | >.999 | −.070 | .235 | .996 | >.999 | >.999 | −.043 | .371 | −.043 | .555 | — |
| lh | PFop | −.034 | .782 | .964 | >.999 | >.999 | .008 | .900 | .996 | >.999 | >.999 | −.067 | .141 | .051 | .467 | — |
| lh | PF | −.053 | .814 | .964 | >.999 | >.999 | −.027 | .740 | .996 | >.999 | >.999 | −.061 | .329 | .012 | .899 | — |
| lh | PFm | −.074 | .899 | .986 | >.999 | >.999 | −.072 | .378 | .996 | >.999 | >.999 | −.047 | .442 | −.041 | .668 | — |
| lh | PGi | −.111 | .968 | .989 | >.999 | >.999 | −.108 | .208 | .996 | >.999 | >.999 | −.071 | .274 | −.062 | .544 | — |
| lh | PGsᵃ | −.077 | .942 | .988 | >.999 | >.999 | −.063 | .354 | .996 | >.999 | >.999 | −.062 | .246 | −.023 | .774 | — |
| lh | V6A | −.021 | .723 | .964 | >.999 | >.999 | .028 | .548 | .996 | >.999 | >.999 | −.068 | .076 | .072 | .193 | — |
| lh | VMV1 | .007 | .425 | .964 | >.999 | >.999 | .047 | .347 | .996 | >.999 | >.999 | −.040 | .326 | .073 | .219 | — |
| lh | VMV3 | −.031 | .786 | .964 | >.999 | >.999 | −.029 | .578 | .996 | >.999 | >.999 | −.021 | .626 | −.016 | .800 | — |
| lh | PHA2 | .026 | .187 | .964 | >.999 | >.999 | .063 | .098 | .996 | >.999 | >.999 | −.025 | .448 | .079 | .086 | — |
| lh | V4t | −.023 | .722 | .964 | >.999 | >.999 | .022 | .665 | .996 | >.999 | >.999 | −.064 | .117 | .063 | .290 | — |
| lh | FST | .002 | .478 | .964 | >.999 | >.999 | .103 | .129 | .996 | >.999 | >.999 | −.111 | **.034** | .174 | **.029** | Match + AK+ |
| lh | V3CD | −.008 | .569 | .964 | >.999 | >.999 | .016 | .788 | .996 | >.999 | >.999 | −.031 | .513 | .036 | .612 | — |
| lh | LO3 | .053 | .089 | .964 | .992 | >.999 | .084 | .104 | .996 | >.999 | >.999 | −.002 | .973 | .085 | .173 | — |
| lh | VMV2 | −.038 | .881 | .985 | >.999 | >.999 | −.033 | .434 | .996 | >.999 | >.999 | −.028 | .425 | −.015 | .765 | — |
| lh | 31pd | .020 | .280 | .964 | >.999 | >.999 | .003 | .955 | .996 | >.999 | >.999 | .032 | .412 | −.018 | .748 | — |
| lh | 31a | −.039 | .795 | .964 | >.999 | >.999 | −.128 | **.044** | .996 | .998 | .998 | .076 | .125 | −.178 | **.019** | AK− |
| lh | VVC | −.004 | .516 | .964 | >.999 | >.999 | .039 | .591 | .996 | >.999 | >.999 | −.050 | .368 | .071 | .407 | — |
| lh | 25 | .044 | .117 | .964 | .998 | >.999 | .012 | .805 | .996 | >.999 | >.999 | .062 | .119 | −.028 | .630 | — |
| lh | s32 | −.041 | .960 | .988 | >.999 | >.999 | −.049 | .098 | .996 | >.999 | >.999 | −.016 | .557 | −.038 | .288 | — |
| lh | pOFC | −.031 | .743 | .964 | >.999 | >.999 | −.014 | .822 | .996 | >.999 | >.999 | −.037 | .457 | .010 | .898 | — |
| lh | PoI1 | −.017 | .650 | .964 | >.999 | >.999 | .001 | .981 | .996 | >.999 | >.999 | −.032 | .490 | .022 | .756 | — |
| lh | Ig | −.012 | .631 | .964 | >.999 | >.999 | .018 | .708 | .996 | >.999 | >.999 | −.042 | .293 | .045 | .440 | — |
| lh | FOP5 | .013 | .392 | .964 | >.999 | >.999 | −.008 | .900 | .996 | >.999 | >.999 | .032 | .542 | −.029 | .714 | — |
| lh | p10p | −.023 | .709 | .964 | >.999 | >.999 | −.029 | .602 | .996 | >.999 | >.999 | −.007 | .867 | −.024 | .714 | — |
| lh | p47r | −.056 | .849 | .973 | >.999 | >.999 | −.064 | .400 | .996 | >.999 | >.999 | −.026 | .645 | −.047 | .601 | — |
| lh | TGv | −.030 | .705 | .964 | >.999 | >.999 | .008 | .920 | .996 | >.999 | >.999 | −.060 | .287 | .046 | .596 | — |
| lh | MBelt | .031 | .214 | .964 | >.999 | >.999 | .002 | .965 | .996 | >.999 | >.999 | .051 | .242 | −.031 | .638 | — |
| lh | LBelt | −.056 | .954 | .988 | >.999 | >.999 | −.075 | .094 | .996 | >.999 | >.999 | −.013 | .724 | −.067 | .218 | — |
| lh | A4 | .044 | .237 | .964 | .998 | >.999 | .097 | .259 | .996 | >.999 | >.999 | −.032 | .620 | .118 | .242 | — |
| lh | STSva | −.090 | .983 | .991 | >.999 | >.999 | −.002 | .968 | .996 | >.999 | >.999 | −.152 | **<.001** | .096 | .169 | Matching |
| lh | TE1m | −.031 | .715 | .964 | >.999 | >.999 | −.001 | .992 | .996 | >.999 | >.999 | −.053 | .333 | .033 | .698 | — |
| lh | PI | .028 | .260 | .964 | >.999 | >.999 | .035 | .555 | .996 | >.999 | >.999 | .009 | .850 | .029 | .674 | — |
| lh | a32pr | .040 | .210 | .964 | .999 | >.999 | .053 | .436 | .996 | >.999 | >.999 | .009 | .858 | .047 | .555 | — |
| lh | p24 | .024 | .299 | .964 | >.999 | >.999 | −.006 | .917 | .996 | >.999 | >.999 | .048 | .337 | −.037 | .620 | — |
| rh | V1 | −.004 | .519 | .964 | >.999 | >.999 | .041 | .598 | .996 | >.999 | >.999 | −.052 | .382 | .075 | .418 | — |
| rh | MST | −.000 | .507 | .964 | >.999 | >.999 | .006 | .907 | .996 | >.999 | >.999 | −.008 | .857 | .011 | .863 | — |
| rh | V6 | .005 | .449 | .964 | >.999 | >.999 | .052 | .334 | .996 | >.999 | >.999 | −.049 | .259 | .084 | .194 | — |
| rh | V2 | −.000 | .492 | .964 | >.999 | >.999 | .038 | .636 | .996 | >.999 | >.999 | −.043 | .480 | .066 | .486 | — |
| rh | V3 | .018 | .364 | .964 | >.999 | >.999 | .063 | .417 | .996 | >.999 | >.999 | −.039 | .509 | .088 | .338 | — |
| rh | V4 | .015 | .383 | .964 | >.999 | >.999 | .052 | .494 | .996 | >.999 | >.999 | −.032 | .584 | .073 | .420 | — |
| rh | V8 | −.088 | .982 | .991 | >.999 | >.999 | −.079 | .178 | .996 | >.999 | >.999 | −.063 | .178 | −.039 | .580 | — |
| rh | 4 | .042 | .227 | .964 | .998 | >.999 | .055 | .482 | .996 | >.999 | >.999 | .012 | .848 | .048 | .607 | — |
| rh | 3b | .017 | .363 | .964 | >.999 | >.999 | .016 | .827 | .996 | >.999 | >.999 | .011 | .833 | .009 | .918 | — |
| rh | FEF | −.038 | .784 | .964 | >.999 | >.999 | .016 | .805 | .996 | >.999 | >.999 | −.084 | .106 | .070 | .367 | — |
| rh | PEF | −.053 | .895 | .986 | >.999 | >.999 | .024 | .679 | .996 | >.999 | >.999 | −.119 | **.008** | .101 | .142 | Matching |
| rh | 55b | −.036 | .751 | .964 | >.999 | >.999 | .005 | .943 | .996 | >.999 | >.999 | −.067 | .220 | .048 | .570 | — |
| rh | V3A | −.010 | .585 | .964 | >.999 | >.999 | −.008 | .898 | .996 | >.999 | >.999 | −.008 | .862 | −.002 | .972 | — |
| rh | RSC | −.016 | .646 | .964 | >.999 | >.999 | −.080 | .113 | .996 | >.999 | >.999 | .062 | .132 | −.121 | **.044** | AK− |
| rh | POS2 | −.006 | .527 | .964 | >.999 | >.999 | −.012 | .873 | .996 | >.999 | >.999 | .004 | .946 | −.015 | .870 | — |
| rh | V7 | −.005 | .538 | .964 | >.999 | >.999 | .025 | .650 | .996 | >.999 | >.999 | −.036 | .420 | .048 | .465 | — |
| rh | IPS1 | .010 | .411 | .964 | >.999 | >.999 | .010 | .847 | .996 | >.999 | >.999 | .005 | .914 | .007 | .914 | — |
| rh | FFC | −.006 | .530 | .964 | >.999 | >.999 | .030 | .698 | .996 | >.999 | >.999 | −.043 | .463 | .058 | .513 | — |
| rh | V3B | −.050 | .939 | .988 | >.999 | >.999 | −.006 | .887 | .996 | >.999 | >.999 | −.079 | **.031** | .045 | .389 | Matching |
| rh | LO1 | .077 | **.022** | .964 | .912 | >.999 | .058 | .259 | .996 | >.999 | >.999 | .068 | .102 | .014 | .814 | — |
| rh | LO2 | −.015 | .652 | .964 | >.999 | >.999 | −.023 | .662 | .996 | >.999 | >.999 | −.000 | .997 | −.023 | .713 | — |
| rh | PIT | .019 | .349 | .964 | >.999 | >.999 | .043 | .528 | .996 | >.999 | >.999 | −.016 | .765 | .053 | .512 | — |
| rh | MT | −.014 | .635 | .964 | >.999 | >.999 | .025 | .640 | .996 | >.999 | >.999 | −.052 | .223 | .059 | .353 | — |
| rh | A1 | .062 | .058 | .964 | .977 | >.999 | .012 | .824 | .996 | >.999 | >.999 | .094 | **.026** | −.048 | .450 | Matching |
| rh | PSL | −.044 | .789 | .964 | >.999 | >.999 | −.005 | .941 | .996 | >.999 | >.999 | −.070 | .217 | .040 | .656 | — |
| rh | SFL | .007 | .440 | .964 | >.999 | >.999 | .039 | .595 | .996 | >.999 | >.999 | −.031 | .584 | .060 | .500 | — |
| rh | PCV | −.094 | .979 | .991 | >.999 | >.999 | −.148 | **.021** | .996 | .941 | .945 | .003 | .949 | −.151 | .050 | — |
| rh | STV | −.025 | .651 | .964 | >.999 | >.999 | .018 | .826 | .996 | >.999 | >.999 | −.062 | .326 | .058 | .557 | — |
| rh | 7Pm | −.016 | .644 | .964 | >.999 | >.999 | −.034 | .539 | .996 | >.999 | >.999 | .010 | .814 | −.041 | .532 | — |
| rh | 7m | −.074 | .949 | .988 | >.999 | >.999 | −.108 | .090 | .996 | >.999 | >.999 | −.007 | .885 | −.104 | .171 | — |
| rh | POS1 | −.027 | .701 | .964 | >.999 | >.999 | −.035 | .593 | .996 | >.999 | >.999 | −.007 | .892 | −.030 | .696 | — |
| rh | 23d | .012 | .360 | .964 | >.999 | >.999 | .039 | .432 | .996 | >.999 | >.999 | −.023 | .575 | .054 | .362 | — |
| rh | v23ab | .031 | .172 | .964 | >.999 | >.999 | .036 | .402 | .996 | >.999 | >.999 | .014 | .709 | .027 | .597 | — |
| rh | d23ab | .007 | .430 | .964 | >.999 | >.999 | −.026 | .665 | .996 | >.999 | >.999 | .041 | .377 | −.053 | .458 | — |
| rh | 31pv | −.015 | .640 | .964 | >.999 | >.999 | −.002 | .975 | .996 | >.999 | >.999 | −.024 | .594 | .014 | .835 | — |
| rh | 5m | −.016 | .640 | .964 | >.999 | >.999 | −.038 | .513 | .996 | >.999 | >.999 | .015 | .737 | −.048 | .485 | — |
| rh | 5mv | −.007 | .560 | .964 | >.999 | >.999 | .041 | .509 | .996 | >.999 | >.999 | −.059 | .234 | .080 | .290 | — |
| rh | 23c | −.008 | .575 | .964 | >.999 | >.999 | −.015 | .797 | .996 | >.999 | >.999 | .003 | .953 | −.017 | .812 | — |
| rh | 5L | −.006 | .549 | .964 | >.999 | >.999 | −.039 | .486 | .996 | >.999 | >.999 | .033 | .459 | −.060 | .369 | — |
| rh | 24dd | .035 | .222 | .964 | >.999 | >.999 | .041 | .506 | .996 | >.999 | >.999 | .014 | .780 | .032 | .651 | — |
| rh | 24dv | .019 | .321 | .964 | >.999 | >.999 | .055 | .331 | .996 | >.999 | >.999 | −.029 | .527 | .073 | .275 | — |
| rh | 7AL | −.004 | .533 | .964 | >.999 | >.999 | −.018 | .754 | .996 | >.999 | >.999 | .013 | .771 | −.027 | .692 | — |
| rh | SCEF | −.010 | .584 | .964 | >.999 | >.999 | .020 | .750 | .996 | >.999 | >.999 | −.041 | .419 | .047 | .540 | — |
| rh | 6ma | −.033 | .720 | .964 | >.999 | >.999 | .034 | .652 | .996 | >.999 | >.999 | −.095 | .102 | .096 | .284 | — |
| rh | 7Am | −.010 | .582 | .964 | >.999 | >.999 | −.026 | .686 | .996 | >.999 | >.999 | .012 | .819 | −.034 | .658 | — |
| rh | 7Pl | .005 | .446 | .964 | >.999 | >.999 | −.001 | .984 | .996 | >.999 | >.999 | .010 | .819 | −.007 | .909 | — |
| rh | 7PC | −.072 | .940 | .988 | >.999 | >.999 | −.062 | .331 | .996 | >.999 | >.999 | −.055 | .271 | −.026 | .728 | — |
| rh | LIPv | −.025 | .737 | .964 | >.999 | >.999 | .005 | .917 | .996 | >.999 | >.999 | −.048 | .243 | .036 | .545 | — |
| rh | VIP | −.032 | .789 | .964 | >.999 | >.999 | −.037 | .496 | .996 | >.999 | >.999 | −.014 | .751 | −.028 | .663 | — |
| rh | MIP | −.007 | .547 | .964 | >.999 | >.999 | .002 | .974 | .996 | >.999 | >.999 | −.015 | .784 | .012 | .885 | — |
| rh | 1 | .051 | .173 | .964 | .995 | >.999 | .076 | .307 | .996 | >.999 | >.999 | .003 | .960 | .075 | .398 | — |
| rh | 2 | −.096 | .973 | .989 | >.999 | >.999 | −.121 | .081 | .996 | >.999 | >.999 | −.030 | .579 | −.101 | .222 | — |
| rh | 3a | −.034 | .806 | .964 | >.999 | >.999 | −.009 | .861 | .996 | >.999 | >.999 | −.049 | .259 | .022 | .724 | — |
| rh | 6d | .031 | .255 | .964 | >.999 | >.999 | .056 | .386 | .996 | >.999 | >.999 | −.010 | .852 | .062 | .422 | — |
| rh | 6mp | .001 | .489 | .964 | >.999 | >.999 | .024 | .722 | .996 | >.999 | >.999 | −.025 | .625 | .040 | .613 | — |
| rh | 6v | .008 | .433 | .964 | >.999 | >.999 | .049 | .512 | .996 | >.999 | >.999 | −.042 | .464 | .076 | .388 | — |
| rh | p24pr | .052 | .091 | .964 | .994 | >.999 | .073 | .162 | .996 | >.999 | >.999 | .007 | .859 | .069 | .272 | — |
| rh | 33pr | .004 | .428 | .964 | >.999 | >.999 | .003 | .915 | .996 | >.999 | >.999 | .004 | .906 | .001 | .977 | — |
| rh | a24pr | −.040 | .843 | .973 | >.999 | >.999 | −.071 | .188 | .996 | >.999 | >.999 | .010 | .813 | −.077 | .230 | — |
| rh | p32pr | −.005 | .540 | .964 | >.999 | >.999 | .021 | .731 | .996 | >.999 | >.999 | −.032 | .512 | .042 | .566 | — |
| rh | a24 | .075 | **.041** | .964 | .920 | >.999 | .082 | .163 | .996 | >.999 | >.999 | .038 | .411 | .057 | .420 | — |
| rh | d32 | .046 | .164 | .964 | .997 | >.999 | −.007 | .920 | .996 | >.999 | >.999 | .088 | .094 | −.064 | .425 | — |
| rh | 8BM | −.013 | .584 | .964 | >.999 | >.999 | −.013 | .870 | .996 | >.999 | >.999 | −.008 | .889 | −.007 | .937 | — |
| rh | p32 | −.036 | .822 | .964 | >.999 | >.999 | −.054 | .298 | .996 | >.999 | >.999 | −.002 | .968 | −.052 | .397 | — |
| rh | 10r | −.013 | .618 | .964 | >.999 | >.999 | .021 | .693 | .996 | >.999 | >.999 | −.046 | .304 | .051 | .442 | — |
| rh | 47m | .006 | .429 | .964 | >.999 | >.999 | .038 | .475 | .996 | >.999 | >.999 | −.031 | .460 | .058 | .355 | — |
| rh | 8Av | −.023 | .632 | .964 | >.999 | >.999 | −.054 | .547 | .996 | >.999 | >.999 | .020 | .765 | −.067 | .523 | — |
| rh | 8Ad | .032 | .265 | .964 | >.999 | >.999 | .065 | .367 | .996 | >.999 | >.999 | −.018 | .738 | .077 | .365 | — |
| rh | 9m | −.032 | .692 | .964 | >.999 | >.999 | .013 | .878 | .996 | >.999 | >.999 | −.070 | .272 | .058 | .558 | — |
| rh | 8BL | −.027 | .653 | .964 | >.999 | >.999 | .007 | .938 | .996 | >.999 | >.999 | −.054 | .415 | .041 | .688 | — |
| rh | 9p | .009 | .418 | .964 | >.999 | >.999 | .006 | .926 | .996 | >.999 | >.999 | .009 | .857 | .000 | .998 | — |
| rh | 10d | −.003 | .518 | .964 | >.999 | >.999 | .010 | .868 | .996 | >.999 | >.999 | −.017 | .710 | .021 | .765 | — |
| rh | 8C | −.008 | .539 | .964 | >.999 | >.999 | −.057 | .523 | .996 | >.999 | >.999 | .049 | .472 | −.088 | .398 | — |
| rh | 44ᵃ | .006 | .458 | .964 | >.999 | >.999 | .037 | .659 | .996 | >.999 | >.999 | −.032 | .622 | .058 | .564 | — |
| rh | 45 | −.020 | .626 | .964 | >.999 | >.999 | .041 | .612 | .996 | >.999 | >.999 | −.080 | .185 | .093 | .330 | — |
| rh | 47l | .017 | .355 | .964 | >.999 | >.999 | .095 | .142 | .996 | >.999 | >.999 | −.077 | .125 | .145 | .061 | — |
| rh | a47r | .001 | .474 | .964 | >.999 | >.999 | .012 | .847 | .996 | >.999 | >.999 | −.011 | .824 | .020 | .803 | — |
| rh | 6r | −.015 | .596 | .964 | >.999 | >.999 | .001 | .986 | .996 | >.999 | >.999 | −.027 | .660 | .019 | .842 | — |
| rh | IFJa | .032 | .213 | .964 | >.999 | >.999 | .036 | .509 | .996 | >.999 | >.999 | .015 | .736 | .027 | .683 | — |
| rh | IFJp | .040 | .100 | .964 | .999 | >.999 | .041 | .307 | .996 | >.999 | >.999 | .023 | .510 | .026 | .578 | — |
| rh | IFSp | −.039 | .759 | .964 | >.999 | >.999 | .061 | .418 | .996 | >.999 | >.999 | −.135 | **.018** | .149 | .094 | Matching |
| rh | IFSa | −.101 | .961 | .988 | >.999 | >.999 | −.090 | .269 | .996 | >.999 | >.999 | −.073 | .238 | −.042 | .654 | — |
| rh | p9-46v | −.022 | .623 | .964 | >.999 | >.999 | −.060 | .506 | .996 | >.999 | >.999 | .029 | .670 | −.078 | .462 | — |
| rh | 46 | −.002 | .502 | .964 | >.999 | >.999 | .007 | .931 | .996 | >.999 | >.999 | −.011 | .846 | .014 | .877 | — |
| rh | a9-46v | −.021 | .643 | .964 | >.999 | >.999 | −.021 | .773 | .996 | >.999 | >.999 | −.012 | .827 | −.013 | .880 | — |
| rh | 9-46d | −.039 | .754 | .964 | >.999 | >.999 | −.061 | .421 | .996 | >.999 | >.999 | .001 | .986 | −.061 | .487 | — |
| rh | 9a | .023 | .326 | .964 | >.999 | >.999 | .063 | .394 | .996 | >.999 | >.999 | −.030 | .595 | .082 | .346 | — |
| rh | 10v | .031 | .204 | .964 | >.999 | >.999 | .096 | .067 | .996 | >.999 | >.999 | −.052 | .213 | .129 | **.036** | AK+ |
| rh | a10p | .019 | .314 | .964 | >.999 | >.999 | .068 | .223 | .996 | >.999 | >.999 | −.042 | .336 | .095 | .149 | — |
| rh | 10pp | .005 | .433 | .964 | >.999 | >.999 | .047 | .381 | .996 | >.999 | >.999 | −.043 | .318 | .074 | .243 | — |
| rh | 11l | −.004 | .522 | .964 | >.999 | >.999 | .064 | .329 | .996 | >.999 | >.999 | −.078 | .124 | .115 | .139 | — |
| rh | 13l | .029 | .235 | .964 | >.999 | >.999 | .096 | .080 | .996 | >.999 | >.999 | −.056 | .194 | .133 | **.040** | AK+ |
| rh | OFC | .007 | .433 | .964 | >.999 | >.999 | .043 | .472 | .996 | >.999 | >.999 | −.037 | .439 | .067 | .348 | — |
| rh | 47s | .051 | .134 | .964 | .994 | >.999 | .081 | .199 | .996 | >.999 | >.999 | −.002 | .973 | .082 | .275 | — |
| rh | LIPd | −.014 | .690 | .964 | >.999 | >.999 | −.000 | .992 | .996 | >.999 | >.999 | −.024 | .456 | .015 | .738 | — |
| rh | 6a | −.076 | .944 | .988 | >.999 | >.999 | −.053 | .434 | .996 | >.999 | >.999 | −.073 | .165 | −.006 | .945 | — |
| rh | i6-8 | −.024 | .675 | .964 | >.999 | >.999 | −.069 | .325 | .996 | >.999 | >.999 | .036 | .515 | −.092 | .267 | — |
| rh | s6-8 | −.000 | .494 | .964 | >.999 | >.999 | .077 | .239 | .996 | >.999 | >.999 | −.086 | .087 | .133 | .086 | — |
| rh | 43 | .065 | .084 | .964 | .968 | >.999 | .161 | **.012** | .996 | .830 | .835 | −.067 | .179 | .204 | **.007** | AK+ |
| rh | OP4 | −.011 | .588 | .964 | >.999 | >.999 | .036 | .567 | .996 | >.999 | >.999 | −.058 | .235 | .073 | .320 | — |
| rh | OP1 | −.016 | .655 | .964 | >.999 | >.999 | .008 | .881 | .996 | >.999 | >.999 | −.037 | .400 | .032 | .629 | — |
| rh | OP2-3 | .034 | .158 | .964 | >.999 | >.999 | .024 | .595 | .996 | >.999 | >.999 | .032 | .403 | .004 | .944 | — |
| rh | 52 | −.024 | .774 | .964 | >.999 | >.999 | −.023 | .574 | .996 | >.999 | >.999 | −.016 | .651 | −.012 | .799 | — |
| rh | RI | −.053 | .967 | .989 | >.999 | >.999 | −.025 | .513 | .996 | >.999 | >.999 | −.063 | .051 | .016 | .719 | — |
| rh | PFcm | −.065 | .938 | .988 | >.999 | >.999 | .021 | .730 | .996 | >.999 | >.999 | −.134 | **.004** | .107 | .121 | Matching |
| rh | PoI2 | .076 | .056 | .964 | .918 | >.999 | .118 | .069 | .996 | >.999 | >.999 | −.001 | .987 | .119 | .124 | — |
| rh | TA2 | .025 | .288 | .964 | >.999 | >.999 | .045 | .470 | .996 | >.999 | >.999 | −.007 | .881 | .049 | .504 | — |
| rh | FOP4 | −.043 | .845 | .973 | >.999 | >.999 | −.042 | .465 | .996 | >.999 | >.999 | −.027 | .539 | −.024 | .723 | — |
| rh | MI | .019 | .342 | .964 | >.999 | >.999 | .018 | .791 | .996 | >.999 | >.999 | .012 | .810 | .010 | .904 | — |
| rh | Pir | .028 | .259 | .964 | >.999 | >.999 | .068 | .258 | .996 | >.999 | >.999 | −.028 | .551 | .086 | .226 | — |
| rh | AVI | .007 | .428 | .964 | >.999 | >.999 | .059 | .276 | .996 | >.999 | >.999 | −.054 | .223 | .093 | .145 | — |
| rh | AAIC | .018 | .330 | .964 | >.999 | >.999 | .065 | .270 | .996 | >.999 | >.999 | −.042 | .373 | .093 | .187 | — |
| rh | FOP1 | −.024 | .724 | .964 | >.999 | >.999 | −.028 | .596 | .996 | >.999 | >.999 | −.010 | .812 | −.021 | .741 | — |
| rh | FOP3 | −.020 | .747 | .964 | >.999 | >.999 | .023 | .560 | .996 | >.999 | >.999 | −.060 | .076 | .062 | .194 | — |
| rh | FOP2 | −.041 | .893 | .986 | >.999 | >.999 | −.021 | .633 | .996 | >.999 | >.999 | −.046 | .216 | .009 | .872 | — |
| rh | PFt | −.039 | .808 | .964 | >.999 | >.999 | −.031 | .618 | .996 | >.999 | >.999 | −.033 | .490 | −.009 | .904 | — |
| rh | AIP | −.065 | .922 | .988 | >.999 | >.999 | −.069 | .271 | .996 | >.999 | >.999 | −.034 | .491 | −.047 | .531 | — |
| rh | EC | .078 | .064 | .964 | .905 | >.999 | .113 | .103 | .996 | >.999 | >.999 | .009 | .870 | .107 | .197 | — |
| rh | PreS | .027 | .245 | .964 | >.999 | >.999 | −.041 | .459 | .996 | >.999 | >.999 | .092 | **.035** | −.100 | .123 | Matching |
| rh | H | −.004 | .530 | .964 | >.999 | >.999 | −.013 | .816 | .996 | >.999 | >.999 | .008 | .859 | −.018 | .790 | — |
| rh | ProS | −.034 | .821 | .964 | >.999 | >.999 | .009 | .852 | .996 | >.999 | >.999 | −.069 | .085 | .054 | .358 | — |
| rh | PeEc | −.005 | .530 | .964 | >.999 | >.999 | .034 | .651 | .996 | >.999 | >.999 | −.046 | .416 | .064 | .468 | — |
| rh | STGa | .073 | .061 | .964 | .936 | >.999 | .147 | **.015** | .996 | .950 | .953 | −.039 | .421 | .172 | **.018** | AK+ |
| rh | PBelt | −.009 | .585 | .964 | >.999 | >.999 | .048 | .401 | .996 | >.999 | >.999 | −.069 | .126 | .092 | .175 | — |
| rh | A5 | .009 | .430 | .964 | >.999 | >.999 | .032 | .715 | .996 | >.999 | >.999 | −.019 | .757 | .044 | .666 | — |
| rh | PHA1 | .087 | **.038** | .964 | .833 | >.999 | .092 | .170 | .996 | >.999 | >.999 | .047 | .375 | .062 | .439 | — |
| rh | PHA3 | .022 | .253 | .964 | >.999 | >.999 | .045 | .284 | .996 | >.999 | >.999 | −.014 | .704 | .054 | .288 | — |
| rh | STSda | −.005 | .531 | .964 | >.999 | >.999 | .021 | .774 | .996 | >.999 | >.999 | −.032 | .572 | .041 | .633 | — |
| rh | STSdp | −.136 | .995 | .998 | >.999 | >.999 | −.112 | .128 | .996 | >.999 | >.999 | −.110 | **.050** | −.040 | .643 | Matching |
| rh | STSvp | −.010 | .559 | .964 | >.999 | >.999 | −.038 | .651 | .996 | >.999 | >.999 | .025 | .695 | −.054 | .589 | — |
| rh | TGd | .023 | .340 | .964 | >.999 | >.999 | .120 | .142 | .996 | >.999 | >.999 | −.094 | .130 | .181 | .060 | — |
| rh | TE1a | −.023 | .653 | .964 | >.999 | >.999 | .015 | .847 | .996 | >.999 | >.999 | −.056 | .331 | .051 | .566 | — |
| rh | TE1p | −.013 | .580 | .964 | >.999 | >.999 | −.016 | .849 | .996 | >.999 | >.999 | −.006 | .926 | −.012 | .898 | — |
| rh | TE2a | −.044 | .778 | .964 | >.999 | >.999 | −.001 | .994 | .996 | >.999 | >.999 | −.075 | .202 | .048 | .599 | — |
| rh | TF | .001 | .487 | .964 | >.999 | >.999 | .002 | .978 | .996 | >.999 | >.999 | −.001 | .984 | .003 | .975 | — |
| rh | TE2p | −.040 | .781 | .964 | >.999 | >.999 | −.026 | .710 | .996 | >.999 | >.999 | −.039 | .461 | −.000 | .996 | — |
| rh | PHT | .002 | .480 | .964 | >.999 | >.999 | .021 | .807 | .996 | >.999 | >.999 | −.019 | .758 | .033 | .736 | — |
| rh | PH | −.018 | .652 | .964 | >.999 | >.999 | −.011 | .856 | .996 | >.999 | >.999 | −.019 | .693 | .001 | .987 | — |
| rh | TPOJ1 | −.015 | .585 | .964 | >.999 | >.999 | −.015 | .862 | .996 | >.999 | >.999 | −.008 | .900 | −.009 | .932 | — |
| rh | TPOJ2 | .025 | .325 | .964 | >.999 | >.999 | .025 | .753 | .996 | >.999 | >.999 | .015 | .798 | .015 | .870 | — |
| rh | TPOJ3 | .002 | .475 | .964 | >.999 | >.999 | .050 | .355 | .996 | >.999 | >.999 | −.052 | .227 | .083 | .195 | — |
| rh | DVT | −.017 | .628 | .964 | >.999 | >.999 | −.041 | .541 | .996 | >.999 | >.999 | .016 | .753 | −.052 | .518 | — |
| rh | PGp | .035 | .232 | .964 | >.999 | >.999 | .020 | .760 | .996 | >.999 | >.999 | .038 | .477 | −.004 | .960 | — |
| rh | IP2 | −.070 | .930 | .988 | >.999 | >.999 | −.051 | .438 | .996 | >.999 | >.999 | −.064 | .209 | −.010 | .895 | — |
| rh | IP1 | −.014 | .615 | .964 | >.999 | >.999 | −.022 | .727 | .996 | >.999 | >.999 | −.000 | .994 | −.021 | .767 | — |
| rh | IP0 | −.006 | .546 | .964 | >.999 | >.999 | −.013 | .834 | .996 | >.999 | >.999 | .004 | .938 | −.015 | .836 | — |
| rh | PFop | −.045 | .845 | .973 | >.999 | >.999 | .000 | >.999 | >.999 | >.999 | >.999 | −.077 | .101 | .050 | .490 | — |
| rh | PF | −.024 | .661 | .964 | >.999 | >.999 | −.017 | .824 | .996 | >.999 | >.999 | −.022 | .701 | −.003 | .973 | — |
| rh | PFm | −.061 | .849 | .973 | >.999 | >.999 | −.090 | .268 | .996 | >.999 | >.999 | −.004 | .947 | −.087 | .370 | — |
| rh | PGi | .016 | .385 | .964 | >.999 | >.999 | −.028 | .738 | .996 | >.999 | >.999 | .058 | .354 | −.065 | .506 | — |
| rh | PGs | −.031 | .712 | .964 | >.999 | >.999 | −.118 | .113 | .996 | >.999 | >.999 | .077 | .177 | −.167 | .056 | — |
| rh | V6A | .001 | .485 | .964 | >.999 | >.999 | .007 | .884 | .996 | >.999 | >.999 | −.006 | .877 | .011 | .846 | — |
| rh | VMV1 | .019 | .327 | .964 | >.999 | >.999 | .042 | .496 | .996 | >.999 | >.999 | −.013 | .782 | .050 | .495 | — |
| rh | VMV3 | .015 | .331 | .964 | >.999 | >.999 | .043 | .390 | .996 | >.999 | >.999 | −.021 | .597 | .057 | .340 | — |
| rh | PHA2 | −.036 | .854 | .976 | >.999 | >.999 | −.030 | .505 | .996 | >.999 | >.999 | −.028 | .454 | −.012 | .824 | — |
| rh | V4t | −.045 | .878 | .985 | >.999 | >.999 | .007 | .895 | .996 | >.999 | >.999 | −.085 | **.044** | .062 | .327 | Matching |
| rh | FST | −.028 | .743 | .964 | >.999 | >.999 | −.084 | .153 | .996 | >.999 | >.999 | .045 | .333 | −.113 | .104 | — |
| rh | V3CD | −.015 | .642 | .964 | >.999 | >.999 | −.012 | .824 | .996 | >.999 | >.999 | −.013 | .764 | −.004 | .950 | — |
| rh | LO3 | −.050 | .872 | .983 | >.999 | >.999 | −.048 | .414 | .996 | >.999 | >.999 | −.032 | .503 | −.027 | .695 | — |
| rh | VMV2 | .029 | .187 | .964 | >.999 | >.999 | .022 | .614 | .996 | >.999 | >.999 | .026 | .472 | .005 | .923 | — |
| rh | 31pd | .067 | .053 | .964 | .963 | >.999 | .016 | .773 | .996 | >.999 | >.999 | .096 | **.028** | −.046 | .485 | Matching |
| rh | 31a | −.027 | .721 | .964 | >.999 | >.999 | −.047 | .437 | .996 | >.999 | >.999 | .006 | .893 | −.051 | .477 | — |
| rh | VVC | .004 | .458 | .964 | >.999 | >.999 | .029 | .676 | .996 | >.999 | >.999 | −.025 | .629 | .045 | .572 | — |
| rh | 25 | .070 | **.021** | .964 | .950 | >.999 | .068 | .132 | .996 | >.999 | >.999 | .044 | .240 | .040 | .465 | — |
| rh | s32 | .005 | .425 | .964 | >.999 | >.999 | .038 | .368 | .996 | >.999 | >.999 | −.033 | .345 | .059 | .237 | — |
| rh | pOFC | −.011 | .589 | .964 | >.999 | >.999 | .072 | .211 | .996 | >.999 | >.999 | −.099 | **.030** | .135 | **.045** | Match + AK+ |
| rh | PoI1 | .027 | .240 | .964 | >.999 | >.999 | .025 | .633 | .996 | >.999 | >.999 | .020 | .643 | .012 | .849 | — |
| rh | Ig | .019 | .275 | .964 | >.999 | >.999 | .004 | .919 | .996 | >.999 | >.999 | .028 | .431 | −.014 | .776 | — |
| rh | FOP5 | .017 | .334 | .964 | >.999 | >.999 | .049 | .390 | .996 | >.999 | >.999 | −.025 | .586 | .065 | .332 | — |
| rh | p10p | .027 | .266 | .964 | >.999 | >.999 | .013 | .828 | .996 | >.999 | >.999 | .032 | .502 | −.008 | .913 | — |
| rh | p47r | −.017 | .623 | .964 | >.999 | >.999 | .002 | .974 | .996 | >.999 | >.999 | −.032 | .535 | .022 | .771 | — |
| rh | TGv | −.038 | .768 | .964 | >.999 | >.999 | −.010 | .883 | .996 | >.999 | >.999 | −.054 | .313 | .025 | .764 | — |
| rh | MBelt | −.008 | .581 | .964 | >.999 | >.999 | .035 | .481 | .996 | >.999 | >.999 | −.052 | .194 | .069 | .244 | — |
| rh | LBelt | −.027 | .755 | .964 | >.999 | >.999 | .001 | .978 | .996 | >.999 | >.999 | −.049 | .256 | .033 | .606 | — |
| rh | A4 | .100 | **.034** | .964 | .685 | >.999 | .178 | **.015** | .996 | .616 | .621 | −.026 | .648 | .195 | **.026** | AK+ |
| rh | STSva | .007 | .447 | .964 | >.999 | >.999 | .056 | .445 | .996 | >.999 | >.999 | −.050 | .375 | .088 | .309 | — |
| rh | TE1m | .067 | .113 | .964 | .961 | >.999 | .138 | .069 | .996 | .986 | .987 | −.038 | .518 | .162 | .070 | — |
| rh | PI | .008 | .416 | .964 | >.999 | >.999 | .003 | .958 | .996 | >.999 | >.999 | .011 | .820 | −.004 | .957 | — |
| rh | a32pr | −.049 | .865 | .979 | >.999 | >.999 | −.064 | .285 | .996 | >.999 | >.999 | −.012 | .789 | −.056 | .431 | — |
| rh | p24 | .044 | .168 | .964 | .998 | >.999 | .030 | .635 | .996 | >.999 | >.999 | .042 | .384 | .003 | .971 | — |

Note. All 360 parcels with complete data are listed for each model; no selection was applied, so every parcel can be looked up under every correction. Inter-subject dissimilarity was computed as 1 − Spearman correlation between participants' parcel-wise representational dissimilarity matrices, rank-transformed and residualised on the covariate inter-subject dissimilarity matrices before analysis; all tests are permutation tests on participant labels with 20,000 permutations. The matching statistic is the rank correlation between neural dissimilarity and |a − b| in early life stress, tested one-tailed positive: it asks whether participants with similar exposure have similar representational geometry. The additive (Anna Karenina) statistic is the rank correlation with mean(a, b), tested two-tailed, because both directions are substantive hypotheses. p is the uncorrected permutation p value; q is the Benjamini–Hochberg false discovery rate across parcels; p FWE-P is corrected by the maximum statistic across parcels within a statistic; p FWE-F is corrected across parcels and both confirmatory statistics jointly, using shared permutations, and is the inferential criterion for the confirmatory family.

β coefficients are partial standardised coefficients from the joint model neural dissimilarity ~ |a − b| + mean(a, b), with two-tailed permutation p values; because the two predictors are close to orthogonal after ranking and covariate residualisation (r = .31 to .65), they estimate the matching and additive components separately. The Mechanism column is derived from these coefficients alone: Matching denotes a reliable partial |a − b| term, AK+ and AK− a reliable partial mean(a, b) term of the stated sign, and an em dash a parcel where neither partial coefficient reached p < .05, including parcels where a marginal statistic did. AK+ denotes greater dissimilarity at high exposure (representational idiosyncrasy at high early life stress); AK− denotes greater similarity at high exposure (convergence at high exposure, idiosyncrasy at low). Values below .05 are set in bold throughout; bold is descriptive and does not itself denote a corrected result.

ᵃ A priori region of interest.
