## Supplementary Table S1 for "The Influence of Early Life Stress on the Development of Neural Representations during Mentalizing"

**Supplementary Table S1. Cluster-corrected univariate surface results.**

| **Direction** | **Hemi** | **Anatomical label (peak vertex)** | **Size (mm²)** | **Vertices** | **Peak MNI305 (x, y, z)** | **Peak t** | **Cluster-wise p** | **90% CI** |
| --- | --- | --- | --- | --- | --- | --- | --- | --- |
| **One-sample (group mean) — Mindreading > control (N = 89, df = 88)** | | | | | | | | |
| Negative | Left | inferiorparietal | 3290.1 | 7293 | -43, -63, 44 | -9.81 | .0001 | [< .0001, .0002] |
| Negative | Left | precuneus | 3270.2 | 6897 | -4, -66, 28 | -9.92 | .0001 | [< .0001, .0002] |
| Negative | Left | superiorfrontal | 1238.5 | 2263 | -20, 10, 58 | -7.28 | .0001 | [< .0001, .0002] |
| Negative | Left | rostralmiddlefrontal | 714.8 | 1014 | -36, 54, -2 | -6.60 | .0001 | [< .0001, .0002] |
| Negative | Left | rostralmiddlefrontal | 704.1 | 1132 | -36, 26, 37 | -7.15 | .0001 | [< .0001, .0002] |
| Negative | Left | posteriorcingulate | 509.6 | 1393 | -7, -30, 40 | -8.89 | .0001 | [< .0001, .0002] |
| Negative | Left | middletemporal | 485.0 | 776 | -62, -35, -16 | -6.01 | .0001 | [< .0001, .0002] |
| Negative | Left | rostralmiddlefrontal | 374.6 | 474 | -24, 57, 14 | -6.75 | .0001 | [< .0001, .0002] |
| Negative | Left | insula | 114.4 | 250 | -37, -6, -1 | -4.02 | .0104 | [.0091, .0117] |
| Negative | Left | precentral | 109.9 | 212 | -32, -14, 65 | -4.04 | .0130ᵃ | [.0116, .0145] |
| Negative | Left | caudalanteriorcingulate | 102.3 | 258 | -11, 7, 37 | -4.20 | .0185ᵃ | [.0168, .0202] |
| Positive | Left | bankssts | 15991.9 | 26420 | -50, -41, 4 | +13.77 | .0001 | [< .0001, .0002] |
| Positive | Left | parstriangularis | 3312.7 | 6441 | -52, 30, 7 | +13.82 | .0001 | [< .0001, .0002] |
| Positive | Left | superiorfrontal | 1626.6 | 2969 | -8, 58, 27 | +9.32 | .0001 | [< .0001, .0002] |
| Positive | Left | caudalmiddlefrontal | 830.3 | 1523 | -43, 3, 48 | +10.37 | .0001 | [< .0001, .0002] |
| Negative | Right | superiorfrontal | 5421.9 | 8848 | 21, 24, 53 | -10.96 | .0001 | [< .0001, .0002] |
| Negative | Right | supramarginal | 4223.2 | 8920 | 52, -40, 44 | -13.48 | .0001 | [< .0001, .0002] |
| Negative | Right | precuneus | 2047.1 | 4407 | 5, -59, 28 | -12.36 | .0001 | [< .0001, .0002] |
| Negative | Right | inferiortemporal | 1304.3 | 2039 | 55, -50, -12 | -8.68 | .0001 | [< .0001, .0002] |
| Negative | Right | posteriorcingulate | 678.2 | 1682 | 5, -31, 38 | -11.32 | .0001 | [< .0001, .0002] |
| Negative | Right | rostralanteriorcingulate | 369.3 | 735 | 9, 36, -5 | -4.73 | .0001 | [< .0001, .0002] |
| Negative | Right | superiorfrontal | 291.0 | 533 | 10, 38, 30 | -6.22 | .0001 | [< .0001, .0002] |
| Negative | Right | lateralorbitofrontal | 179.7 | 341 | 23, 35, -12 | -4.38 | .0007 | [.0004, .0010] |
| Negative | Right | rostralmiddlefrontal | 155.8 | 219 | 24, 46, 32 | -5.82 | .0015 | [.0010, .0020] |
| Negative | Right | parahippocampal | 113.5 | 226 | 32, -25, -23 | -5.24 | .0090 | [.0078, .0102] |
| Negative | Right | insula | 104.8 | 267 | 37, -2, 7 | -5.09 | .0140ᵃ | [.0125, .0155] |
| Positive | Right | superiortemporal | 13401.1 | 22489 | 50, -4, -17 | +13.00 | .0001 | [< .0001, .0002] |
| Positive | Right | parstriangularis | 1369.8 | 2517 | 54, 27, 8 | +11.72 | .0001 | [< .0001, .0002] |
| Positive | Right | postcentral | 663.2 | 1368 | 30, -30, 55 | +5.48 | .0001 | [< .0001, .0002] |
| Positive | Right | precentral | 373.9 | 625 | 44, 2, 46 | +8.02 | .0001 | [< .0001, .0002] |
| Positive | Right | superiorfrontal | 258.8 | 549 | 8, 12, 61 | +6.93 | .0001 | [< .0001, .0002] |
| Positive | Right | superiorfrontal | 206.7 | 371 | 8, 56, 31 | +6.13 | .0001 | [< .0001, .0002] |
| Positive | Right | paracentral | 201.0 | 416 | 6, -26, 64 | +4.30 | .0002 | [< .0001, .0004] |
| Positive | Right | supramarginal | 109.6 | 350 | 38, -22, 21 | +4.19 | .0131ᵃ | [.0117, .0146] |
| **One-sample (group mean) — Correct > incorrect (N = 89, df = 88)** | | | | | | | | |
| Negative | Left | lateraloccipital | 3212.2 | 4211 | -21, -98, 2 | -7.06 | .0001 | [< .0001, .0002] |
| Negative | Left | lateralorbitofrontal | 2596.0 | 5617 | -28, 23, -2 | -10.45 | .0001 | [< .0001, .0002] |
| Negative | Left | superiorfrontal | 1552.8 | 2798 | -9, 25, 37 | -10.19 | .0001 | [< .0001, .0002] |
| Negative | Left | rostralmiddlefrontal | 510.0 | 715 | -39, 40, -3 | -5.71 | .0001 | [< .0001, .0002] |
| Negative | Left | superiorparietal | 129.9 | 334 | -25, -57, 51 | -4.51 | .0052 | [.0043, .0061] |
| Positive | Left | precuneus | 2671.3 | 6085 | -4, -61, 20 | +6.48 | .0001 | [< .0001, .0002] |
| Positive | Left | inferiorparietal | 1756.0 | 3290 | -45, -73, 27 | +6.68 | .0001 | [< .0001, .0002] |
| Positive | Left | lingual | 577.2 | 676 | -11, -76, -7 | +5.78 | .0001 | [< .0001, .0002] |
| Positive | Left | rostralanteriorcingulate | 555.2 | 1013 | -8, 39, -5 | +6.05 | .0001 | [< .0001, .0002] |
| Positive | Left | superiortemporal | 552.3 | 957 | -49, -11, -17 | +5.05 | .0001 | [< .0001, .0002] |
| Positive | Left | superiortemporal | 487.0 | 1072 | -54, -4, -5 | +5.35 | .0001 | [< .0001, .0002] |
| Positive | Left | superiorfrontal | 388.6 | 688 | -22, 23, 48 | +4.66 | .0001 | [< .0001, .0002] |
| Positive | Left | cuneus | 258.8 | 304 | -4, -88, 16 | +4.51 | .0001 | [< .0001, .0002] |
| Positive | Left | supramarginal | 239.8 | 491 | -62, -32, 27 | +4.99 | .0002 | [< .0001, .0004] |
| Positive | Left | parahippocampal | 231.7 | 495 | -30, -39, -12 | +4.43 | .0002 | [< .0001, .0004] |
| Positive | Left | precentral | 166.0 | 404 | -54, -7, 10 | +5.26 | .0016 | [.0011, .0021] |
| Positive | Left | parsopercularis | 108.8 | 332 | -38, 6, 13 | +5.32 | .0130ᵃ | [.0116, .0145] |
| Positive | Left | superiortemporal | 106.9 | 229 | -63, -27, 6 | +3.88 | .0140ᵃ | [.0125, .0155] |
| Positive | Left | supramarginal | 100.3 | 227 | -46, -34, 22 | +4.71 | .0196ᵃ | [.0178, .0214] |
| Negative | Right | lateraloccipital | 5483.3 | 7457 | 26, -83, -10 | -7.92 | .0001 | [< .0001, .0002] |
| Negative | Right | insula | 2024.5 | 4161 | 35, 17, -4 | -10.46 | .0001 | [< .0001, .0002] |
| Negative | Right | superiorfrontal | 1118.9 | 2294 | 8, 24, 42 | -8.70 | .0001 | [< .0001, .0002] |
| Negative | Right | caudalmiddlefrontal | 143.6 | 296 | 33, 6, 30 | -4.48 | .0030 | [.0023, .0037] |
| Positive | Right | inferiorparietal | 2612.8 | 5448 | 44, -66, 26 | +5.57 | .0001 | [< .0001, .0002] |
| Positive | Right | precuneus | 2227.8 | 5962 | 15, -47, 33 | +6.16 | .0001 | [< .0001, .0002] |
| Positive | Right | precentral | 1479.5 | 3406 | 33, -18, 48 | +5.80 | .0001 | [< .0001, .0002] |
| Positive | Right | cuneus | 1403.2 | 1919 | 6, -81, 13 | +5.79 | .0001 | [< .0001, .0002] |
| Positive | Right | superiorparietal | 1138.9 | 2565 | 20, -42, 61 | +6.37 | .0001 | [< .0001, .0002] |
| Positive | Right | supramarginal | 768.4 | 1825 | 50, -30, 29 | +5.49 | .0001 | [< .0001, .0002] |
| Positive | Right | precuneus | 730.6 | 1566 | 5, -58, 22 | +6.44 | .0001 | [< .0001, .0002] |
| Positive | Right | superiortemporal | 678.3 | 1501 | 54, -5, -12 | +5.76 | .0001 | [< .0001, .0002] |
| Positive | Right | precentral | 376.6 | 914 | 55, -1, 9 | +5.62 | .0001 | [< .0001, .0002] |
| Positive | Right | postcentral | 331.2 | 982 | 48, -17, 20 | +4.84 | .0001 | [< .0001, .0002] |
| Positive | Right | medialorbitofrontal | 258.5 | 460 | 12, 37, -7 | +4.83 | .0001 | [< .0001, .0002] |
| Positive | Right | lingual | 218.1 | 248 | 13, -67, -4 | +4.47 | .0001 | [< .0001, .0002] |
| Positive | Right | precentral | 177.3 | 557 | 36, 7, 13 | +5.90 | .0005 | [.0002, .0008] |
| **One-sample (group mean) — Condition × correctness interaction (N = 89, df = 88)** | | | | | | | | |
| Negative | Left | lingual | 727.6 | 867 | -5, -82, 1 | -5.92 | .0001 | [< .0001, .0002] |
| Negative | Left | superiortemporal | 313.7 | 563 | -50, -11, -15 | -4.78 | .0001 | [< .0001, .0002] |
| Negative | Left | inferiorparietal | 214.8 | 462 | -38, -56, 18 | -4.00 | .0001 | [< .0001, .0002] |
| Negative | Left | middletemporal | 187.6 | 290 | -55, -60, 7 | -4.46 | .0003 | [.0001, .0005] |
| Negative | Left | inferiorparietal | 117.4 | 262 | -45, -64, 10 | -3.94 | .0093 | [.0081, .0105] |
| Negative | Left | cuneus | 111.0 | 123 | -4, -87, 14 | -3.87 | .0120 | [.0106, .0134] |
| Negative | Left | supramarginal | 100.3 | 222 | -58, -47, 28 | -3.66 | .0202ᵃ | [.0184, .0220] |
| Positive | Left | lateraloccipital | 748.4 | 989 | -28, -89, 7 | +5.21 | .0001 | [< .0001, .0002] |
| Positive | Left | rostralmiddlefrontal | 190.4 | 364 | -37, 23, 23 | +4.78 | .0006 | [.0003, .0009] |
| Positive | Left | lateralorbitofrontal | 119.9 | 309 | -27, 25, -8 | +4.45 | .0082 | [.0071, .0094] |
| Negative | Right | inferiorparietal | 1460.7 | 3279 | 45, -48, 20 | -5.24 | .0001 | [< .0001, .0002] |
| Negative | Right | cuneus | 609.7 | 777 | 5, -82, 12 | -5.81 | .0001 | [< .0001, .0002] |
| Negative | Right | superiorparietal | 316.2 | 442 | 20, -79, 34 | -4.57 | .0001 | [< .0001, .0002] |
| Negative | Right | precuneus | 293.6 | 856 | 14, -52, 37 | -4.89 | .0001 | [< .0001, .0002] |
| Negative | Right | posteriorcingulate | 145.0 | 345 | 6, -26, 40 | -4.70 | .0027 | [.0020, .0034] |
| Negative | Right | lingual | 137.2 | 165 | 7, -70, 2 | -4.19 | .0034 | [.0027, .0042] |
| Negative | Right | bankssts | 128.7 | 290 | 60, -34, 6 | -4.69 | .0048 | [.0039, .0057] |
| Negative | Right | superiortemporal | 118.7 | 197 | 51, -5, -15 | -5.01 | .0068 | [.0058, .0079] |
| Negative | Right | middletemporal | 112.8 | 268 | 49, -33, -8 | -4.71 | .0092 | [.0080, .0104] |
| Negative | Right | rostralmiddlefrontal | 100.3 | 201 | 27, 33, 34 | -3.71 | .0181ᵃ | [.0164, .0198] |
| Positive | Right | lateraloccipital | 976.9 | 1411 | 30, -89, 13 | +5.39 | .0001 | [< .0001, .0002] |
| Positive | Right | lingual | 865.8 | 1069 | 16, -79, -12 | +5.15 | .0001 | [< .0001, .0002] |
| Positive | Right | rostralmiddlefrontal | 283.4 | 545 | 40, 24, 22 | +4.53 | .0001 | [< .0001, .0002] |
| **Retrospective ELS, adjusted — Mindreading > control (N = 85, df = 79)** | | | | | | | | |
| Negative | Left | no suprathreshold cluster | — | — | — | — | — | — |
| Positive | Left | no suprathreshold cluster | — | — | — | — | — | — |
| Negative | Right | no suprathreshold cluster | — | — | — | — | — | — |
| Positive | Right | no suprathreshold cluster | — | — | — | — | — | — |
| **Retrospective ELS, adjusted — Correct > incorrect (N = 85, df = 79)** | | | | | | | | |
| Negative | Left | no suprathreshold cluster | — | — | — | — | — | — |
| Positive | Left | no suprathreshold cluster | — | — | — | — | — | — |
| Negative | Right | no suprathreshold cluster | — | — | — | — | — | — |
| Positive | Right | no suprathreshold cluster | — | — | — | — | — | — |
| **Retrospective ELS, adjusted — Condition × correctness interaction (N = 85, df = 79)** | | | | | | | | |
| Negative | Left | no suprathreshold cluster | — | — | — | — | — | — |
| Positive | Left | no suprathreshold cluster | — | — | — | — | — | — |
| Negative | Right | no suprathreshold cluster | — | — | — | — | — | — |
| Positive | Right | no suprathreshold cluster | — | — | — | — | — | — |
| **Retrospective ELS, unadjusted — Mindreading > control (N = 85, df = 83)** | | | | | | | | |
| Negative | Left | middletemporal | 104.6 | 216 | -51, -36, -9 | -4.63 | .0166ᵃ | [.0150, .0182] |
| Positive | Left | no suprathreshold cluster | — | — | — | — | — | — |
| Negative | Right | no suprathreshold cluster | — | — | — | — | — | — |
| Positive | Right | no suprathreshold cluster | — | — | — | — | — | — |
| **Retrospective ELS, unadjusted — Correct > incorrect (N = 85, df = 83)** | | | | | | | | |
| Negative | Left | no suprathreshold cluster | — | — | — | — | — | — |
| Positive | Left | no suprathreshold cluster | — | — | — | — | — | — |
| Negative | Right | no suprathreshold cluster | — | — | — | — | — | — |
| Positive | Right | no suprathreshold cluster | — | — | — | — | — | — |
| **Retrospective ELS, unadjusted — Condition × correctness interaction (N = 85, df = 83)** | | | | | | | | |
| Negative | Left | no suprathreshold cluster | — | — | — | — | — | — |
| Positive | Left | no suprathreshold cluster | — | — | — | — | — | — |
| Negative | Right | no suprathreshold cluster | — | — | — | — | — | — |
| Positive | Right | no suprathreshold cluster | — | — | — | — | — | — |
| **Prospective ELS, adjusted — Mindreading > control (N = 89, df = 83)** | | | | | | | | |
| Negative | Left | no suprathreshold cluster | — | — | — | — | — | — |
| Positive | Left | no suprathreshold cluster | — | — | — | — | — | — |
| Negative | Right | no suprathreshold cluster | — | — | — | — | — | — |
| Positive | Right | no suprathreshold cluster | — | — | — | — | — | — |
| **Prospective ELS, adjusted — Correct > incorrect (N = 89, df = 83)** | | | | | | | | |
| Negative | Left | no suprathreshold cluster | — | — | — | — | — | — |
| Positive | Left | no suprathreshold cluster | — | — | — | — | — | — |
| Negative | Right | no suprathreshold cluster | — | — | — | — | — | — |
| Positive | Right | no suprathreshold cluster | — | — | — | — | — | — |
| **Prospective ELS, adjusted — Condition × correctness interaction (N = 89, df = 83)** | | | | | | | | |
| Negative | Left | no suprathreshold cluster | — | — | — | — | — | — |
| Positive | Left | no suprathreshold cluster | — | — | — | — | — | — |
| Negative | Right | no suprathreshold cluster | — | — | — | — | — | — |
| Positive | Right | no suprathreshold cluster | — | — | — | — | — | — |
| **Prospective ELS, unadjusted — Mindreading > control (N = 89, df = 87)** | | | | | | | | |
| Negative | Left | no suprathreshold cluster | — | — | — | — | — | — |
| Positive | Left | no suprathreshold cluster | — | — | — | — | — | — |
| Negative | Right | no suprathreshold cluster | — | — | — | — | — | — |
| Positive | Right | no suprathreshold cluster | — | — | — | — | — | — |
| **Prospective ELS, unadjusted — Correct > incorrect (N = 89, df = 87)** | | | | | | | | |
| Negative | Left | no suprathreshold cluster | — | — | — | — | — | — |
| Positive | Left | no suprathreshold cluster | — | — | — | — | — | — |
| Negative | Right | no suprathreshold cluster | — | — | — | — | — | — |
| Positive | Right | no suprathreshold cluster | — | — | — | — | — | — |
| **Prospective ELS, unadjusted — Condition × correctness interaction (N = 89, df = 87)** | | | | | | | | |
| Negative | Left | no suprathreshold cluster | — | — | — | — | — | — |
| Positive | Left | no suprathreshold cluster | — | — | — | — | — | — |
| Negative | Right | no suprathreshold cluster | — | — | — | — | — | — |
| Positive | Right | no suprathreshold cluster | — | — | — | — | — | — |

Note. Vertex-wise cluster-forming threshold p < .001; cluster-wise family-wise error corrected using FreeSurfer's precomputed Monte Carlo simulations, applied separately within each hemisphere and each direction. Direction refers to the sign of the group-mean effect in the one-sample model and to the sign of the early life stress slope in the between-subjects models. Peak coordinates are in MNI305 space (fsaverage). Anatomical labels denote the Desikan–Killiany parcel containing the Peak t is the t-statistic at the cluster's peak vertex; degrees of freedom differ across models and are given in each section heading. extend across parcel boundaries. Em dashes denote contrasts with no suprathreshold clusters.

ᵃ Cluster-wise p exceeds 0.0125, the threshold corresponding to family-wise error of .05 across both hemispheres and both directions; reported as exploratory.
