## Supplementary figures and images for "The Influence of Early Life Stress on the Development of Neural Representations during Mentalizing"

### Supplemetary Figure S1

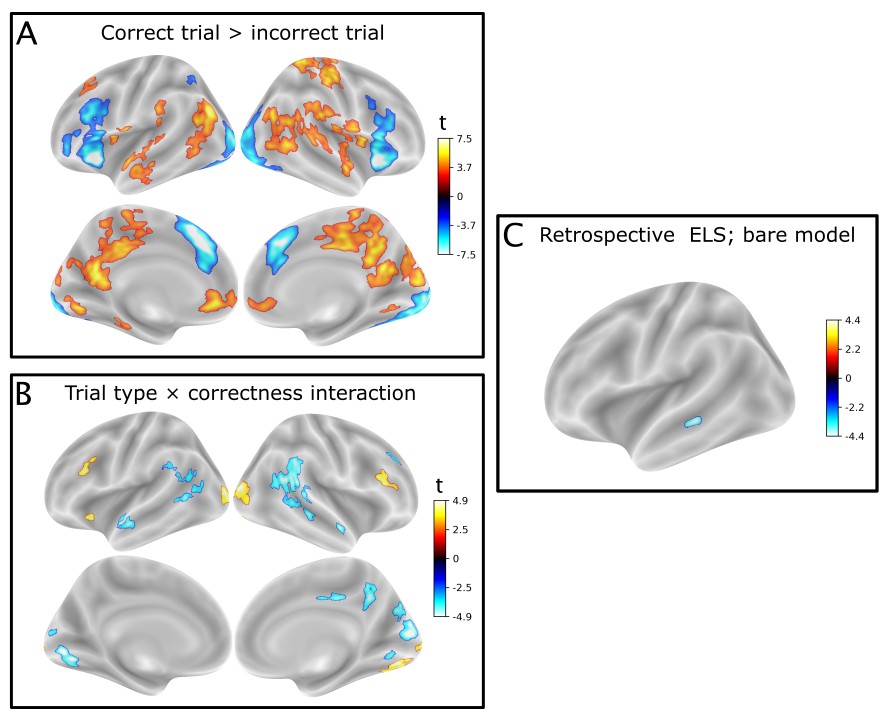
